# Propionate reinforces epithelial identity and reduces aggressiveness of non-small cell lung carcinoma via chromatin remodelling

**DOI:** 10.1101/2023.01.19.524677

**Authors:** Vignesh Ramesh, Paradesi Naidu Gollavilli, Luisa Pinna, Mohammad Aarif Siddiqui, Simon Toftholm Jakobsen, Elisa Le Boiteux, Karen Ege Olsen, Ole Nørregaard Jensen, Rasmus Siersbæk, Kumar Somyajit, Paolo Ceppi

## Abstract

Epithelial to mesenchymal transition (EMT) is a developmental cellular program driving metastasis and chemo-resistance in cancer, but its pharmacological treatment has been so far challenging. Targeting deregulated metabolic processes in cancer is emerging as a realistic therapeutic strategy. Here, we used an EMT-focussed integrative functional genomic approach and identified negative association of the short-chain fatty acids propionate and butanoate with EMT in non-small cell lung cancer (NSCLC) patients. Strikingly, in vitro treatment of lung cancer cell lines with propionate reinforced the epithelial transcriptional program promoting cell adhesion and reverting the aggressive and chemoresistant EMT phenotype. Propionate treatment reduced cells metastatic ability in nude mice and limited lymph nodal spread in a genetic NSCLC mouse model. Further analyses indicated chromatin remodeling via H3K27 acetylation (p300-mediated) as the mechanism shifting the EMT balance towards epithelial state upon propionate. Propionate administration could be tested in the clinic for reducing NSCLC aggressiveness.

**Highlights:** An EMT-centric investigation of metabolic processes in a comprehensive lung cancer transcriptome profiles identified negative associations between EMT and SCFAs (propionate and butyrate)

Propionate enhances the epithelial features both at the molecular and cellular levels

Pre-treatment of cells with propionate inhibits EMT associated processes including migration and sensitizes the cells to chemotherapeutic drug cisplatin

Oral administration of propionate inhibits EMT-mediated lung colonization ability of NSCLC cells, and lymph node metastasis in a genetic mouse NSCLC model

Molecular mechanistic investigation of propionate revealed chromatin remodelling through p300-mediated histone acetylation in E-cadherin gene regulation along with epithelial features reinforcement

## Introduction

Lung cancer is the second most frequently diagnosed cancer and a leading cause of cancer deaths with non-small-cell lung cancer (NSCLC) histological subtype accounts to about 85% of lung cancer [1,2]. Development of next generation molecular targeted therapeutics and immunotherapy for lung cancer such as with osimertinib and permbrolizumab, respectively, have shown improvements in recent years in the treatment of advanced or metastatic lung cancer patients [3,4]. However, the NSCLC still present a poor prognosis rate with the 5-year overall survival of around 25% [5].

Epithelial-to-mesenchymal transition (EMT) is a developmental phenotypic plasticity program transforming from an epithelial-like phenotype to loosely attached mesenchymal-like phenotype. In cancer, EMT confers more aggressive features of tumorigenesis including increased migratory and invasive ability, immune suppression, chemoresistance and metastatic colonization with poor patient survival [6–9]. The importance of EMT as a key early step in tumorigenic events in lungs has also been highlighted [10,11]. Several factors most importantly the transcription factors like ZEB1, SNAI1, SNAI2 and TWIST have been deeply studied for their role in the induction of EMT [12]. However, the pharmacological targets or inhibition of these transcription factors are still challenging [13]. Apart from the transcription factors, EMT in cancer is induced or modulated by microRNAs [14], epigenetic alterations [15] or as an interplay of these factors [16]. In recent years, metabolic reprogramming has been shown to function as a crucial hallmark feature of EMT regulation in various cancer types. For instance, EMT is induced with elevated levels of polyol pathway in the conversion of glucose to fructose in lung cancer [17], by branched chain amino acid metabolic deregulation [18] or induced by 2-hydroxyglutarate, oncometabolite, in colorectal cancer [19]. Numerous metabolic pathway inhibitors targeting EMT have been identified with clinical benefits in many cancer types [20]. However, several metabolic inhibitors still face the limitations of specificity, toxicity issues and patient stratification for improved responsiveness [21–23]. This is primarily due to the limited or lack of understanding of the complex metabolic changes during the advanced stages of cancer progression [21]. This emphasis the need for a comprehensive characterization of cellular, molecular, and functional level processes associated with the metabolic impairment in EMT condition for therapeutic interventions.

As EMT in cancer is a very dynamic process with varied spectrum of phenotypic states, assessing EMT in the patients-derived samples is still technically challenging [24], and deeper exploration of the interplay between tumor and systemic metabolism would be highly useful in targeting the metabolic control of aggressive tumors [25]. With this in mind, we have performed a global comprehensive transcriptome analysis of lung cancer gene expression profiles in the context of EMT associated metabolic processes using gene-sets to identify the metabolic pathways that could inhibit or reverse the EMT condition. From the analysis, we identified and deeply explored the role of propionate’s mechanism of action in inhibiting EMT-mediated aggressive features including metastasis, and also inducing chemosensitization of the cells with a specific role of chromatin re-modelling in enhancing the lung epithelial aspects.

## Methods

### Cell culture

Lung cancer cell lines were purchased from ATCC and KPL cell line was derived from lung tumorigenic mice intubated with AAV-KPL virus. A549 and KPL cell lines were cultured in DMEM high glucose medium (Gibco), SKMES1 and NCI-H520 were cultured in RPMI-1640 medium (Gibco), and NCI-H23 was cultured in RPMI-1640 with 1 mM sodium pyruvate (Sigma). All media were supplemented with 2 mM of L-glutamine (Gibco), 100 U/ml of Penicillin-Streptomycin (Gibco), and 10% Fetal bovine serum (Gibco). Cell lines were maintained at 37° C with 5% CO_2_ level in a humidified incubator.

### Drug Treatment

For drug treatment studies, around 1.5 × 10^5^ cells were seeded for the treatment in a 6-well cell culture dish. After 24 hours of seeding, cells were treated with the appropriate inhibitors in the presence or absence of sodium proprionate (Sigma, P1880). Inhibitors used in the study are Recombinant Human TGF-β1 protein (R&D Systems, 240-B-002), SAHA (Sigma, SML0061), HAT inhibitor VII, CTK7A (Sigma, 382115), A-485 (MedChemExpress, HY-107455), AZD3965 (MedChemExpress, HY-12750), Syrosingopine (Sigma, SML1908), AR420626 (Sigma, SML1339), mocetinostat (MedChemExpress, HY-12164), MI192 (Sigma, SML1451), GW9662 (Sigma, M6191), Troglitazone (Sigma, T2573), 5-Aza-2’-deoxycytidine (Sigma, A3656), GSK126 (MedChemExpress, HY-13470), and GSK343 (MedChemExpress, HY-13500). Sodium butyrate (303410) and sodium acetate (S2889) were purchased from Sigma.

### EDTA Detachment

For experiments involving EDTA mediated detachment, cells were harvested by washing twice with 1X PBS without Ca^2+^/Mg^2+^ (Lonza), incubated with 5 ml of 0.5 mM EDTA (Lonza) at 37° C for 2 minutes to detach the cells, mixed well to obtain single cell suspension with equal volume of ice-cold 1X PBS containing Ca^2+^/Mg^2+^ (Sigma), and centrifuged at 800 rpm for 3 minutes at 4° C. Cells were then mixed with 5 ml of warm media and incubated at 37° C for 1 hour with frequent agitation to re-express E-cadherin. After 1 hour of incubation, cells were then washed twice with 1X PBS containing Ca^2+^/Mg^2+^ and centrifuged at 800 rpm for 4 minutes to proceed for the downstream experimental purposes.

### Lung experimental metastasis model

NSG mice strain was purchased from Jackson Laboratory for the experimental lung metastasis study. Around 1 × 10^6^ cells of A549-PFUL2G or SKMES1-PFUL2G were seeded in a 10-cm dish and after 24 hours of seeding, cells were treated with sodium propionate (5 mM) for 3 days. The cells were then harvested with EDTA detachment method. Around 500,000 cells were resuspended in 100 μl of 1X PBS containing Ca^2+^/Mg^2+^ and injected in the tail vein of NSG mice using 27G needle. Lung colonization was examined using in vivo imaging system (IVIS, Perkin Elmer) in the anaesthetized mice by intraperitoneal injection of 3 mg/ml of D-luciferin (Cayman Chemicals) as substrate for luciferase enzyme produced by PFUL2G cells in the lungs, and bioluminescence signal was recorded as radiance (p/sec/cm^2^/sr) and reported the overall bioluminescence signal as total flux (p/s). Significance was calculated using the unpaired t-test between the groups.

### CRISPR/Cas9-mediated lung tumorigenesis in mice and SP administration in drinking water

Ultra-purified recombinant adeno-associated virus (AAV-KPL) was produced and obtained from VectorBuilder Inc., USA from the AAV:ITR-U6-sgRNA(Kras)-U6-sgRNA(p53)-U6-sgRNA(Lkb1)-pEFS-Rluc-2A-Cre-shortPA-KrasG12D_HDRdonor-ITR vector (addgene, #60224). Adult in-house bred C57BL/6 strain constitutively expressing Cas9 (B6J.129(B6N)-*Gt(ROSA)26Sor^tm1(CAG-cas9*,-EGFP)Fezh^*/J) mice were randomized based on gender and age. Pre-administration of sodium chloride (SC, 150 mM) or sodium propionate (SP, 150 mM) in mice drinking water was started 1 week before lung delivery of the virus and continued until the end of the study. Drinking water was changed twice a week with freshly prepared compounds. AAV-KPL virus was oropharyngeally delivered to mice resulting in a B/6.*Kras^G12D/G12D^p53^Δ/Δ^Lkb1^Δ/Δ^* genotype (referred to as KPL mice). For oropharyngeal virus delivery, mice were anaesthetized using isoflurane, and 3 × 10^11^ viral units per mouse were administered dissolved in 25 μl of 0.9% sodium chloride. Mice were sacrificed at humane endpoint, scoring breathing patterns and body weight loss. Mice were euthanized by cervical dislocation to collect lungs and lymph nodes for H&E staining, and mice survival was recorded. Presence of lung tumors and lymph node metastases was evaluated by a trained pathologist at the Odense University hospital (KEO). Animal protocols were approved by the Danish Animal Welfare Authority (approval 2020-15-0201-00607).

### Flow cytometry

For the Flow cytometry analysis, A549 cells treated with SP (5 mM) for 3 days in 10-cm dish were detached using EDTA detachment protocol and around 5 × 10^5^ cells were washed once with 3 ml of blocking buffer (2% BSA (Roth, 8076.4) in 1X PBS containing Ca^2+^/Mg^2+^), and centrifuged the cells at 800 rpm for 3 minutes. Cells were then stained with 2.5 μl of PE anti-human E-cadherin (50 μg/ml; Biolegend, 324106) or matched concentration of PE Mouse IgG1, κ Isotype ctrl (FC) (200 μg/ml; Biolegend, 400113) as per manufacturer’s protocol (Biolegend) for 1 hour in dark at room temperature with frequent agitation. Cells were then washed once with blocking buffer and resuspended in 500 μl FACS resuspension buffer (2% FBS with 5 mM EDTA in 1X PBS) containing 5 μl of DAPI (20 μg/ml), and were immediately analyzed using CytoFLEX (Beckman Coulter) and the analysis was performed using FlowJo software v10.6.

### Immunofluorescence

Around 1 × 10^5^ cells were seeded on a glass coverslip for 3 days SP treatment in a 12-well cell culture plate. After 3 days SP treatment, cells were washed with 1X PBS followed by the addition of ice-cold methanol (Sigma), and incubated the coverslips for 20 minutes at room temperature. Cells were then washed with 1X PBS and blocked with blocking buffer (3% BSA in 1X PBS) at room temperature for 1 hour and washed again with 1X PBS. Cells were incubated overnight at 4° C with the primary antibodies. Primary antibodies used in the study were: E-cadherin (1:250, mouse, Cell Signaling, 14472), EPCAM (1:100, rabbit, Invitrogen, PA5-29634) and ZO-1 (1:150, goat, abcam, ab190085) prepared in the blocking buffer. Next day, coverslips were washed thrice with 1X PBS and incubated with the corresponding fluorochrome conjugated secondary antibodies for 1 hour in dark at room temperature. Secondary antibodies used in the study were: anti-mouse Alexa Fluor 488 conjugate (1:250, ThermoFisher Scientific, A28175), anti-rabbit Alexa Fluor 488 conjugate (1:200, Cell Signaling, 4412S) and anti-goat Alexa Fluor 488 conjugate (1:200, abcam, ab150129) prepared in the blocking buffer. After secondary antibody incubation, cells were washed with 1X PBS, and the coverslips mounted on a glass slide using FluoroSheild with DAPI containing mounting medium (VWR, F6057). Cells were then visualized using Leica DM5500B fluorescence microscope or with Nikon Widefield Ti-2, and the images were acquired either using Leica Application Suita-X software or with NIS-Elements Viewer 5.21.

### Western blot

Cells were harvested for protein using Pierce RIPA lysis buffer (Thermo Scientific) containing 1X Halt Protease & Phosphatase inhibitor cocktail (Thermo Scientific) and estimated the proteins using Pierce BCA Protein Assay Kit as per manufacturer’s protocol (ThermoFisher Scientific). 15 – 30 μg of proteins were resolved in 8% SDS-PAGE separating gel and proteins were then allowed to transfer from the resolved gel to a PVDF membrane (Thermo Scientific). Membranes were blocked with the blocking buffer (5% non-fat dried milk powder or 3% BSA prepared in 1X TBS-T) for 1 hour at room temperature, and then incubated overnight at 4° C with the primary antibodies. Primary antibodies with the dilutions used in the study are: E-cadherin (1:5000, mouse, Cell Signaling, 14472), ZEB1 (1:2000, rabbit, Sigma, HPA027524), γH2AX (1:2000, rabbit, Cell Signaling, 9718S), GRHL1 (1:1000, rabbit, abcam, ab111582), OVOL2 (1:1000, mouse, abcam, ab169469), H3K27ac (1:3000, rabbit, abcam, ab177178), H3K27me3 (1:3000, mouse, abcam, ab6002), H3 (1:10000, rabbit, Cell Signaling, 9715S), Acetyl-CBP-Lys1535/p300-Lys1499 (1:1000, rabbit, Cell Signaling, 4771S), TUBA4A (1:10000, mouse, Sigma, T6199) and β-Actin HRP conjugated (1:10000, Cell Signaling, 12262). Next day, the membranes were washed thrice with 1X TBS-T and incubated the membrane blots in secondary antibody dilution (Southern Biotech). Secondary antibodies conjugated with HRP used in the study are: Goat Anti-mouse IgG1-HRP (1:10000, Southern Biotech, 1071-05), Goat Anti-rabbit IgG-HRP (1:10000, Southern Biotech, 4030-05) and Goat Anti-mouse IgG3-HRP (1:10000, Southern Biotech, 1101-05). Protein bands were detected using Pierce ECL western blotting solution (Thermo Scientific) with the development of X-ray films (Thermo Scientific).

### Cell proliferation assay and dose response curve analysis

Approximately 1000 cells per well were seeded in a low density (5-10% confluence) in a 96-well plate in triplicates, and after 24 hours, cells were treated with sodium propionate at 5 mM. Plates were then incubated at 37° C in IncuCyte S3 (Sartorius) and real-time live-cell proliferation analysis was performed at regular intervals of 4 hours for 12 days with phase contrast image acquisition mode using Incucyte S3 software. Proliferation was plotted as percent confluence over time.

For dose response curve analysis with cisplatin, 1000 cells/well were seeded and pre-treated with SCFAs (SP or SB) for 48 hours followed by cisplatin (Tocris, 2251) treatment in a dose dependent manner for 72 hours. Plates were incubated at 37° C in IncuCyte S3, and real-time live-cell proliferation analysis was performed at regular intervals of 4 hours for 3 days with phase contrast image acquisition mode to quantify growth using Incucyte S3 software. To quantify cell death, Cytotox Green (Sartorius) was mixed in the media and the image acquisition mode was set to green channel. Dose responsive curve was generated using a non-linear fit of log(inhibitor) vs normalized response using Graphpad with a Two-way ANOVA analysis to obtain the significance between the conditions.

### Migration assay

A549 cells were seeded in 96-well plate of around 3000 cells per well and pre-treated with sodium propionate (5 mM) for 72 hours. Once the cells reach confluency, scratch was made using Incucyte 96-well Woundmaker Tool (Essen BioScience), washed the cells with 1X PBS to remove the debris, and incubated the plates at 37° C in IncuCyte S3. Relative wound density was measured using the integrated quantification module for scratch wound at regular intervals of 4 hours for 3 days in Incucyte S3.

### ECM cell adhesion assay

ECM cell adhesion assay was performed as per manufacturer’s instructions (Merck). Briefly, 70,000 cells per well were seeded in the wells of the strips coated with 7 different ECM proteins. Cells were allowed to attach to the surface for 1 hour at 37° C incubator, and then gently washed twice with the provided assay buffer. After washing, 100 μl of the cell stain solution was added to each well, and allowed to incubate for 5 minutes followed by cell stain removal by washing the wells thrice with deionized water. Wells were then air dried, added with 100 μl of extraction buffer, and incubated on a rotating shaker until the stain was completely solubilized. The absorbance was then measured at 540 nm in a microplate reader.

### Quantitative image-based cytometry (QIBC)

A549 cells of around 2.5 × 10^5^ cells per well were seeded on a round glass coverslip in a 6-well plate. After 24 hours of seeding, cells were pre-treated with sodium propionate at 5 mM for 24 hours followed by the treatment with cisplatin in a dose dependent concentration for around 6 hours. Cells were then added with 10 μM of EdU in each well and the plates were incubated for last 20 minutes before fixation. Cells were fixed with 4% formaldehyde stabilized with 0.5-1.5% methanol (VWR, 9713) for 12 minutes at room temperature, washed with ice-cold 1X PBS, and further incubated with 1X PBS containing 0.2% Triton X-100 (Sigma, X100) for 5 minutes. In case of assessing chromatin-based proteins for QIBC, cells were pre-extracted for the chromatin-bound proteins by washing the treated cells with ice-cold 1X PBS followed by incubating the cells with ice-cold 1X PBS containing 0.2% Triton X-100 for 90 seconds. Cells were then washed again with ice-cold 1X PBS and fixed the cells with 4% formaldehyde for 12 minutes at room temperature. After fixation, Click-iT Edu reaction was allowed to perform using A647-Azide (ThermoFisher Scientific, A10277) through a copper-catalyzed click chemistry for 30 minutes in dark and washed once with 1X PBS containing 0.1% Tween20 (Sigma, P1379), and thrice with 1X PBS. Coverslips were then incubated with primary antibodies for 1 hour in dark at room temperature and washed with 1X PBS containing 0.1% Tween20. Primary antibodies for QIBC used were: H3K27ac (1:500 rabbit, abcam, ab177178), H3K4me3 (1:1000, rabbit, abcam, ab8580), HP1 (1:500, mouse, Santa Cruz, SC515341), H3S10ph (1:2000, mouse, abcam, ab14955), γH2AX (1:2000, mouse, Biolegend, 613402), RAD51 (1:1000, mouse, abcam, ab213), TP53BP1 (1:2000, mouse, Millipore, Mab3802). Next, cells were incubated with the secondary antibodies containing DAPI (0.5 μg/ml) as nuclear counterstain for 30 minutes in dark at room temperature. Secondary antibody conjugates of Alexa Fluor 488 or 568 for mouse (Cat. No. A-11029 or A-11031) and rabbit (Cat. No. A-11034 or A-11036) from ThermoFisher Scientific in 1:2000 dilutions were used. Coverslips were then washed once with 1X PBS containing 0.1% Tween20, and thrice with 1X PBS. Coverslips were then rinsed with MilliQ water, air dried and mounted onto the glass slide using Mowiol 4-88 mounting medium. Approximately 40-80 images were acquired amounting atleast 1000 cells to 10000 cells per sample condition using ScanR inverted microscope High-content Screening Station (Olympus) with 20X optics with in-built ScanR acquisition software as described previously [26]. The acquired imaging parameters were then exported to TIBCO software to analyze and report the differences between the conditions at the population level.

### siRNA Transfection

Reverse transfection of propionate metabolism-specific siRNAs was performed using Lipofectamine RNAiMAX transfection reagent as per manufacturer’s protocol (ThermoFisher Scientific). Briefly, transfection complex was prepared by mixing 50 nM of SMARTPool ON-TARGETplus Human siRNAs (horizon, PerkinElmer) with 3 μl of Lipofectamine RNAiMAX transfection reagent (ThermoFisher Scientific) in 200 μl Opti-MEM (ThermoFisher Scientific) and incubated the complex for 15 minutes at room temperature. 200 μl of transfection complex was then mixed with 800 μl of A549 cells (3×10^5^ cells) while seeding of the cells in a 24-well plate. After 48 hours of siRNA transfection, cells were treated with sodium propionate (5 mM) for 24 hours, and the cells were harvested for western blot analysis of EMT markers.

### RNA Isolation, cDNA synthesis and Real-time PCR

A549 cells were harvested using 700 μl of QIAzol reagent (Qiagen). Total RNA was isolated as per the manufacturer’s protocol using miRNeasy kit (Qiagen) and eluted the RNA in 40 μl of Nuclease free water (VWR). 500 ng of isolated RNA was converted to cDNA using Tetro cDNA synthesis kit (meridian BioScience) with random hexamers using the routine protocol. Real-Time PCR was performed using 2X Taqman Universal master mix II, no UNG buffer (Applied Biosystems). Briefly, 5 μl of synthesized cDNA was mixed with 1X Universal Taqman master mix buffer along with 1X Taqman probes and the reaction was set in Roche 96-well system with pre-incubation at 95° C for 10 mins followed by 40 cycles of 95° C for 15s and 60° C for 1 minute in a FAM acquisition mode. Real-time qPCR analysis was performed with Ct values obtained from LightCycler 96 SW 1.1 (Roche) and determined the folds as test over control using the ΔΔCt method.

### RNA-Sequencing

Total RNA was isolated from A549 cells treated with sodium propionate for 3 hours, 24 hours, 3 days and 12 days. 1 μg of RNA was diluted in 25 μl of Nuclease free water. Sequencing libraries were constructed using NEBNext Ultra RNA Library Prep Kit for Illumina according to the manufacturer’s protocol (NEB) and paired-end sequencing was performed with NovaSeq 6000 platform (Illumina).

RNA-Seq analysis was performed by subjecting the Fastq files for the QC analysis using FastQC followed by aligning the reads to the reference genome GRCh38 release 105 with the respective human gene annotation file using STAR aligner v2.7.9a [27]. The aligned files were then used for counting the raw reads using the Featurecounts function in Rsubread v2.10.5 package [28] in R 4.2.1. Raw count reads were then used for the differential gene expression analysis with DESeq2 v1.36.0 [29] package in R with an adjusted p-value < 0.0001 and fold change of 2. Mean normalized values from DESeq2 were used for the expression analysis of gene-set between conditions.

### scRNA-Seq

A549 cells for single cell RNA sequencing (scRNA-Seq) was performed using 10X Genomics guidelines at the Institute of Human Genetics, Friedrich-Alexander-University of Erlangen-Nürnberg, Erlangen, Germany. Around 6000 cells were read with approximately 25,000 reads per cell. Briefly, the filtered_feature_bc_matrix data containing barcodes, features and matrix obtained from cellranger-4.0.0 pipeline were further analyzed by Seurat v4.2.1 package in R. Quality control followed by pre-processing of the file was performed with the filtering of the data by removing cells with less than 500 genes, UMIs with <2500 & >45000 total number of molecules, removed more than 10% mitochondrial genes and > 5% largest gene. Log_10_ genes per UMI was set at > 0.85 for filtering. Counts were then normalized for the library size, identified 2000 variable genes to perform PCA. Variations including mitochondrial genes, cell cycle genes, and genes & UMI counts per cell were all regressed out for the downstream analysis. First 25 principal components from PCA of the data were selected to determine the cell clusters using Lovain algorithm with a resolution of 0.4 and visualized the cell clusters using tSNE dimentionality reduction method. Marker genes were then identified for each cluster. AddModuleScore function in Seurat was applied to identify the gene-set activity metrics in each cell clusters for epithelial, mesenchymal and SP regulated gene-sets.

### Gene Enrichment analysis

Gene set enrichment analysis (GSEA) was performed for the patient samples categorized as low and high based on the median of the z-score activation for SP-regulated genes with the gene ranks calculated using Signal2Noise metric. For the association of SP-regulated gene-sets with the Hallmark Adherens Junction or Hallmark Adherens Surface, continuous label of SP gene-set z-score activation in the expression profile using the gene ranks calculated using Pearson ranked gene metric was employed. Significance was set to nominal p-value<0.05 and FDR<0.25. Gene enrichment analysis using representational overlap analysis was carried out using Enrichr [30] for SP-regulated genes identified from RNA-Seq analysis or for the cell cluster marker genes identified from scRNA-seq of A549 cell lines. L1000CDS^2^ was used to identify the small molecule mimics of SP-regulated gene signature identified from RNA-Seq [31].

### ChIP-Seq

H3K27ac ChIP-seq was performed on SP treated A549 cells as described previously [32]. In brief, cells were double crosslinked in 2 mM DSG and 1% FA before harvesting of the cells by scraping from the culture dish. Cell pellets were washed and sonicated until fragment size ranges between 200bp and 500bp. Sonicated chromatin was then immunoprecipitated overnight using Dynabeads protein A (Dynabeads, 10002D) coated with the antibody of interest. Next day, the beads were washed six times in cold RIPA buffer, and the DNA was decrosslinked and purified using standard phenol-chloroform purification. Purified DNA was submitted to NGS library preparation using NEBNext Ultra II DNA Library Prep Kit (New England BioLabs, E765) and sequenced with NovaSeq 6000 (Illumina) platform to reach approximately 25 million paired-end reads per sample. Paired-end reads were aligned to the human genome using Hisat2 v2.1.0. Duplicated reads from PCR amplification were removed using samtools [33] before peaks were called with the default setting in MACS2 [34]. Only peaks called within all replicates were considered as consensus peaks. Homer [35] was used to visualize heatmaps and for counting normalised tag counts within promoter regions.

### SingScore

RNA-Seq normalised counts from DESeq2 were ranked for each condition and SingScore was used to calculate gene set enrichment scores as described previously [36]. The calculated EMT score represents the difference between the mesenchymal and the epithelial gene-sets obtained from previous report [6].

### Histone extraction and digestion

Histones were extracted from frozen cell pellets by acid extraction. Briefly, nuclei were isolated using nuclear isolation buffer (15 mM Tris-HCl pH 7.5, 60 mM KCl, 11 mM CaCl_2_, 5 mM NaCl, 5 mM MgCl_2_, 250 mM sucrose, 1 mM dithiothreitol, 10 mM sodium butyrate and 0.1% Igepal) supplemented with protease (cOmplete™ Protease Inhibitor Cocktail, Roche) and phosphatase (PhosSTOP, Roche) inhibitors. Nuclei were pelleted by centrifugation (1,000g – 5 min) and washed twice with nucleus isolation buffer without Igepal. Histones were extracted by resuspending the pellet in 0.2N H_2_SO_4_ for 1 hour, and the supernatant was collected after centrifugation (20,000g – 5 min). Histones were precipitated overnight after adding trichloroacetic acid to a final concentration of 20%. Histones were pelleted by centrifugation (20,000g – 15 min) and washed once with 0.1% HCl in acetone and then twice with pure acetone. After the last wash, histones were air-dried and resuspended in H_2_O. All steps were performed at 4°C. Purity of histones was evaluated by SDS-PAGE and protein concentration was determined using Qubit™ Protein Assay Kit (Invitrogen). Histones were digested using the Propionylation-PIC method (Maile *et al*., 2015). For each reaction, 5μg of histones were diluted in H_2_O to a total volume of 9μL. pH was adjusted with 0.4μL of 5% NaOH and buffered with 1μL of 1M EPPS. Before propionylation, cysteine residues were reduced and alkylated with 12.5 mM dithiothreitol and 25 mM iodoacetamide, respectively. Propionylation reaction was performed by adding 1.5μL of 1% propionic anhydride in acetonitrile for 2 min at RT and was quenched with 1.5μL hydroxylamine 80mM. Histones were digested overnight at 37°C using 0.1μg of trypsin. For derivatization, peptides were incubated at 37°C one additional hour with 4.5μL of 1% phenylisocyanate in acetonitrile. Histone peptides were desalted using homemade C18-stage-tips.

### LC-MS/MS analysis

Stage-tip desalted histone peptides were resuspended in HPLC solvent A (0.1% formic acid in H_2_O). LC-MS/MS analysis was performed with ~500ng peptides using a nano-flow HPLC system (EASY-nLC 1000, ThermoScientific) coupled with an Orbitrap mass spectrometer (Exploris480, ThermoScientific). Peptides were loaded onto a ~4cm long, 100μm ID precolumn packed with 5μm C18 particles and separated onto a 18cm long, 75μm ID analytical column packed with 3μm C18 particles. Peptides were separated at a flow rate of 250nL/min by a linear gradient from 2% solvent B (0.1% formic acid in 95% acetonitrile) to 45% solvent B over 40 min followed by a ramp to 100% solvent B in 3 min and stabilization at 100% solvent B during 7 min (total run time: 50 min). Full mass range spectra were acquired at a resolution of 120,000 (at *m/z* 200) with a scan range of *m/z* 300-2000, and the 15 most intense precursors were selected for MS/MS. Fragmentation was performed by high energy collisional dissociation (HCD) with 30% normalized collision energy. MS/MS spectra were acquired at a resolution of 30,000 (at m/z 200).

### Epiproteomic data analysis

Thermo .raw files were processed with Proteome Discoverer 2.5 (ThermoScientific) using MASCOT and Percolator with label-free quantification node. Spectra were searched with MASCOT against a human histone database (Swiss-Prot reviewed, downloaded from www.uniprot.org), using ArgC as digestion enzyme with 1 missed cleavage accepted. Full MS and MS/MS tolerances were set to 5 ppm and 0.05 Da, respectively. Propionylation of lysines and alkylation of cysteines with iodoacetamide were mentioned as static modifications. Several dynamic modifications were considered for the search: propionylation of protein N-terminus, derivatization of peptide N-terminus by phenylisocyanate, acetylation of protein N-terminus, methylation/dimethylation/trimethylation of lysines and phosphorylation of serines and threonines. The identity of each peak was verified by hand using retention times and MS/MS spectra. The area under the curve (AUC) was calculated by the software for each specific proteoform and was normalized to the global amount of peptides with the same primary sequence. To identify changes in histone PTM abundances, log_2_ fold-change values of SP-treated cells compared to control were calculated for each point.

### Gene-sets source

Metabolic process associated gene-sets, Hallmark gene-sets, and EMT associated gene-sets were collected from MSigDb v6.2. In total 335 metabolic process associated gene-sets were collected, 135 KEGG and 200 REACTOME gene-sets. Metabolic process associated gene-sets collectively represent diverse metabolic processes from KEGG or REACTOME representing carbohydrates, steroids, amino acids, vitamins, lipids, fatty acids, catabolic and anabolic, enzymatic activities including coenzymes or co-factors, hormones, nucleotide, biopolymer or macromolecular, alcohol, amine, drug, organic, inorganic and acyl chain related metabolisms. It also includes EMT associated gene-sets as a positive control for the pathway activation study.

### Metabolic process activation analysis

Metabolic process activity score was calculated as described previously [37]. Fold expression values (log_2_) for each gene in the tumor samples was calculated relative to the median expression value across the tumor samples as reference. Next, mean and standard deviation of the fold expression in the whole gene expression profile for each tumor sample was calculated. Similarly, mean was calculated from the fold expression value for the metabolic gene-sets for each tumor samples by extracting the fold expression values for all genes in the gene-set. The metabolic process activity score (z-score) was calculated by subtracting the mean fold expression value of the whole gene expression from the mean fold expression value of the metabolic gene-set, and divided the obtained value with the standard deviation of the whole fold expression value. Finally, the derived activity score was normalized by multiplying the obtained score with the square root of the number of genes in the gene-set to obtain a normalized z-score activity score for the metabolic process. Heatmap representation of the metabolic process activity levels was carried out using R v4.2.1 or dChip software. Metabolic process activity scores were then associated with EMT activation score by Pearson’s correlation method in R. A meta-correlation was performed using metacor.DSL function in R wherein the number of samples in each profile was included for the analysis and significantly associated pathways were identified with meta-r>0.3 and meta-r <-0.3 as positively and negatively associated, respectively, with meta-pvalue <0.05. Network visualization of the associated pathways was carried out using VisANT software.

### Survival analysis

Lung cancer gene expression profiles were obtained from GEO as normalized values and for TCGA profiles from cbioportal platform as mRNA z-score values. SP gene signature score was calculated similar to the z-score pathway activation with the difference of up-regulated and down-regulated activity scores in the patient samples. Samples were then categorized as SP-low and SP-high based on the median of the SP gene signature activity score and generated survival curve based on Kaplan-Meier estimate analysis. Significant difference between the two categorized samples was estimated using log-rank test in R software.

### Statistical analysis

All statistical analysis was performed using GraphPad Prism9 software or using R v4.2.1 software.

## Results

### Short-chain fatty acids show negative association with EMT in lung cancer

Towards exploration of inherent molecular or metabolic processes that could inhibit or reverse the EMT features in lung cancer, we performed an integrative functional genomic analysis in a comprehensive collection of patient-derived lung cancer gene expression profiles from 6 different datasets (N=1,476) (**Supplementary Table 1**). To perform this, we obtained a robust pan-cancer derived EMT gene signature containing 52 mesenchymal and 25 epithelial genes from a previous report [38] (**Supplementary Table 2**). The obtained EMT gene signature’s expression was validated in the contexts of genetic contributors of EMT (SNAIL, TWIST, GSC, TGF-β, and knockdown of E-cadherin) in HMLE cell line profile, and also upon treatment with TGF-β1 in multiple lung cancer cell line (A549, NCI-358 and HCC827) gene expression profiles. As expected, we found a clear increase in most of the mesenchymal genes expression in the induced-EMT state with a decrease in the epithelial genes expression (**Supplementary Figure 1**). Secondly, for the investigation of metabolic processes in the EMT context, we extracted, in total 335 metabolic oriented gene-sets derived from KEGG (n=135) and REACTOME (n=200) deposited in MSigDB signature database. In general, the collected KEGG and REACTOME metabolic gene-sets collectively represented major and several metabolic processes related to biochemical and enzymatic activities, metabolisms of carbohydrate, protein, lipid, long- and short-chain fatty acids, steroids, cholesterol, drug, nucleotide, organic and inorganic, acyl chain, alcohol, macromolecular, biopolymer, vitamins, co-factor and co-enzymatic activity.

An EMT-centered investigation was then performed using the pan-cancer EMT gene signature to identify the positively and negatively associated metabolic processes in lung cancer gene expression profiles using z-score based pathway activation approach and meta-correlation analysis as described in Methods. With a stringent cut-off of clinically accepted meta-correlation (r<-0.3 and r>0.3), we identified several previously reported positively and negatively associated processes with EMT (**Figure 1A**, **Supplementary Figure 2** and **Supplementary Table 3-4**). For instance, we observed chondroitin sulphate biosynthesis [39] or heparan biosynthesis [40] to have positive association with EMT gene signature including TGF signaling, ECM receptors interaction, and gap junction degradation. Interestingly, we found EMT to have negative associations with the short-chain fatty acids (SCFAs), propionate (meta r = −0.46; meta p-value<0.0001) and butanoate (meta r = −0.46; meta p-value<0.0001) metabolic processes (**Figure 1B-D**; **Supplementary Table 3**), which have not been investigated in detail previously in the context of lung cancer mediated EMT process. As the integrative genomic analysis was done with the microarray-based profiles, the analysis was also extended to TCGA pan-cancer profiles performed using RNA-Seq and observed a similar trend of negative associations between EMT and propionate (r=-0.46; p<0.0001) or butanoate (r=-0.53; p<0.0001) in lung adenocarcinoma patients (N=510) (**Supplementary Figure 3A-B**). The negative associations between SCFAs and EMT was further validated with different source of EMT gene signatures from the literature obtained from MSigDB (**Supplementary Figure 3C-F**), and with the GSEA analysis of Hallmark EMT gene signature significantly enriched in low-propionate or low-butanoate lung cancer patient samples (**Figure 1E**; **Supplementary Figure 3G**). Interestingly, a high survival probability of lung cancer patients with high-propionate or high-butanoate gene-set expression was observed implicating good prognosis of SCFAs (**Figure 1F**; **Supplementary Figure 4**).

**Figure 1.**
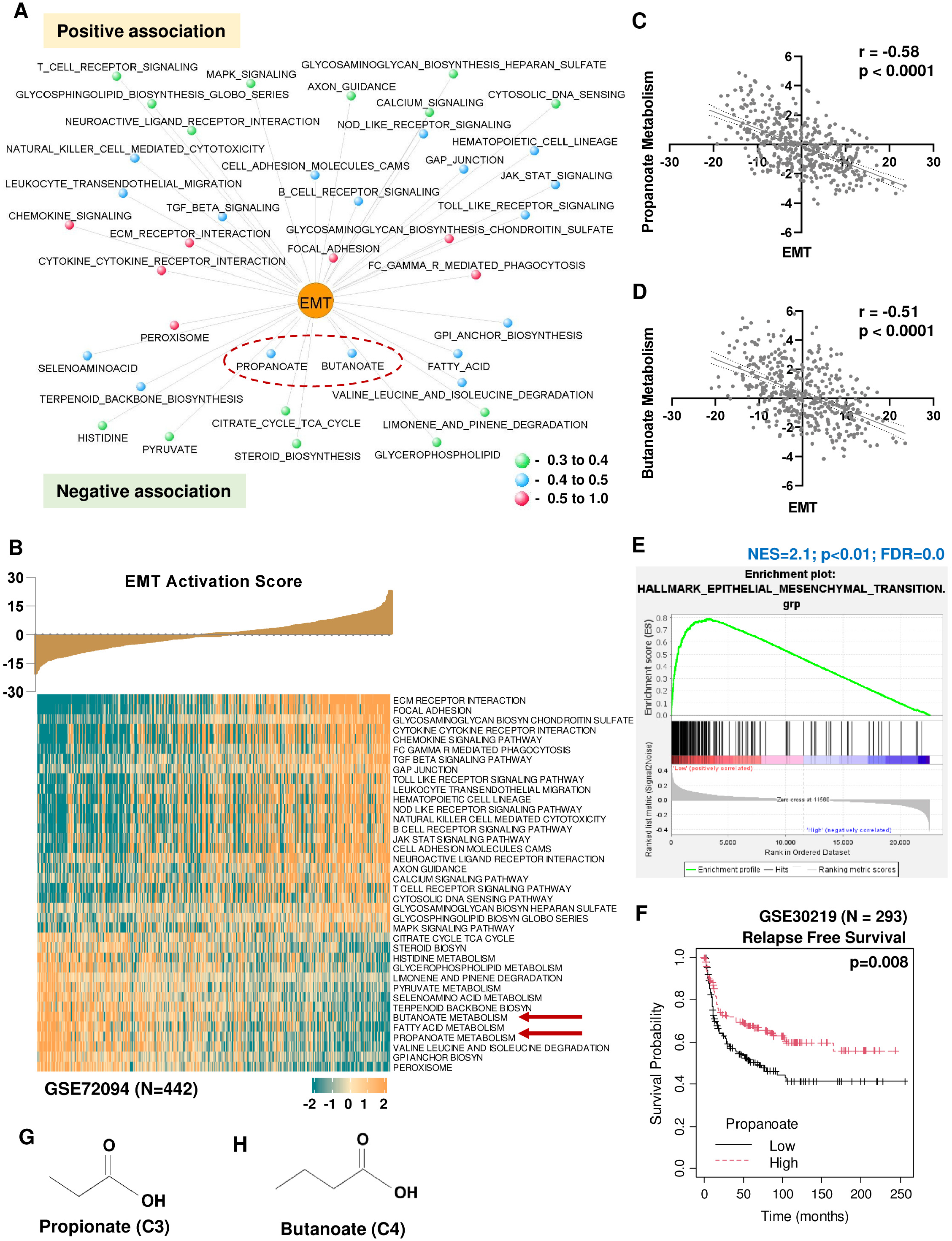
Propionate and butanoate are negatively associated with EMT in lung cancer gene expression profiles identified from integrative functional genomic analysis. A) Network visualization of positively and negatively associated metabolic processes with EMT inferred from the correlation analysis of the activation scores for EMT and metabolic processes of KEGG gene-sets in lung cancer gene expression profiles (N=1476) from 6 datasets. Significantly associated metabolic processes with EMT were assessed from the meta-correlation of all lung tumor datasets used in the analysis. meta-r<-0.3 and meta-r>0.3; meta-p<0.05. B) Representative heatmap visualization of the activation scores of significant positively and negatively associated metabolic processes gene-set with the increasing activation levels of EMT gene signature as a bar plot in GSE72094 (N=442) above heatmap. Propionate and butanoate metabolisms showing negative association with EMT activation levels were highlighted with red arrows. C&D) Correlation plot between the metabolic process activation scores of short-chain fatty acids (propionate (C) or butanoate (D)) with EMT gene signature activation in GSE72094 profile (N=442). E) Gene-set enrichment analysis of Hallmark EMT gene-sets with the lung cancer patient samples (GSE72094; N=442) categorized as low and high based on the propionate gene-set activation levels show an enrichment of EMT in low propionate patient samples. Ranking of genes with signal2noise metric was used for GSEA. F) Relapse free survival analysis in lung cancer patient samples (GSE30219; N=293) categorized as low- and high-propionate levels based on the median showed good prognosis for propionate gene-set. p-value was calculated using log-rank method. G&H) Structure of short-chain fatty acids, propionate (G) and butanoate (H).

Collectively, the integrative genomic analysis of metabolic processes in the context of EMT showed a negative role for SCFAs in EMT process with good prognostic events in multiple lung cancer gene expression profiles.

### Propionate and butanoate, but not acetate, are involved in EMT inhibiting processes with an increase in the functional E-cadherin expression

Having observed a negative association between short-chain fatty acids with pan-cancer EMT gene signature, *in vitro* action of predominantly found SCFAs in the human system (acetate, propionate and butanoate) was investigated in inhibiting or reversing the effect of EMT process. Treatment of A549, a NSCLC adenocarcinoma cell line, with sodium acetate (SA), sodium propionate (SP) and sodium butanoate (SB) showed a drastic increase in the protein expression of a key epithelial gene marker, E-cadherin, with a decrease in the mesenchymal master regulator, ZEB1, in a dose- and time-dependent manner for propionate and butanoate but not for acetate (**Figure 2A-C**). This indicates that the EMT inhibiting feature is merely specific for propionate (C3) and butanoate (C4) but not for acetate (C2) in the abundant short-chain fatty acid group in humans. The experiment was controlled using sodium chloride to rule out the effect of changes due to sodium ion (**Supplementary Figure 5A**). Further, this phenomenon was validated in multiple lung cancer cell lines such as with SKMES1, NCI-H520 and NCI-H23, and, for both propionate and butanoate treatments in a dose- and time-dependent manner (**Figure 2D-E**; **Supplementary Figure 5B-C**) with an increase in the E-cadherin with a decrease in the ZEB1 expression levels. Independently, by immunofluorescence, a clear increase in the E-cadherin expression was observed in propionate treated A549 and SKMES1 cell lines at the cytoplasmic membrane region with the increase seen at the juncture of cell-to-cell contact which is the site of E-cadherin’s functional role in cell adhesion and cell-to-cell contact (**Figure 2F**). Similar effect was also observed in pancreatic cancer cell line with epithelial morphology PANC1, indicating that other cancers could be subjected to EMT repression by SCFA (**Supplementary Figure 5D-F**). In addition, flow cytometry analysis of cell surface stained E-cadherin in the SP treated cells showed a substantial increase (2.9 folds) in mean fluorescence intensity in SP treated compared to control (**Figure 2G**; **Supplementary Figure 5G**). Thus, propionate treatment increases the functionally active E-cadherin levels important in the cell-to-cell contact establishment for the epithelial process.

**Figure 2.**
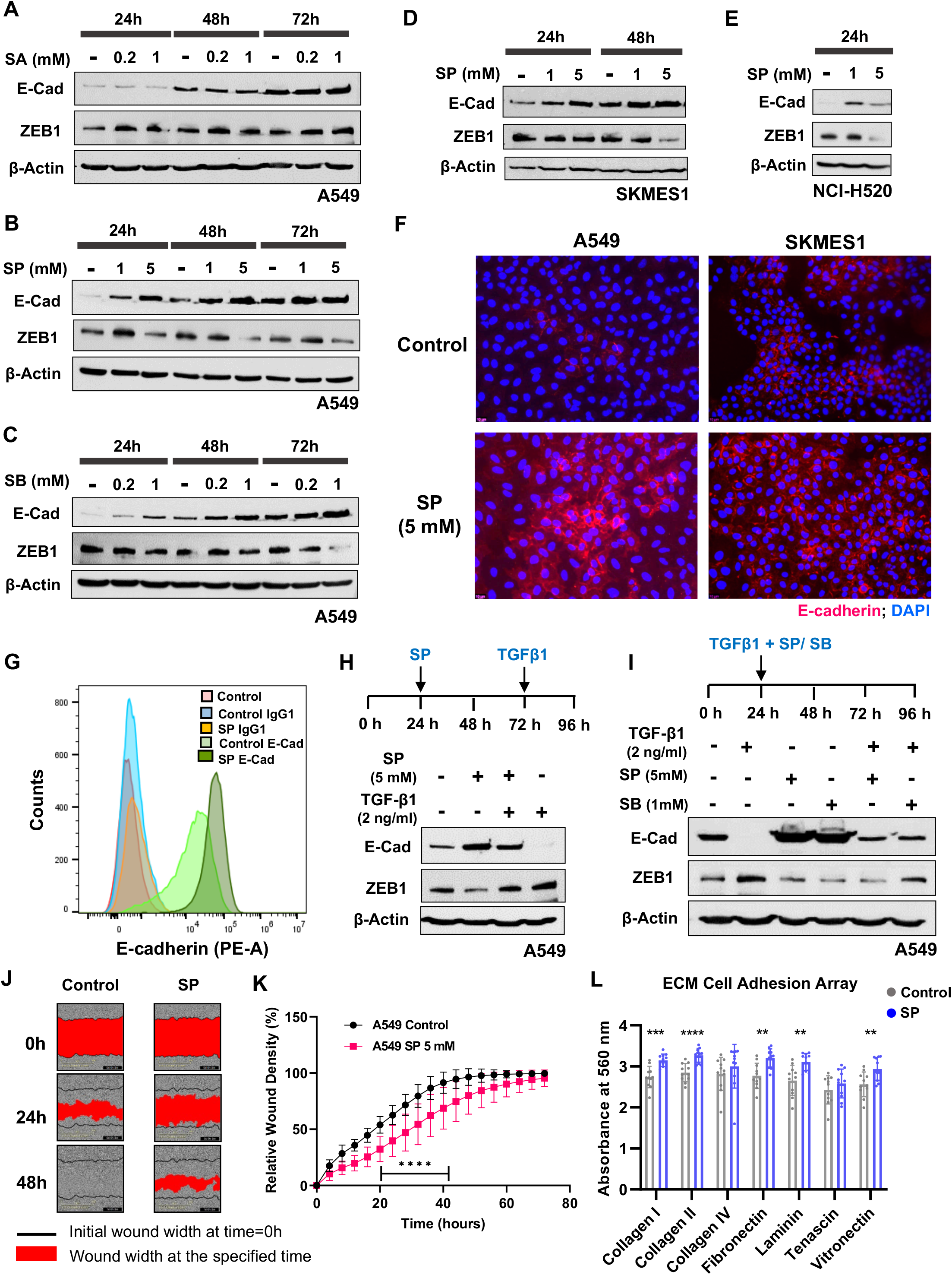
Chemopreventive effect of propionate and butanoate on EMT related markers. A-B) Western blot analysis of E-cadherin and ZEB1 protein levels in A549 cells treated with sodium acetate (SA) (A), sodium propionate (SP) (B) or sodium butanoate (SB) (C) in the indicated dose- and time-dependent manner. β-Actin was used as an internal control. D-E) Western blot analysis of E-cadherin and ZEB1 in NSCLC cell lines SKMES1 (D) and NCI-H520 (E) treated with sodium propionate (SP) in the indicated dose and time points. β-Actin was used as an internal control. F) Immunofluorescence staining of E-cadherin in NSCLC cell lines, A549 and SKMES1, treated with sodium propionate (5 mM) for 3 days. DAPI was used as a nuclear stain. Scale bar represents 10 μm. G) Flow cytometry analysis of membrane-associated PE-conjugated E-cadherin in A549 cells treated with sodium propionate (SP) at 5 mM concentration for 48h. PE-conjugated IgG1 was used as a control for flow cytometry. Control DAPI was used for dead cell exclusion and gating purpose. H) Western blot analysis of E-cadherin and ZEB1 protein levels in A549 cells pre-treated with sodium propionate (SP, 5 mM) for 48 hours followed by TGF-β1 (2 ng/ml) for 24 hours. β-Actin was used as an internal control. I) Western blot analysis of E-cadherin and ZEB1 protein levels in A549 cells co-treated with sodium propionate (SP, 5 mM) or sodium butanoate (SB, 1 mM) in combination with TGF-β1 (2 ng/ml) for 72 hours. β-Actin was used as an internal control. J) Images represent the wound width (highlighted by red region) at the indicated time points in A549 cells treated with SP for 3 days. K) Line plot depicts the rate of wound healing, expressed as relative wound density in A549 cells treated with SP over the course of 72 hours from making the wound. M) Cell adhesion assay in A549 cells treated with sodium propionate (5 mM) for 72 hours. Data points (n=8) are represented as mean ± SD and p-value was calculated from t-test. ****- p-value <0.0001; *** - p-value <0.001; ** - p-value <0.01.

Next, these SCFAs were tested for their ability to reverse or inhibit EMT upon treatment with TGFβ1, a known master regulator of EMT process in cancer. Treatment of A549 cells with SP or SB could not reverse the E-cadherin expression in 48 hours TGFβ1-induced EMT condition implying these metabolites have no ability to revert the phenotype with completely transitioned to the mesenchymal state (**Supplementary Figure 5H-I**). On the other hand, pre-treatment of A549 cells with SP or SB for 48 hours showed a substantial inhibitory effect on the TGFβ1-mediated EMT program. While cells lost E-cadherin expression with an increase in ZEB1 in TGFβ1 treated condition, SP or SB pre-treated cells with TGFβ1 almost retained the E-cadherin expression with the repression of ZEB1 (**Figure 2H**; **Supplementary Figure 5J**). In another instance, co-treatment of A549 cells with TGF-β1 and SP or SB for 72 hours could inhibit the TGF-β1 induced EMT to considerable extent with a decrease in the ZEB1 levels compared to TGF-β1 alone treatment (**Figure 2I**). This clearly suggest that propionate and butanoate can noticeably inhibit or halt the process of EMT progression, and that it could act as a chemopreventive agent in reducing the aggressive state of the cancerous condition.

Migration, another EMT feature, was also shown to have a significant inhibition till 48 hours in A549 cells pre-treated with SP (**Figure 2J-K**). To attribute the SP’s role for the enhanced epithelial program at the cellular level, cell adherence was assessed for several cell adhesion factors in A549 cells treated with SP for 3 days, and significantly observed an increase in the extracellular matrix (ECM) proteins implying propionate enhances the cell-cell contact and cell-to-surface contacts in the treated cells as observed with the increase in collagen I/II, fibronectin, laminin and vitronectin (**Figure 2L**).

Altogether, propionate and butanoate exhibit EMT inhibiting properties with an increase in the functionally active epithelial gene marker, E-cadherin, contributing to important function of cell-cell contact and cell adhesion.

### Propionate inhibits lung colonization and lymph node metastasis in vivo

Both propionate and butanoate showed a similar increase of E-cadherin expression and EMT inhibiting features from our *in vitro* study, and both are in clinical trials for various conditions like diabetes, obesity, inflammatory bowel disease and end stage renal disease. However, butanoate’s suitability in the clinical and nutrition context is limited due to its unpleasant odor and rapid absorption rate in the upper gastrointestinal tract [41,42], while propionate is a safe food ingredient with clinical benefits [43]. Therefore, we decided to focus more towards propionate’s role in lung cancer. First, the proliferative effect of propionate on A549 cells was tested and there was no significant change observed in the proliferation of cells till 12 days SP treatment (**Figure 3A**). An experimental lung metastasis model was then assessed wherein A549-PFUL2G cells (expressing constitutive luciferase gene) were treated with SP for 3 days followed by tail vein injection of the cells in NSG mice, and monitored the lung colonization of the cells using in vivo imaging of the luciferase activity after 3 days. There was a marked decrease in the lung colonization ability of SP treated cells compared to the control cells implying that propionate can inhibit metastasis (**Figure 3B-C**) independent of proliferation. The metastatic inhibitory ability of SP was further verified in another NSCLC cell line, SKMES1-PFUL2G, with a similar reduced lung colonization in NSG mice independent of proliferation (**Figure 3D-F**).

**Figure 3.**
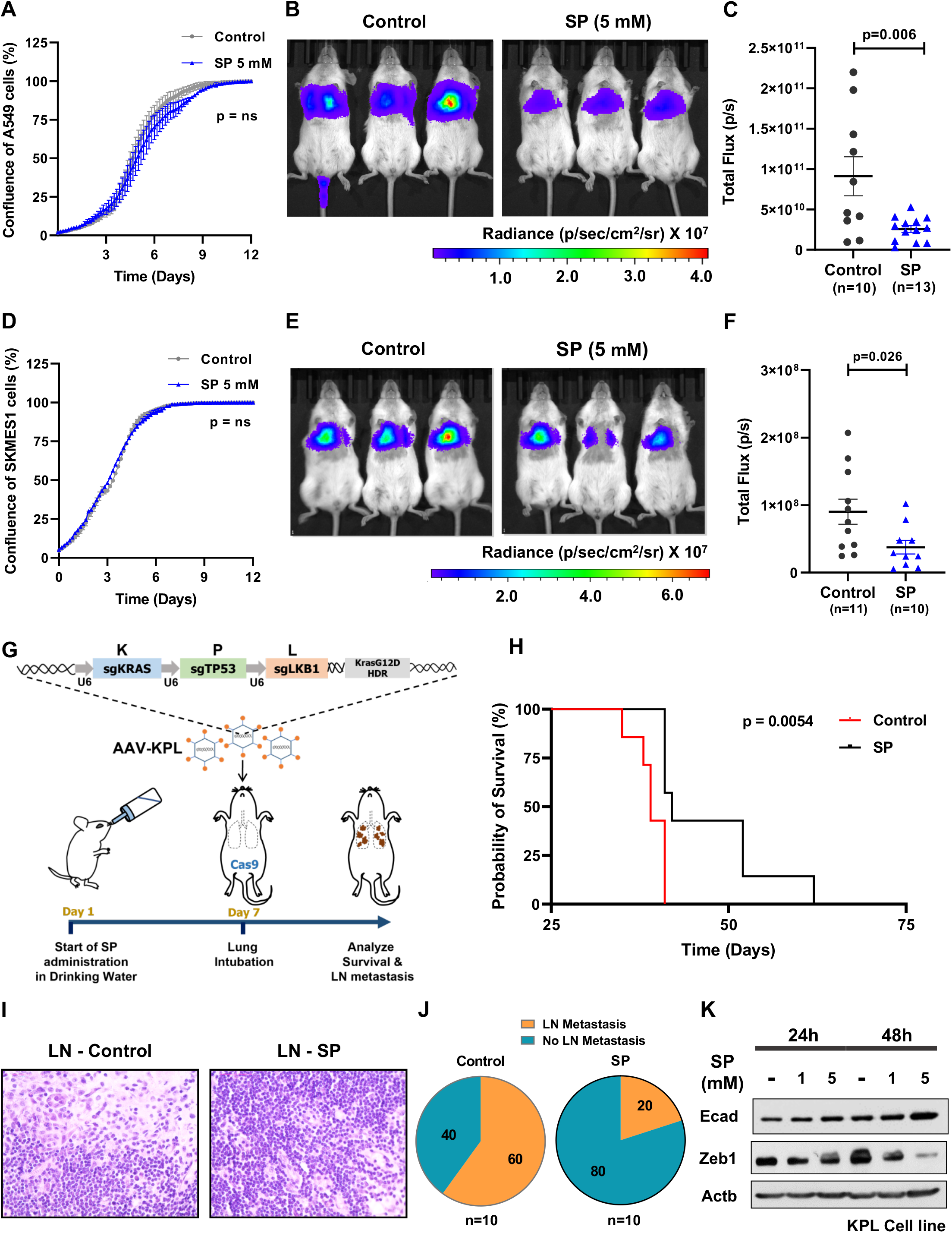
*In vivo* effect of propionate in lung metastatic mouse models. A) Real-time cell proliferation analysis of A549 cells treated with sodium propionate (5 mM) for 12 days (n=3). Data points are represented as mean ± SD and p-value was calculated from Two-way ANOVA. B) *In vivo* imaging of lung colonization ability of A549-PFUL2G cells treated *in vitro* with sodium propionate (5 mM) for 3 days followed by tail-vein injection in NSG mice. C) Quantification of luciferase activity as bioluminescence signal (total flux) in the lung metastatic NSG mice injected with A549-PFUL2G cells treated *in vitro* with sodium propionate (5 mM) for 3 days. Data are represented as mean ± SEM with significance calculated using un-paired two tailed t-test. D) Real-time cell proliferation analysis of SKMES1 cells treated with sodium propionate (5 mM) for 12 days (n=3). Data points are represented as mean ± SD and p-value was calculated from Two-way ANOVA. E) *In vivo* imaging of lung colonization ability of SKMES1-PFUL2G cells treated *in vitro* with sodium propionate (5 mM) for 3 days followed by tail-vein injection in NSG mice. F) Quantification of luciferase activity as bioluminescence signal (total flux) in the lung metastatic NSG mice injected with SKMES1-PFUL2G cells treated *in vitro* with sodium propionate (5 mM) for 3 days. Data are represented as mean ± SEM with significance calculated using un-paired two tailed t-test. G) Schematic representation of analysis of survival probability and lymph node metastatic inhibitory activity of sodium propionate by administering in drinking water to AAV-KPL virus lung intubated C57BL/6 mice. H) Survival analysis of C57BL/6 mice with lung tumorigenesis intubated with AAV-KPL virus and administered with sodium propionate (SP) in drinking water. Significance was calculated using log-rank method. I) Representative images of lung cancer metastasis to axial lymph nodes in C57BL/6 mice of control (LN-Control) condition while no metastasis in the sodium propionate (LN-SP) condition. Scale bar represents 200 μm. J) Pie chart representation of percent lung cancer metastasis to axial lymph nodes in C57BL/6 mice administered with sodium propionate (SP) in drinking water. K) Western blot analysis of E-cadherin and Zeb1 levels in KPL cell line derived from C57BL/6 mice with lung tumorigenesis intubated with AAV-KPL virus. β-Actin was used as an internal control. ns – non-significant.

To further evaluate the positive role of propionate’s metastasis inhibiting property, another mouse metastatic model was employed wherein gender- and age-controlled Cas9-C57BL/6 mice were administered with sodium propionate in drinking water (150 mM) as pre-treatment from one week before lung intubation of KPL adenovirus to assess survival and lymph node metastasis (**Figure 3G**). KPL adenovirus carrying KRAS G12D oncogenic mutation with loss-of-function mutations in TP53 and LKB1 [44] develops tumors in Cas9-bearing C57BL/6 mice when delivered oropharyngeal in the lungs, and the tumors can metastasize to the sentinel axial lymph nodes. Oral administration of SP was continued throughout the study. There was no difference in the drinking water consumption or body weight between the groups (**Supplementary Figure 6A-B**). Interestingly, mice administered with sodium propionate showed a significant increase in the survival time (**Figure 3H**) and a reduced lymph node metastasis was observed in mice administered with SP (60% lymph node metastasis in untreated versus 20% lymph node metastasis from SP treated animals) (**Figure 3I-J**). Necroscopic analysis of mice showed no significant difference in the number of primary tumor lesions in the lungs between the groups (**Supplementary Figure 6C-D**). Further, a KPL-derived cell line treated with sodium propionate showed an increase in the E-cadherin protein levels along with a decrease in the Zeb1 levels in a dose- and time-dependent manner indicating that KPL model is strongly responsive to EMT inhibition by sodium propionate (**Figure 3K**).

Overall, these *in vivo* results strongly support that sodium propionate possess an EMT inhibiting feature by reducing the metastasis as observed with the lung colonization ability, and with improved survival rate of the KPL-tumor bearing mice with reduced lymph node metastasis by dietary administration of sodium propionate.

### Sodium propionate increased the cisplatin sensitivity of the cells *in vitro*

EMT is a known feature of chemoresistance and lung cancer patients develop chemoresistance during the treatment regimen to cisplatin, a widely used primary line of treatment for the patients [45,46]. Therefore, SCFAs were evaluated for their increase in the sensitivity of the cells to cisplatin. Pre-treatment of cells with sodium propionate or butanoate for 48 hours followed by cisplatin treatment in a dose-dependent manner significantly showed a decreased cell growth ability as evaluated by real-time cell imaging compared to control cells in both A549 (**Figure 4A**; **Supplementary Figure 7A**) and SKMES1 cell lines (**Figure 4B**; **Supplementary Figure 7B**) suggesting an improved sensitivity to cisplatin by propionate or butanoate. In addition, cell death was more pronounced with propionate (or butanoate) in combination with cisplatin in a dose-dependent manner compared to the control (**Figure 4C-E**; **Supplementary Figure 7C**). Notably, the increased cell death difference was significant even with low dose of cisplatin at 2.5 μM in combination with SP compared to the control and better than butanoate’s effect (**Supplementary Figure 7D**). Increased sensitivity of cells to cisplatin by propionate was further validated with the γH2AX levels, a sensitive biomarker of double strand DNA damage, which showed a clear increase in the protein levels with the pre-treatment of A549 or SKMES1 cells with sodium propionate followed by cisplatin treatment (**Figure 4F-G**). This implicates that compared to cisplatin standalone treatment, pre-treatment with SP increases the cisplatin’s action of DNA damage in the cancerous cells. This was further confirmed by quantitative image-based cytometry (QIBC) analysis performed with the SP pre-treated cells in combination with cisplatin, which showed a clear dose-dependent increase in the levels of γH2AX at the single cell (**Figure 4H**), and at foci count levels (**Figure 4I**). Interestingly, RAD51 and TP53BP1, markers of homologous recombination and non-homologous end-joining repair process, respectively, did not show any difference in SP treated cells compared to the control with no changes in cell cycle phases (**Supplementary Figure 8A-F**) indicating that SP increases the sensitivity of cells towards cisplatin without impacting DNA damaging effect. Thus, sensitivity of cisplatin could be improved with the pre-treatment scenario for the treatment purposes.

**Figure 4.**
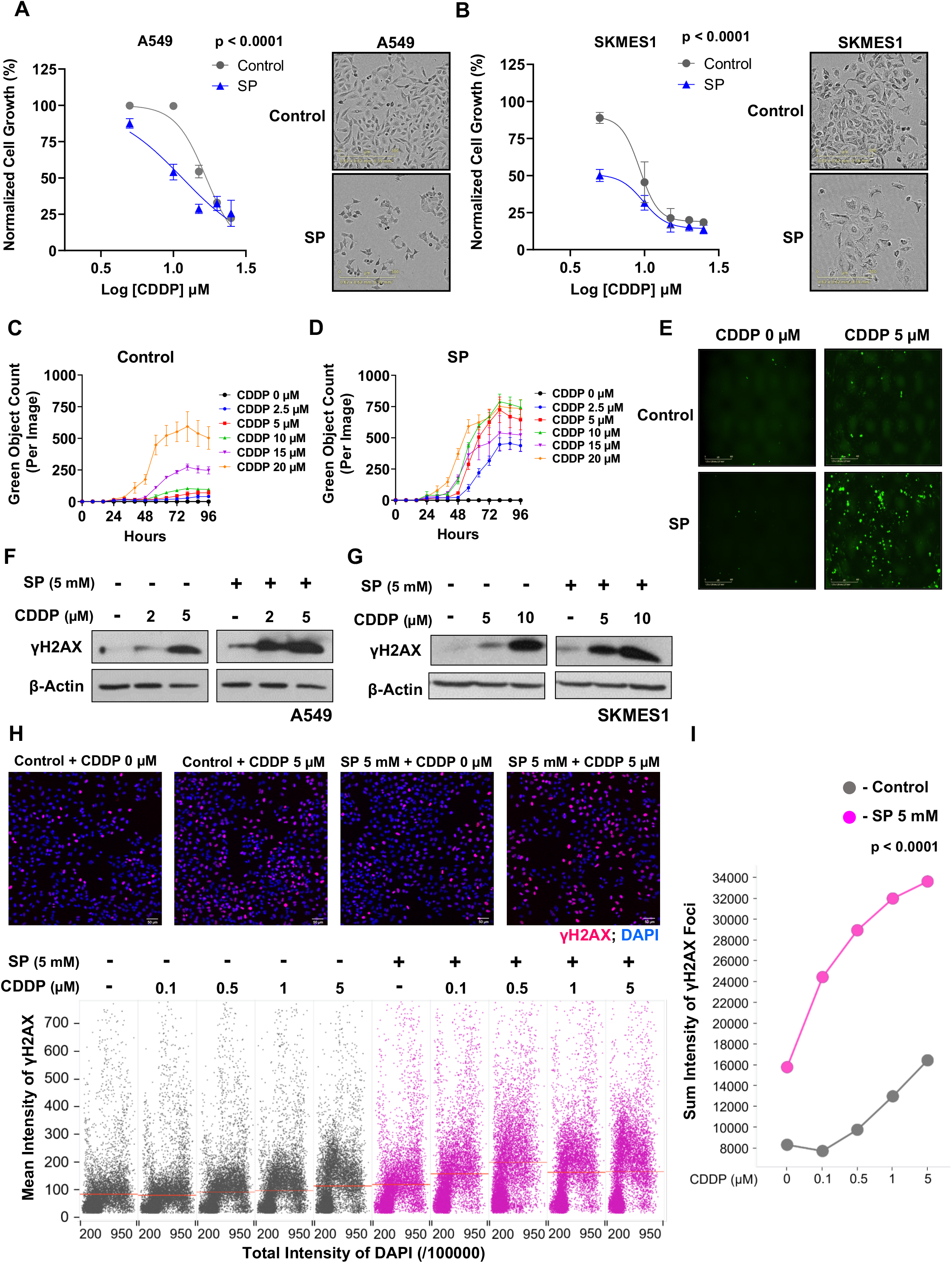
Sodium propionate sensitizes cells to cisplatin in lung cancer cell lines. A) Dose responsive curve of cisplatin treatment in combination with SP (5 mM) in A549 cell line. Data points are represented as mean ± SD normalized to control with p-value calculated from Two-way ANOVA analysis. In the right, images represent A549 cell line growth inhibition by cisplatin (10 μM) treated in the presence and absence of SP. Scale bar represents 300 μm. B) Dose responsive curve of cisplatin treatment in combination with SP (5 mM) in SKMES1 cell line. Data points are represented as mean ± SD normalized to control with p-value calculated from Two-way ANOVA analysis. In the right, images represent SKMES1 cell line growth inhibition by cisplatin (5 μM) treated in the presence and absence of SP. Scale bar represents 300 μm. C-D) Quantification of dead cells as green object count using Cytotox Green in A549 cells treated with dose-dependent cisplatin in the absence (C) and presence of SP (D). Data points are represented as mean ± SD. E) Representative images of A549 cells treated with cisplatin (5 μM) in combination with SP (5 mM). Scale bar represents 400 μm. F-G) Western blot analysis of γH2AX levels in A549 cells (F) and SKMES1 cells (G) treated with cisplatin in the indicated dose-dependent concentrations along with SP (5 mM). β-Actin was used as an internal control. I) QIBC analysis of γH2AX levels in A549 cells treated in the indicated dose-dependent concentrations of cisplatin for around 6 hours in combination with SP pre-treatment (5 mM) for 24 hours (bottom). Significance (p<0.0001) was calculated using Two-way ANOVA. Top, Images represent the γH2AX levels in A549 cells treated in the presence and absence of cisplatin (5 μM) along with SP (5 mM). DAPI was used as a nuclei stain. Scale bar represents 50 μm. J) QIBC analysis of sum intensity levels of γH2AX foci in A549 cells showing an increasing trend with the treatment of cisplatin in the indicated dose-dependent concentrations in combination with SP (5 mM). p-value was calculated using Two-way ANOVA.

### Expression profiling of propionate reveals enhanced lung epithelial features in inhibiting EMT

Having observed propionate’s role in inhibiting EMT-mediated metastasis with increased cisplatin sensitization, the molecular mechanism of propionate’s action behind these features was investigated by gene expression profiling. Initially, the consistency of propionate mediated E-cadherin expression in A549 cell line was tested for longer time periods over a course of 12 days, and observed consistent increased E-cadherin expression with decrease in ZEB1 expression both at the translation and transcription levels (**Figure 5A**; **Supplementary Figure 9A-B**). Therefore, RNA-Seq profiling was performed in A549 cells treated with sodium propionate (5 mM) for 3 days and 12 days. PCA and unsupervised hierarchical clustering analysis showed high variance and separate cluster of SP treated samples from the control, respectively, and identified differentially expressed genes between the conditions (**Figure 5B**; **Supplementary Figure 9C-E**). The up-regulated and down-regulated differentially expressed genes from 3 days and 12 days SP treatment showed 65.3% and 49.7% overlap, respectively, indicating that there is a similar level of gene expression maintenance over the longer period of SP treatment (**Supplementary Figure 9F-G**). Moreover, the expression pattern in SP condition was analyzed and validated at multiple levels: i) Real-time quantitative PCR analysis showed several epithelial genes expression with significantly high folds (*KRT18, KRT19, EPCAM, ICAM1* and *CDH1*) (**Figure 5C**), ii) a significant enrichment of hallmark apical junction and apical surface gene-sets were observed with SP 3 days and 12 days gene-sets activation levels in lung cancer patient samples (GSE72094, N=442) from GSEA (**Figure 5D**; **Supplementary Figure 10**), iii) immunofluorescence staining of few important epithelial gene markers involved in cell adhesion (EPCAM) and tight junctions (ZO-1) showed a similar increase in the membrane expression of these markers with propionate’s treatment in A549 cells (**Figure 5E**). Importantly, iv) an increased protein levels of transcription factors important for epithelial program, such as GRHL1 [47] and OVOL2 [48], upon SP treatment was observed (**Figure 5F**).

**Figure 5.**
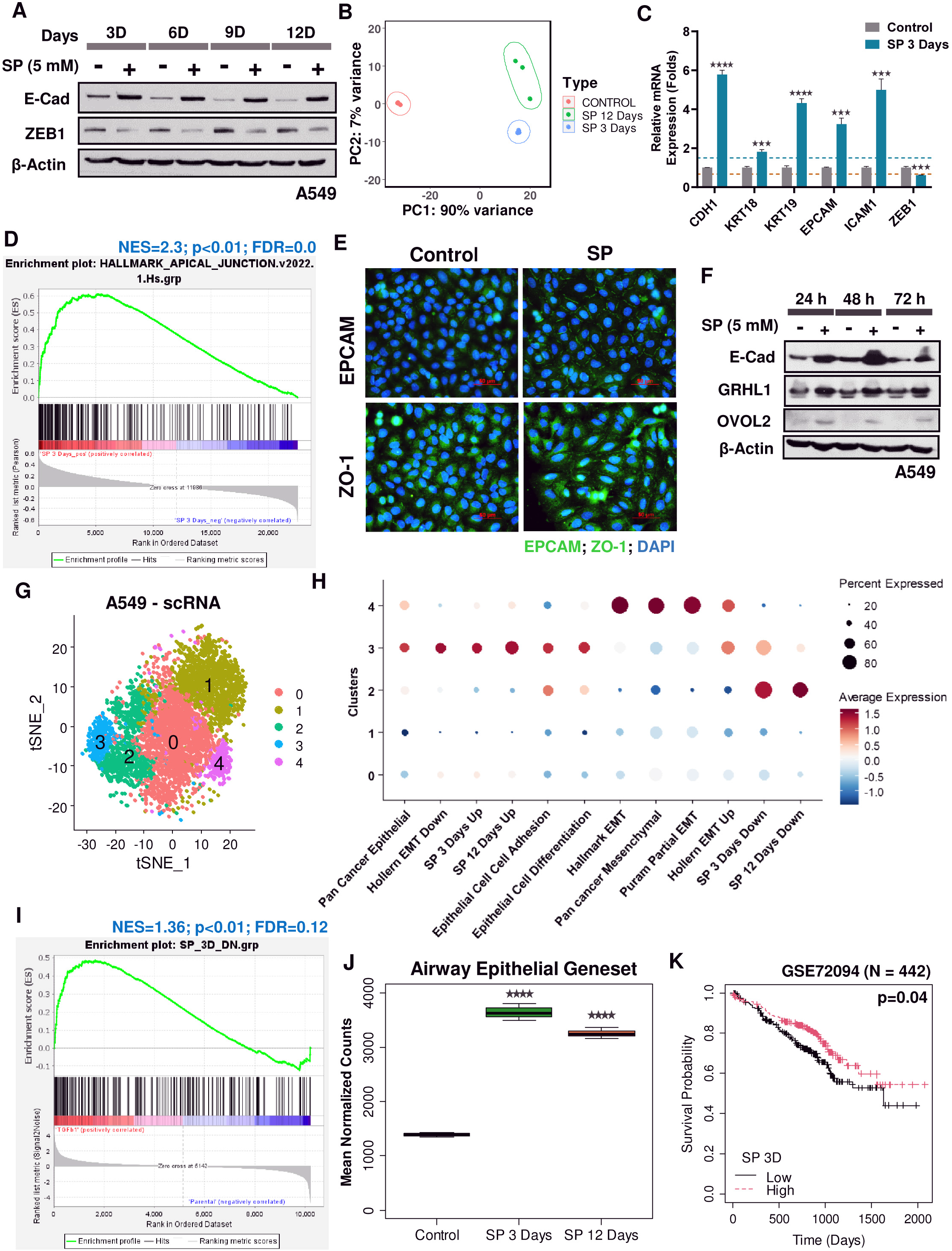
RNA-Seq expression profiling of SP treated A549 cell line reveals enriched epithelial gene expression. A) Western blot analysis of E-cadherin and ZEB1 protein expression in A549 cell line treated with sodium propionate at 5 mM in time series for 12 days. β-Actin was used as an internal control. B) Principal component analysis of RNA-Seq expression profile of A549 cell line treated with SP for 3 days and 12 days reveals a high variance in the gene expression profiles between the treated and the control samples. C) Real-time quantitative PCR analysis of epithelial genes (*CDH1, KRT18, KRT19, EPCAM* and *ICAM1*) and mesenchymal gene (*ZEB1*) in A549 cell line treated with sodium propionate (SP) at 5 mM for 3 days. *GAPDH* was used as an internal control. Blue and red dotted line represents the fold change cut-off at 1.5 and 0.67, respectively. Significance was calculated using unpaired t-test. D) Gene-set enrichment analysis of Hallmark Apical Junction gene-set enrichment in a lung cancer gene expression profile GSE72094 (N=442) as continuous label of SP 3 days gene-set z-score activity. Ranking of genes was based on Pearson’s correlation metric in GSEA. E) Immunofluorescence staining of EPCAM and ZO-1 in A549 cell line treated with sodium propionate (SP) at 5 mM for 3 days. DAPI was used as a nuclear stain. Scale bar represents 50 μm. F) Western blot analysis of GRHL1 and OVOL2 protein levels in A549 cell line treated in time series for 3 days with sodium propionate (SP) at 5 mM showed an increase in the epithelial-specific transcription factors expression. β-Actin was used as an internal control. G) Single cell RNA sequencing of A549 cell line revealed five different clusters of cells obtained from t-SNE analysis with a resolution of 0.4, perplexity of 50 from first 25 principal components. H) Dot plot pattern analysis of epithelial and mesenchymal gene-sets using AddmoduleScore analysis in Seurat showed the enrichment of SP up-regulated genes along with the epithelial gene-sets in cluster 3 of A549 cells from scRNA sequencing. I) Gene-set enrichment analysis of SP 3 days down-regulated gene set in TGFβ1-induced HMLE cell line compared to the control obtained from GSE24202 (n=3). Ranking of genes with signal2noise metric was used for GSEA. J) Box plot visualization of the enriched airway epithelial cell-type associated gene-set in SP 3 days and SP 12 days samples compared to the control. Significance was calculated using unpaired t-test between the SP treated cells and the control. K) Overall survival analysis in lung cancer patient samples (GSE72094; N=442) categorized as low- and high-propionate levels of SP 3 days gene-set z-score activity based on the median showed good prognosis. p-value was calculated using log-rank method. ****- p-value <0.0001; *** - p-value <0.001.

Next, we compared the RNA-Seq data with single cell RNA sequencing performed on the same A549 parental cells at baseline and interestingly, delineated a cluster of epithelial-like (cluster 3) and mesenchymal-like (cluster 4) features defined from the gene content overlap analysis of cell cluster’s marker genes using Enrichr (**Figure 5G**; **Supplementary Figure 11A**; **Supplementary Table 5**). Enrichr was performed for the marker genes in each cluster for the PanglaoDB and MSigDB collections. While cluster 3 marker genes were significantly enriched with epithelial cell-types including airway goblet cells and apical junction feature, cluster 4 marker genes were enriched with mesenchymal cell-types including fibroblasts and stromal cells together with the enrichment of EMT process, TGF-β1, TNFα signaling via NFκB and hypoxia signaling pathways compared to other clusters (**Supplementary Figure 11B-J**). This suggests that parental A549 cells are partial-EMT carrying hybrid-modes of epithelial-like and mesenchymal-like cells. With the defined clusters from scRNA-Seq, we analyzed the percent gene expression pattern of several epithelial-like and mesenchymal-like gene-sets along with the SP 3 days and 12 days gene-sets using modular score evaluation in Seurat scRNA analysis tool. Interestingly, A549 cells inherently possessing the epithelial-like cluster was found to have the SP up-regulated gene-sets from 3 days and 12 days in cluster 3 along with other epithelial-associated gene-sets although SP down-regulated gene-sets were not found in cluster 4 which possess partial EMT and mesenchymal gene-sets activity (**Figure 5H**). In contrast, the SP-down regulated genes were found in cluster 2 which is enriched with KRAS signaling gene-set (**Supplementary Figure 11C**). Nevertheless, GSEA of SP down-regulated gene-sets from 3 days or 12 days were found enriched in the genetically-induced EMT conditions (TGFβ1, TWIST, GSC and SNAIL) in HMLE cell line, respectively (**Figure 5I**; **Supplementary Figure 12**). This implies that propionate down-regulates the genes involved in the TGFβ1-induced or other master EMT regulators.

On the other hand, to further confirm the role of SP’s involvement in lung epithelial feature enhancement, the expression pattern of epithelial gene-sets related to lungs and other epithelial structural processes were examined. As expected, an increased expression levels (mean normalized counts) of gene-sets related to airway epithelial cells, markers of lung epithelial cells, pulmonary alveolar type II, and establishment of epithelial cell polarity, including hallmark apical surface were found highly expressed in both SP 3 days and 12 days compared to control (**Figure 5J**; **Supplementary Figure 13A-D**). Thus, this suggests the responsiveness of lung epithelial tissues to propionate. Survival analyses of 3 days and 12 days SP gene-sets were observed to have a significant good prognosis in lung cancer patients (**Figure 5K**; **Supplementary Figure 13E**).

Altogether, by independent approaches of validations, RNA-Seq expression profiling of propionate treated cells showed robust epithelial reprogramming in the context of lung along with the inhibition of mesenchymal processes.

### Propionate reinforces epithelial identity via chromatin remodelling

To understand the molecular mechanism of propionate’s mediation of E-cadherin expression or epithelial gene regulation, a search for small molecule mimics based on the biological gene signature of widely used pharmacological compounds was performed in L1000CDS^2^ search engine with the derived SP-regulated gene-sets from 3 days and 12 days. Based on the similarity metrics, HDAC inhibitors (vorinostat, trichostatin, and mocetinostat) were inferred in mimicking sodium propionate’s expression pattern (**Figure 6A**, **Supplementary Figure 14A**; **Supplementary Table 6-7**). Therefore, the epigenetic role of propionate’s mechanism was investigated in E-cadherin regulation. Upon treatment with HDAC inhibitors (vorinostat, mocetinostat and MI192), an increase in the E-cadherin levels was observed at protein levels (**Figure 6B**; **Supplementary Figure 14B-C**). However, the increase in E-cadherin was not an additive effect in combination treatment with SP. On the contrary, treatment with HAT inhibitor (HAT inhibitor CTK7A) in A549 and SKMES1 cell lines blocked the effect of SP’s action in E-cadherin expression in a dose- and time-dependent manner along with an increase in ZEB1 expression (**Figure 6C**). This suggests that propionate could regulate the expression of E-cadherin or epithelial program in more of an acetylation manner with epigenetic mode of regulation.

**Figure 6.**
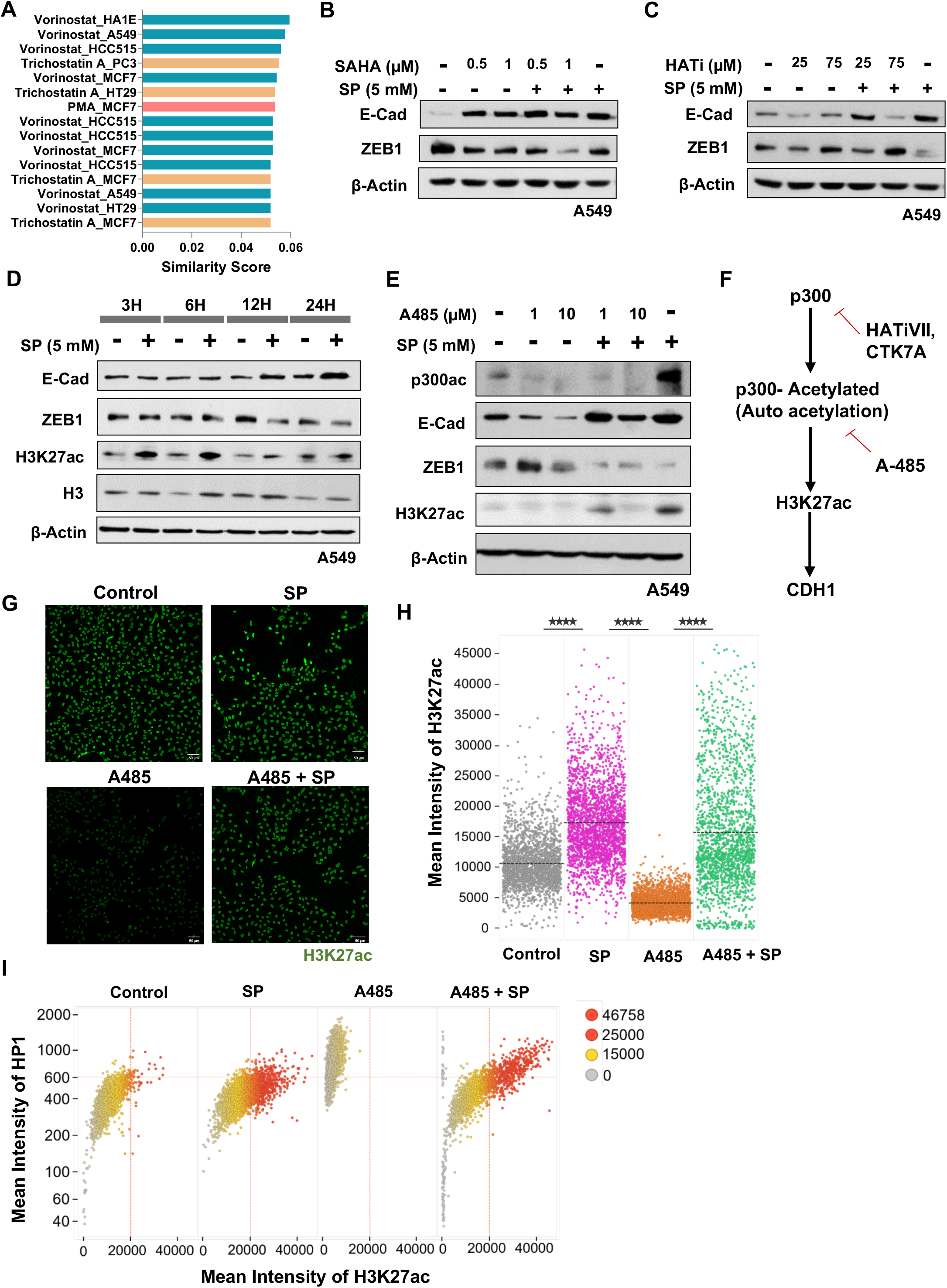
Sodium propionate modulates E-cadherin expression through p300 acetylation. A) Bar plot indicates the top ranked drug gene signatures similar to SP 12 days gene-set based on the similarity score identified from L1000CDS^2^. Identical drugs from different treatment conditions are colored the same. B) Western blot analysis of E-cadherin and ZEB1 protein expression in A549 cells treated with vorinostat (SAHA) in the indicated dose-dependent concentrations in combination with sodium propionate (5 mM) for 24 hours. β-Actin was used as an internal control. C) Western blot analysis of E-cadherin and ZEB1 protein expression in A549 cells treated with Histone Acetyl Transferase inhibitor VII, CTK7A (HATi) in the indicated dose-dependent concentrations in combination with sodium propionate (5 mM) for 24 hours. β-Actin was used as an internal control. D) Western blot analysis of E-cadherin, ZEB1, H3K27ac and H3 in A549 cells treated with sodium propionate (5 mM) in time series for 24 hours. β-Actin was used as an internal control. E) Western blot analysis of p300 acetylation, E-cadherin, ZEB1 and H3K27ac in A549 cell line treated with HAT inhibitor, A-485, in the indicated dose-dependent concentrations in combination with sodium propionate (5 mM) for 24 hours. β-Actin was used as an internal control. F) Schematic flowchart of sodium propionate’s role in the regulation of E-cadherin expression through p300 histone acetyl transferase activity on H3K27ac mark. G) Images represent the H3K27ac mark analysis by QIBC in A549 cells treated with A-485 (10 μM) in the presence and absence of sodium propionate (5 mM) along with SP (5 mM). DAPI was used as a nuclei stain. Scale bar represents 50 μm. H) QIBC analysis of mean intensity levels of H3K27ac mark in A549 cells treated with A-485 (10 μM) in the presence and absence of SP (5 mM) for 24 hours. Significance was calculated using One-way ANOVA with post-hoc analysis of t-test between the conditions. I) Biplot representation of mean intensity of HP1 to H3K27ac mark levels from QIBC analysis in A549 cells treated with A-485 (10 μM) in the presence and absence of SP (5 mM) for 24 hours. Color intensity values represent the mean intensity levels of the histone marks with grey representing null to red indicating high intensity. ****- p-value <0.0001

Other possible propionate’s mechanisms of action such as GPR signaling [49], MCT transporters [50] and PPAR signaling [51] were investigated (**Supplementary Figure 15A**). However, knockdown of critical genes (*PCCB, MCEE, MUT, HIBCH* and *ALDH6A1*) involved in propionate’s main and shunt metabolism showed no changes in E-cadherin expression (**Supplementary Figure 15B-C**). Similarly, i) blocking MCT1 and MCT4 transporters with AZD3965 or syrosingopine (**Supplementary Figure 15D-E**), ii) modulating GPR41 signaling with AR420626 (**Supplementary Figure 15F**) or iii) modulating PPAR pathway using inhibitor (GW9662) or activator (Troglitazone) (**Supplementary Figure 15G-H**) had no effects in E-cadherin regulation.

Therefore, an in-depth investigation was extended on the SP’s role of HAT mediated E-cadherin regulation. A recent report identified SCFAs (propionate and butyrate) role in acetylation event by inducing auto-acetylation of p300, and that this regulation involves H3K27ac [52] which was in line with our study. A time series experiment was performed to know the initial time point of high H3K27ac induction as well as E-cadherin expression with SP treatment, and found that H3K27ac was induced as early as at 3 hours, whereas E-cadherin was found expressed at 12 hours implying strongly that acetylation event precedes the E-cadherin expression upon propionate’s treatment (**Figure 6D**). Next, treatment with A-485, a specific catalytic inhibitor of p300/CBP [53], in A549 cell line at 24 hours observed to have a clear positive regulation of p300 in H3K27ac mediated E-cadherin expression by propionate. E-cadherin level was found decreased with A-485 in combination with SP, while H3K27ac was ablated at 24 hours (**Figure 6E**). More importantly, the level of acetylated-p300 was found increased with the SP treatment compared to the control strongly signifies that propionate induces p300-mediated signaling event with an increased H3K27 acetylation process (**Figure 6E-F**). This was also verified in SKMES1 cell line (**Supplementary Figure 16A**). Further, A-485 independently decreased E-cadherin expression on its own in a dose-dependent manner at 24 hours. There was no effect of H3K27me3 in E-cadherin regulation when treated the cell lines with EZH2 methyltransferase inhibitors (GSK126 or GSK343) in combination with sodium propionate (**Supplementary Figure 16B-C**). This indicates the specificity of H3K27 acetylation’s involvement in propionate’s mechanism of E-cadherin expression. Moreover, QIBC analysis detected a considerable increase of H3K27ac levels in SP treated cells compared to the controls. While A-485 treatment alone abrogated the acetylation event, combination treatment with SP preserved H3K27ac significantly (**Figure 6G-H**; **Supplementary Figure 16D**). Similarly, HP1, marker of heterochromatin formation, was found increased with A-485 treatment (**Supplementary Figure 16E**). Biplot of mean intensity expression of H3K27ac and HP1 showed that A-485 treated cells contained increased HP1 levels, whereas in combination with sodium propionate the process was reverted with an enhanced H3K27ac levels pointing towards a scenario of an involvement of global transcriptional activation involving active promoters and enhancers (**Figure 6I**).

In order to identify the precise increase in H3K27ac by SP as a preceding event with the occupancy of the mark at the promoter region of E-cadherin, ChIP-Seq experiment was performed for H3K27ac mark with 3 hours SP treated cells (**Supplementary Figure 17A**). Interestingly, a global acetylation event was detected with SP treatment, wherein 5,209 promoter-localized sites and 10,743 enhancer-localized sites were found acetylated with sodium propionate treatment (**Figure 7A**). A more pronounced acetylation event was noticed at the promoter site of *CDH1* gene indicating a definite occupancy by H3K27ac mark in regulating the gene expression (**Figure 7B**). A parallel RNA-Seq for 3 hours and 24 hours propionate treated cells was also performed. PCA and unsupervised hierarchical clustering analysis showed high variance and separate cluster of SP treated 24 hour samples from the control and 3 hours, respectively, and identified differentially expressed genes between the conditions (**Figure 7C**; **Supplementary Figure 17B**). As expected, up-regulated and down-regulated gene-sets from 24 hours SP treated samples had apical junction and hallmark EMT enrichment, respectively (**Supplementary Figure 17C-D**). In addition, single-sample scoring of EMT gene signature in 3 hours and 24 hours SP treated individual RNA-Seq samples showed a clear time-dependent decrease in the EMT gene signature [6] score significantly from 3 hours to 24 hours compared to the control (**Figure 7D**). In other words, SP preferentially induces the epithelial-specific gene expression program from 3 hours onwards with high expression reaching towards 24 hours. The global acetylation program by propionate on the active gene program was also investigated with the integration of ChIP-Seq profile with the RNA-Seq profile by binning the gene expression in the control cells based on low-to-high levels (**Figure 7E**). Increased global acetylation event tries to reprogram the chromatin in a manner that tends the low expressing genes towards active gene expression with the occupancy of H3K27ac marks at their promoters, while minimal levels of acetylation increase was far sufficient in promoting the expression of active genes in high levels.

**Figure 7.**
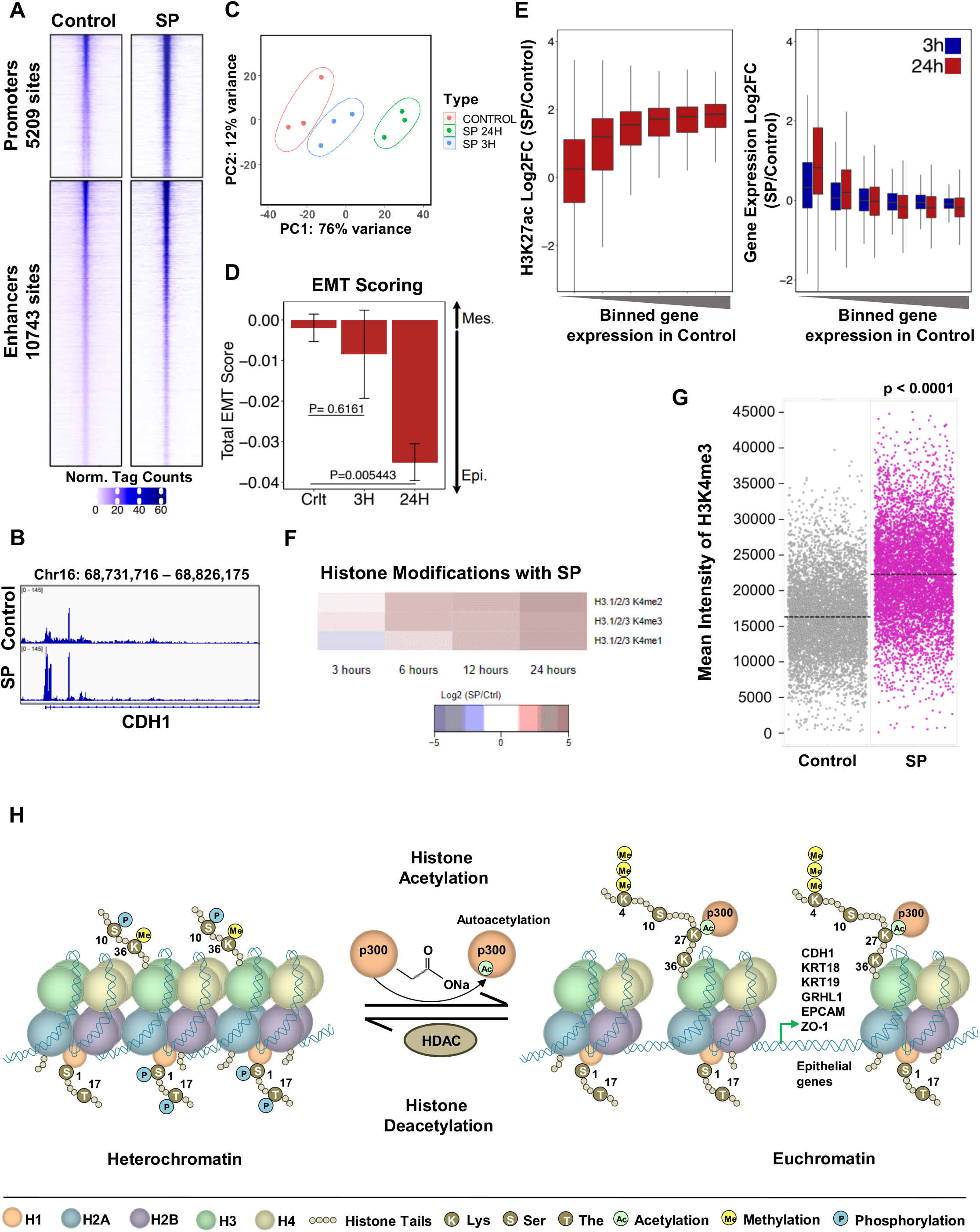
Sodium propionate induces epigenetic re-programming towards transcriptional initiation or activation of epithelial oriented program. A) Heatmap visualization of ChIP-Sequencing for H3K27 acetylation (H3K27ac) mark enrichment at promoters and enhancers sites in A549 cells treated for 3 hours with sodium propionate (SP, 5 mM). Color intensity scale bar represents the normalized tag counts or read density at a given region. B) UCSC Genome browser screen shot visualization of H3K27ac peak enrichment at *CDH1* promoter region identified from ChIP-Seq for H3K27ac in A549 cells treated for 3 hours with sodium propionate (SP, 5 mM). Peaks represented in the tracks are the normalized FPKM counts. C) Principal component analysis of RNA-Seq expression profile of A549 cell line treated with SP for 3 hours and 24 hours. D) Bar plot represents the EMT score for control, 3 hours and 24 hours RNA-Seq samples (n=3) of A549 cells treated with sodium propionate. EMT score is the difference between the mesenchymal gene-set score and epithelial gene-set score obtained from SingScore. p-value was calculated from Welsch’s two-sided t-test. E) Box plot visualization of H3K27ac and gene expression changes with sodium propionate treatment in A549 cells for all expressed genes sorted low-to-high binning of gene expression from control. Left plot represent the log2 fold difference in H3K27ac profile during SP treatment for 3 hours while the right plot represents the log2 fold difference in the gene expression profile of SP for 3 hours and 24 hours. F) Heatmap representation of top modulated post-histone modifications in A549 cells treated with sodium propionate (SP) in time series for 24 hours. Color intensity scale bar represent the log2 fold difference of SP with control (blue, log2 fold-change < 0; white, log2 fold-change = 0; red, log2 fold-change > 0; black, NA). G) QIBC analysis of mean intensity levels of H3K4me3 in A549 cells treated with sodium propionate (SP, 5 mM) for 24 hours. Dots represent the cells and significance between the conditions was calculated using un-paired t-test. H) Schematic representation of the molecular mechanistic role of sodium propionate in epigenetic reprogramming to induce epithelial genes expression.

Motivated by the increased H3K27ac mark signifying active promoters and enhancers, a further investigation of other histone post-translational modifications (PTM) was carried out to understand the methylation and phosphorylation marks in the transcriptional activation programs initiated by SP. A549 cell line was treated with SP in a time-dependent manner (3 hours to 24 hours) followed by mass spectrometry based analysis of all histone marks related to methylation and phosphorylation. A high fold time-dependent increase in H3K4me1, H3K4me2 and H3K4me3 marks were identified with SP treatment (**Figure 7F**), while down-regulated histones marks include H3S10ph, H3K36me1, H1S1ph and H1T17ph (**Supplementary Figure 17E**). H3K4me marks represent transcriptionally active chromatin regulation with marks majorly found near active promoters, while H3S10ph is a mark of condensed chromatin during mitosis. H3K4me3 level was significantly found increased with SP by H3K4me3 mark staining by QIBC analysis (**Figure 7G**; **Supplementary Figure 17F-G**). In addition, *CDH1* gene and top up-regulated SP genes (*SCUBE1, ESPN* and *GPR4*) from RNA-Seq were analyzed for their peak enrichment in ChIP-Seq profile for H3K4me3 mark obtained from GEO (GSE35583) and the results showed increased peak enrichment for these genes (**Supplementary Figure 18**). Again, spheroid culture of A549 cells treated with TGFβ/TNFα showed a decrease in the histone marks of H3K4me1/2/3 and H3K27ac for the epithelial genes (*CDH1, KRT19*, and *OVOL2*) and increased mark levels for the *ZEB2* (**Supplementary Figure 19**), and thus validating the involvement of H3K4me1/2/3 and H3K27ac histone modifications in epithelial program modifications by propionate.

Collectively, our results indicate that propionate induces broad transcriptional initiation or activation marks in the regulation of epithelial gene expression encompassing H3K4me1/2/3 and H3K27ac marks. More importantly, E-cadherin expression by SP is mediated by p300 activation of H3K27ac mark at the promoter region of *CDH1* gene and shifting the cellular balance towards the epithelial state.

## Discussion

Deregulated metabolism is widely now recognized as a driving hallmark feature of cancer, and considerable attention has now been focussed on targeting oncogenic mediated metabolic processes involved in the aggressive state of lung cancer [54]. Notably, various lines of evidences highlight the metabolic rewiring as a hallmark of EMT in cancer, and modulating the metabolic state could inhibit or reverse the aggressive feature of EMT [20]. Further, making EMT and actionable therapeutic target could also have a profound implications in other EMT-induced pulmonary disorders including idiopathic pulmonary fibrosis and chronic obstructive pulmonary disease [55]. While several inhibitors are being surpassed to clinical trials, metabolic inhibitors still face major challenges [21–23] and realistic metabolic target inhibitors at the translational level are required. Therefore, in the current study, using lung cancer patients’ gene expression profiles as a solid source, a comprehensive screening of metabolic processes was performed to identify negatively associated metabolic processes with EMT using integrative functional genomics approach. Several positively and negatively associated metabolic processes with EMT were identified which were found concordant in multiple profiles of lung cancer performed using microarray or RNA-Seq technique showing that the pathway-centric analysis carried out here is technologically independent. Previously reported EMT-associated metabolic processes were also found to have both positive (chondriotin sulphate [39], heparan sulphate [40]), and negative (TCA, oxidative phosphorylation, and fatty acid synthesis [56]) connections in the current study. While short-chain fatty acids, propionate and butyrate, were the most negatively associated metabolic processes with EMT, several unreported roles of glycosaminoglycans such as dermatan sulphate or vitamin B5 metabolic processes were not explored deeply in the literature, and requires further investigation in the context of EMT in cancer.

Short-chain Fatty acids (SCFAs), produced by the gut microbiome with the breakdown of nondigestible fibres, have now attained considerable attention as dietary supplementation having beneficial and therapeutic potential including in cancer treatment [57,58]. Recent studies have highlighted the role of short-chain fatty acids in gluconeogenesis, energy metabolism, immune modulation, and maintenance of epithelial barrier integrity in gut, and few suggest the existence of gut-brain and gut-lung axis crosstalk in SCFAs effect [59–62]. SCFAs function either by directly involving in the core metabolism or in a receptor-mediated molecular process and as an epigenetic modulator [63]. Few studies showed the role of sodium butyrate and sodium propionate as a proliferation inhibitor in a breast cancer cell line, MCF-7 [64] or with the SP treatment at 10 mM in lung cancer cells (H1299 and H1703) [65]. However, we did not observe proliferative differences with propionate treatment at 5 mM in multiple lung cancer cell lines until 12 days. Both H1299 and H1703 are E-cadherin negative [66], and therefore a different mechanism could be involved with propionate in inhibiting the proliferation. In fact, upon testing H1299 with the treatment of SP, we could not detect any baseline E-cadherin expression in the cell line and so do any increase with SP (data not shown). Furthermore, gene signature enrichment analysis of RNA-Seq profiles from 24 hours, 3 days and 12 days treatment with SP did not show any significant down regulation of proliferative genes. Similarly, a previous report observed propionate’s role in pulmonary fibrosis by blocking LPS-induced EMT with the activation of PI3K/Akt/mTOR signaling pathway using BEAS-2B, a non-tumorigenic lung epithelial cell line, and *in vivo* by blocking inflammation and ECM deposition leading to pulmonary fibrosis [67].

*In vitro* treatment with sodium propionate showed a standalone EMT marker genes modulation with an increase in the key epithelial gene expression of E-Cadherin along with the repression of ZEB1 in multiple NSCLC cell lines. ZEB1 is a classical and master regulator of EMT process involved in several features of EMT such as in cancer stemness, invasion, metastasis and therapy resistance [12]. In fact, ZEB1-induced EMT contributes to early tumorigenesis, invasion and metastasis in non-small cell lung cancer [68]. SCFAs, mainly butyrate and propionate, have been found to increase the ZO-1 expression to restore and maintain the impaired barrier function of non-cancerous human bronchial airway epithelial cells [69]. In the current study, in addition to E-cadherin increase, we also observed membrane-associated increase in other epithelial-specific marker genes ZO-1 and EPCAM which plays a major role in the tight junction formation and cell adhesion, respectively. Also, we have seen a high-fold increase in the level of integrins from RNA-Seq analysis (**Supplementary Table 8**). Moreover, functional based cellular assay of propionate showed an enhanced epithelial property with an increase in the substrate adhesion. In addition, SP treated RNA-Seq profile showed a significant increased expression of the airway epithelial cell-type genes more specifically the pulmonary aleveolar type II cell-type markers implying the epithelial integrity could be achieved with propionate in the lungs. And single cell RNA sequencing of A549 cells showed that inherently parental A549 cells possess hybrid EMT modes of cells with both epithelial (cluster 3) and mesenchymal (cluster 4) featured cells. More interestingly, propionate specific up-regulated genes from RNA-seq were found enriched in the cluster 3 epithelial cells of scRNA analysis. This corroborates the identified propionate-specific epithelial program at the single cell level with the inherent A549 cells sequencing. On the other hand, we failed to observe any enrichment of overlap of SP down-regulated genes in cluster 4, and the genes had high overlap with cluster 2. However, identification of SP up-regulated genes in cluster 3 cells strongly suggests that SP is more of an epithelial enhancing factor and thereby controls or reduces the mesenchymal process.

Pre-treatment of cells with propionate possess the chemopreventive effect from TGFβ1-induced EMT state with decreased E-cadherin and increased ZEB1. Various chemopreventive drugs, either FDA approved or in phase trials, are now being identified with the metabolic block of EMT markers in several types of cancers such as with intake of prescribed statin medication prior to surgery for pancreatic ductal adenocarcinoma [70] or rolipram in lung cancer [71]. Importantly, propionate lost its ability to regain the cells from already transitioned mesenchymal cell state indicating that balancing or reinforcing effect of propionate towards the epithelial-oriented program is way more important (**Figure 2E**). Notably, our lung experimental metastasis model evidently showed that lung cancer cells treated with sodium propionate drastically possessed EMT inhibiting feature of metastasis or lung colonization ability. All these cellular, molecular and genomic analysis of propionate in the context of EMT implies that SP is involved more towards enhancing the lung-specific epithelial program thereby reducing the mesenchymal features. Therefore, lung cancer patients undergoing surgery or other treatment modes could be subjected for the short-chain fatty acids in dietary mode to have a chemopreventive effect.

Short-chain fatty acids are generally produced in the gut with the highest levels in the large intestine (30 to 150 mM), where bacterial fermentation of dietary fibre occurs to improve or maintain the host health, and it is then transported across the gut epithelium through liver into the systemic circulation (0.1 to 5 mM) [72]. Propionate and butyrate are in the concentration range of 1 to 15 μM in the systemic circulation [73]. On the other hand, there is no consensus range of SCFAs concentration in the lungs or about its substrate source. SCFAs have been detected in the sputum which has at 0.1 to 5 mM concentration possibly indicating the availability of SCFAs in the lungs, and suggesting an existence of gut-lung axis connection [61]. Moreover, while there are reports indicating that gut microbiota produced SCFAs diffuse into the blood to reach the lung, SCFAs are also found to be higher in the lungs compared to the blood in a study of people with HIV [74]. Treatment of primary human lung fibroblasts and airway smooth muscle cells with a supraphysiological concentrations (10-25 mM) of propionate or butyrate in combination with TNFα resulted in a pro-inflammatory effects of SCFAs [72]. The concentrations used in our study were based on the SCFA’s concentration observed in the human lungs [61,72]. Notably, our data on the propionate’s enhancement of epithelial features in the lung cells suggest potential possibility of specific production of propionate from lung-resident microbiome which would be of future investigation.

Despite the improvements in the molecular targeted therapies for lung cancer, around 80% of advanced NSCLC patients still receive standard first-line treatment of platinum-based chemotherapy such as cisplatin [46,75]. However, EMT is known to contribute to chemoresistance in many different cancer types including cisplatin-resistance in NSCLC [45,76]. Our *in vitro* study on the propionate’s role on inducing the chemosensitization of NSCLC cells is striking with the increase in γ-H2AX levels with cisplatin treatment. This is in line with a similar and recent observation of cisplatin sensitization with propionate through an increase in the H3 acetylation levels through HDAC inhibition [77]. While we observed an increase in the γ-H2AX levels with propionate treatment alone, we did not observe any changes in the DNA breaks or repair with SP on its own as assessed from RAD51 or TP53BP1 expression, sensitive markers of homologues or non-homologues repair mechanism, respectively. We postulate that propionate could increase the genome accessibility to cisplatin’s action of DNA break or damage which would be of future investigation in the clinical aspect.

Dysregulation of propionate metabolism was observed to play an anaplerotic reaction in the accumulation of methylmalonic acid (MMA) resulting in breast and lung cancer with an increased metastatic potential [78], and accumulation of MMA confers poor prognosis in aged patients to contribute for metastasis [79]. However, the current study did not observe any propionate-specific metabolic perturbations while propionate treatment epigenetically reinforced the epithelial identity. This implies a dual-role of propionate in EMT and a potential feedback loop process controlling the EMT plasticity which would be of further exploration.

Molecular mechanistic investigation of propionate’s role in inducing epithelial-specific program found a receptor-independent role of propionate in modulating the expression of epithelial-genes through epigenetic mechanism. Evidences showed that HDAC inhibitors (HDACi) can increase the E-cadherin levels in the cells treated with various HDACi such as vorinostat and mocetinostat [80]. In fact, we also observed that SP treated RNA-seq profile showed similarity metrics with HDACi, and that these inhibitors (vorinostat, mocetinostat and MI192) on its own can induce the E-cadherin expression. However, treatment with HAT inhibitors (HAT inhibitor VII and A-485) showed convincing down-regulation of E-cadherin levels along with SP. HAT inhibitor VII is a mixed mode selective inhibitor of p300/CBP and PCAF histone acetyltransferases which blocks the autoacetylation of p300 and PCAF, while A-485 is a more selective potent inhibitor of p300/CBP catalytic site involved in the p300 autoacetylation. A-485 does not inhibit other acetyltransferase such as PCAF, GCN5L2, MYST3/4, HAT1, MOF and TIP60 [53]. In addition, our ChIP-Seq analysis of H3K27ac with SP treatment substantially showed that *CDH1* gene is increased with the occupancy of the histone activation mark. All these could evidently point to the possibility that SP regulate the E-cadherin expression through p300 signaling event with H3K27ac mark increase at the *CDH1* locus. Recently, it has been shown that propionate and butyrate metabolized to their corresponding acyl-coAs can activate p300 auto-acylation for histone modification events including H3K27ac in HEK293, HepG2 and MCF7 [52]. A receptor-independent effect of propionate and butyrate, but not acetate, was also observed albeit in allergic airway inflammation [81]. In an attempt to deeply understand the epigenetic roles of propionate, we further investigated all other possible modes of epigenetic modifiers (histone methyltransferase and DNA methyltransferase). While we did not observe any changes in the levels of DNA methyltransferase by propionate in A549 cells upon treatment with 5-Aza (**Supplementary Figure 16F**), we found a drastic involvement of histone-associated methyltransferase. For the first time, we observed propionate to have a dual ability to induce H3K27ac and H3K4me levels. H3K27ac is an active promoter and enhancer mark, whereas H3K4me3 is a mark of transcriptionally active chromatin and found around the transcriptional start site of active genes [82]. In the EMT context, it has been shown that Snai1 mediates histone demethylase LSD1 to demethylate H3K4me2 at E-cadherin and other epithelial gene promoter sites to transcriptionally repress epithelial genes to promote EMT [83]. Similarly, loss of H3K27ac mark leads to the repression of epithelial identity genes mediated by MNT, a transcriptional regulator, during EMT process [84]. All these histone mark dynamicity is in accordance with our observations showing that SP induces H3K4 methylation or H3K27 acetylation levels with the occupancy of marks at the epithelial genes expression site to enhance the epithelial identity thereby reducing promotion of mesenchymal features. From our study, we also identified the down-regulation of H3K36me1, H3S10ph, H1S1ph and H1T17ph whose role in the context of EMT promotion would be future investigation.

Besides the genetic and epigenetic alterations observed in the transformation of normal to abnormally proliferative cancerous cells, altered gut microbiome are also known to have a profound role in the tumorigenesis through changes in the metabolic processes [85,86]. EMT has been shown as an important factor in early steps of lung tumorigenesis as observed with the oncogenic transformed human bronchial epithelial cells [10] and with the cigarette smoke-induced epigenetic alterations persuading the EMT process [11]. In fact, our KPL mouse model administered orally with sodium propionate in drinking water showed significant attenuation of the aggressive state of lung tumorigenesis as noticed with the repression of metastatic spread to lymph nodes. Again, KPL cell line derived from the lung tumorigenic mouse showed that the cells are responsive to propionate in increasing the epithelial gene marker E-cadherin. Recently, gut microbiota metabolomics profiling of NSCLC patients undergoing immunotherapy with nivolumab showed that long term responders were majorly characterized with the presence of SCFA metabolites (propionic, butyric, acetic and valeric acids) with beneficial effects [88]. Therefore, propionate could be used as a combinatorial drug along with immunotherapy for NSCLC patients showing improved therapeutic efficacy [89]. Interestingly, SCFAs levels in the body can be modulated either by prebiotic or postbiotic or combination of both [74], and propionate is a safe food ingredient [43]. Thus, propionate being a natural dietary and symbiotically produced compound has the potential to inhibit metastasis, and increase chemosensitivity of the drugs, while enhancing the lung epithelial identify providing beneficial effects which in turn warrants further clinical level exploration for NSCLC patients.

## Conclusion

The present study identified the therapeutic ability of SCFA propionate in inhibiting EMT process and its associated features for lung cancer. Further, propionate mediates epigenetic reprogramming with an increase in the transcriptionally active chromatin marks to reinforce epithelial gene expression pattern in facilitating lung-specific epithelial identity with potential clinical benefits.

## Abbreviations

TCGA: The Cancer Genome Atlas
GEO: Gene Expression Omnibus
EMT: Epithelial-to-mesenchymal transition
NSCLC: Non-small-cell lung cancer
ECM: Extracellular matrix
CDDP: cis-diamminedichloroplatinum (II)
SCFA: Short-chain Fatty Acids
SP: Sodium propionate
SB: Sodium butyrate
SA: Sodium acetate
AAV: Adeno-associated virus
HAT: Histone Acetyl Transferase
HDAC: Histone deacetylase

## Conflict of Interests

The authors declare no conflict of interest.

## Author’s contribution

Conceived the concept: VR and PC; Designed the experiments: VR, RS, KS, ONJ and PC; Performed the experiments: VR, PNG, LP, STJ, ELB, MAS and KS. Performed computational and statistical analysis: VR, STJ, ELB, LP, PNG and PC; Evaluated H&E staining: KEO; Wrote the manuscript: VR and PC. All the authors read and approved the final manuscript.

## Acknowledgements

This work was supported by the Interdisciplinary Center for Clinical Research of the University of Erlangen-Nuremberg, the German Research Foundation (DFG, CE 281/6-1), the Novo Nordisk Foundation (Hallas-Møller Ascending Investigator Grant 0066909), and by the Danish Cancer Society (A18859). Sequencing was performed at the Center for Functional Genomics and Tissue Plasticity, Functional Genomics & Metabolism Research Unit, University of Southern Denmark. The authors thank Tenna P. Mortensen, Maibrith Wishoff and Ronni Nielsen for sequencing assistance. The authors also thank Danish Molecular Biomedical Imaging Center (DaMBIC), University of Southern Denmark, Odense, for providing access to use fluorescent microscope. Results have been partially presented at the X TEMTIA meeting in Paris, France.

## Supplementary Figure Legends

**Supplementary Figure 1. Expression pattern of pan-cancer EMT gene signature in genetically-induced or TGFβ1-treated cell line gene expression profiles.** A-C) Heatmap represents the gene expression pattern of pan-cancer EMT gene signature showing an increase in the mesenchymal genes with a decrease in the epithelial genes in the TGF-β1 treated cells. Gene expression profile of non-small cell lung carcinoma cell lines (NCI-H358, HCC827 and A549) treated with TGF-β1 for 3 weeks were obtained from GEO (GSE49644). D-H) Heatmap represents the gene expression pattern of pan-cancer EMT gene signature showing an increase in the mesenchymal genes with a decrease in the epithelial genes in the EMT induced conditions. Gene expression profile of retrovirally transduced EMT-inducing genes (TGFb1, SNAIL, TWIST and GOSSECOID) and knock down of E-cadherin in immortalized HMLE breast epithelial cells were obtained from GEO (GSE24202).

**Supplementary Figure 2. Integrative genomic analysis of EMT associated REACTOME related metabolic processes.** Network visualization of REACTOME metabolic processes positively and negatively associated with EMT (meta-r>0.3 and meta-r<-0.3; meta-p<0.05). z-score based activation analysis of pan-cancer EMT gene signature and REACTOME gene-set metabolic processes was performed in 1476 gene expression profiles from 6 datasets of lung cancer patient samples followed by meta-correlation analysis with the activation scores.

**Supplementary Figure 3. Validation by independent approaches of the negative association between the metabolic processes of SCFAs with EMT gene signature.** A-B) Correlation plot of short-chain fatty acids (propionate or butanoate) with pan-cancer EMT gene signature in lung adenocarcinoma patients (N=510) from The Cancer Genome Atlas (TCGA). C-F) Correlation plot of activation scores of short-chain fatty acids (propionate and butanoate) with different source of EMT gene signatures from Gotzmann (C-D) or Jechlinger (E-F) in lung cancer patients samples obtained from GEO (GSE72094, N=442). G) Gene-set enrichment analysis of Hallmark EMT gene-sets with the lung cancer patient samples (GSE72094; N=442) categorized as low and high based on the butanoate gene-set activation levels shows an enrichment of EMT in low butanoate patient samples. Ranking of genes with signal2noise metric was used for GSEA.

**Supplementary Figure 4. Survival analysis shows good prognosis for propionate or butanoate in lung cancer patient samples.** A-F) Overall and relapse free survival analysis in the indicated datasets of lung cancer patient samples categorized as low and high propionate (A-C) or butanoate (D-F) levels based on the median showed good prognosis for propionate or butanoate gene-set. p-value was calculated using log-rank method.

**Supplementary Figure 5. *In vitro* treatment effect of SCFAs, propionate or butanoate, in EMT marker gene expression.** A) Western blot analysis of E-cadherin and ZEB1 in A549 cell line treated with equimolar concentration of sodium chloride or sodium propionate in a time-dependent manner. β-Actin was used as an internal control. B) Western blot analysis of E-cadherin and ZEB1 in SKMES1 cells treated with sodium butanoate in a dose- and time-dependent manner. β-Actin was used as an internal control. C) Western blot analysis of E-cadherin and ZEB1 in NCI-H23 cell line treated with 1 mM of sodium propionate for 24 hours. β-Actin was used as an internal control. D-E) Western blot analysis of E-cadherin and ZEB1 in PANC1 cells treated with sodium propionate (D) or sodium butanoate (E) in a dose- and time-dependent manner. β-Actin was used as an internal control. F) Immunofluorescence staining of E-cadherin in PANC1 cell line treated with sodium propionate (5 mM) for 3 days. DAPI was used as a nuclear stain. G) Flow cytometry analysis of membrane-associated E-cadherin (PE-conjugated E-cadherin) in A549 cells treated with sodium propionate at 5 mM concentration for 48 hours. Mean fluorescence intensity of PE-E-cadherin in SP treated cells and control were 62858 and 21999, respectively. PE-conjugated IgG1 was used as a control for flow cytometry. H) Western blot analysis of E-cadherin and ZEB1 in A549 cells treated with TGF-β1 (2 ng/ml) for 48 hours followed by treatment with sodium propionate (5 mM) for 24 hours. β-Actin was used as an internal control. I) Western blot analysis of E-cadherin and ZEB1 in A549 cells treated with TGF-β1 (2 ng/ml) for 48 hours followed by treatment with sodium butanoate (1 mM) for 24 hours. β-Actin was used as an internal control. J) Western blot analysis of E-cadherin and ZEB1 in A549 cells pre-treated with sodium butanoate (1 mM) for 48 hours followed by TGF-β1 (2 ng/ml) for 24 hours. β-Actin was used as an internal control. ns – non-significant.

**Supplementary Figure 6. Effect of sodium propionate on lung tumorigenesis in KPL virus intubated C57BL/6 mice administered orally with sodium propionate in drinking water.** A) Bar plot of consumption of drinking water containing sodium propionate (150 mM) or sodium chloride (150 mM) by C57BL/6 mice over the course of 5 weeks. Sodium propionate (150 mM) or sodium chloride (150 mM) was freshly prepared in drinking water by changing the water twice a week for mice. Significance was calculated using Two-way ANOVA. B) Line plot of body weight measurements over the course of study of C57BL/6 mice with lung tumorigenesis induced with AAV-KPL virus. Sodium propionate (150 mM) or sodium chloride (150 mM) was administered orally in drinking water. C) H&E staining of lung and axial lymph nodes of C57BL/6 with lung tumorigenesis induced with AAV-KPL virus and orally administered with sodium chloride (MAS53 to MAS59) or sodium propionate (MAS60 to MAS66) in drinking water. D) Comparison of number of tumor lesions from C57BL/6 mice with lung tumorigenesis induced with AAV-KPL virus. Sodium propionate (150 mM) or sodium chloride (150 mM) was administered orally in drinking water. Data points represent mean ± SEM. Significance was calculated using unpaired t-test with alpha value set at 0.05. ns – non-significant.

**Supplementary Figure 7. SCFAs propionate or butanoate sensitizes the cells to cisplatin treatment in NSCLC cell lines.** A-B) Dose responsive curve of cisplatin treatment in combination with sodium butanoate (SB, 5 mM) in A549 (A) or SKMES1 (B) cell lines. Data points are represented as mean ± SD with p-value calculated from Two-way ANOVA analysis. C) Quantification of dead cells as green object count using Cytotox Green in A549 cells treated with dose-dependent cisplatin in combination with 5 mM SB. Data points are represented as mean ± SD. D) Line plot comparison of the green object count for the cells treated with cisplatin (2.5 μM) in combination with SP (5 mM) or SB (1 mM). Significance was calculated from Two-way ANOVA.

**Supplementary Figure 8. SCFAs propionate sensitizes the cells to cisplatin treatment without inducing DNA damages.** A) QIBC analysis of RAD51 foci levels in A549 cells treated in the indicated dose-dependent concentrations of cisplatin for around 6 hours in combination with SP pre-treatment (5 mM) for 24 hours. B) QIBC analysis of sum intensity levels of RAD51 foci in A549 cells plotted as bar plot showing no much difference between treatment of cisplatin in the indicated dose-dependent concentrations in combination with SP (5 mM). C) QIBC analysis of sum intensity levels of TP53BP1 foci spots in A549 cells showing no much difference between the control and SP (5 mM) treatment. D) Bar plot representation of sum intensity levels of TP53BP1 foci spots indicating no changes in the levels between the conditions. E) QIBC analysis of cell cycle phases segregated based on Edu staining in A549 cells treated with sodium propionate (5 mM) for 24 hours. F) Stacked bar plot represents the percent distribution levels of cell cycle phases (G1, S and G2) in A549 cell lines treated with sodium propionate (5 mM) for 24 hours. ns – non-significant.

**Supplementary Figure 9. RNA-Seq expression profiling of SP treated samples from 3 days and 12 days show similar gene expression pattern.** A-B) Real time quantitative PCR analysis of E-cadherin (A) and ZEB1 (B) in A549 cell lines treated with sodium propionate at 5 mM in time series. Dotted line represents the fold change cut-off at 1.5 (A) or at 0.67 (B). Significance was calculated using un-paired t-test. C) Unsupervised hierarchical clustering analysis of RNA-seq samples of A549 cell line treated with SP for 3 days and 12 days along with control. D-E) MA plot of differentially expressed genes identified in SP treated 3 days (D) and 12 days (E) compared to the control. Blue dots represent the significantly differentially expressed genes with a fold change of log2 (1). F-G) Venn diagram representation of overlap analysis between SP 3 days and SP 12 days up-regulated (F) or down-regulated (G) genes. ****- p-value <0.0001; *** - p-value <0.001, ns – non-significant.

**Supplementary Figure 10. GSEA analysis of Hallmark apical surface and Hallmark apical junction enrichment in 3 days and 12 days SP treated expression profiles.** A-C) GSEA of Hallmark apical surface gene-set enrichment in SP 3 days samples compared to the control (A), and enrichment of Hallmark Apical Junction gene-set (B) or Hallmark Apical Surface (C) in SP 12 days samples compared to the control. Ranking of genes based on signal2noise metric was used for GSEA.

**Supplementary Figure 11. Single cell RNA-Seq of parental A549 cell line shows cell clusters of epithelial and mesenchymal cell type enrichment.** A) Table list the number of cells identified in each cluster of parental A549 cells sequenced by single cell RNA sequencing. B-H) Enrichr based gene enrichment analysis of marker genes of cell clusters 1 to 4 identified from scRNA-seq of A549 cells for MSigDB gene-sets collection (B-E), and PanglaoDB (F-H) gene-sets collection. Significant enrichment was considered with p-value < 0.001. I-J) Venn diagram representation of overlap analysis between Hallmark EMT gene signature and cluster 1 marker genes (I) or cluster 4 marker genes (J). Overlapping genes are listed below the respective venn diagrams.

**Supplementary Figure 12. Validation of SP down-regulated gene-set enrichment in genetically-induced EMT state in HMLE cell line gene expression profiles.** A-D) Gene-set enrichment analysis of SP 12 days down-regulated gene-set in TGFβ1-, TWIST-, GOOSECOID- and SNAIL-induced HMLE cell line compared to the control obtained from GEO (GSE24202). Ranking of genes with signal2noise metric was used for GSEA.

**Supplementary Figure 13. Validation of lung-specific epithelial gene-set expression in RNA-Seq expression profile of A549 cell lines treated with SP for 3 days and 12 days.** A-D) Box plot visualization of the enriched markers of lung epithelial cell-type gene-set (A), pulmonary alveolar type II cell-type gene-set (B), establishment of epithelial cell polarity gene-set (C), and Hallmark apical surface gene-set (D) in control, SP 3 days and SP 12 days RNA-Seq samples of A549 cell line. Significance was calculated using un-paired t-test between the SP treated cells and the control. All the lung epithelial cell-type associated gene-sets were collected from PanglaoDB (A-C) and hallmark apical surface gene-set from MSigDB (D). E) Overall survival analysis in lung cancer patient samples (GSE72094; N=442) categorized as low- and high-propionate levels of SP 12 days gene-set z-score activity based on the median showed good prognosis. p-value was calculated using log-rank method. **** - p-value <0.0001; *** - p-value <0.001; ** - p-value <0.01; * - p-value <0.05

**Supplementary Figure 14. Identification of HDAC inhibitors based on similarity search analysis of SP gene-set expression pattern in L1000CDS^2^ search engine.** A) Bar plot indicates the top ranked drug gene signatures similar to SP 12 days gene-set based on the similarity score identified from L1000CDS^2^ web tool. Identical drugs from different treatment conditions are colored the same. B) Western blot analysis of E-cadherin and ZEB1 in A549 cells treated with HDAC inhibitor, mocetinostat in the indicated dose- and time-dependent manner in combination with SP 5 mM. β-Actin was used as an internal control. C) Western blot analysis of E-cadherin and ZEB1 in A549 cells treated with HDAC inhibitor, MI192 in the indicated dose- and time-dependent manner in combination with SP 5 mM. β-Actin was used as an internal control.

**Supplementary Figure 15. Identification of possible mechanistic role of propionate’s action in E-cadherin’s increased expression.** A) Schematic representation of possible modes of propionate’s mechanism of action in the cell involving MCT transporters, epigenetic mechanism, metabolic pathway, and GPR41 signaling. B) Schematic representation of propionate metabolic pathways with the enzymes involved in the metabolic conversions. Green highlighted enzymatic genes were used for siRNA-mediated knockdown study. C) Western blot analysis of E-cadherin and ZEB1 in A549 cells knock down with propionate-specific metabolic genes (*PCCB, MCEE, MUT, HIBCH* and *ALDH6A1*) for 48 hours followed by treatment with sodium propionate for 24 hours. Non-targeting siRNA was used as a control. TUBA4A was used as an internal control. D-E) Western blot analysis of E-cadherin and ZEB1 in A549 cells treated with MCT transport inhibitors, AZD3965 (D) and Syrosingopine (E), in the indicated dose-dependent concentrations in combination with sodium propionate (SP, 5 mM) for the indicated time points. β-Actin was used as an internal control. F) Western blot analysis of E-cadherin and ZEB1 in A549 cells treated with GPR41 modulator, AR420626 (10 μM) in combination with sodium propionate (SP, 5 mM) for the indicated time points. β-Actin was used as an internal control. G-H) Western blot analysis of E-cadherin and ZEB1 in A549 cell line treated with PPARγ inhibitor, GW9662 (G) and PPARγ activator, Troglitazone (H), in the indicated concentration in combination with sodium propionate (SP, 5 mM) for the indicated time points. β-Actin was used as an internal control.

**Supplementary Figure 16. Evaluation of H3K27 acetylation increase in the regulation of *CDH1* gene expression.** A) Western blot analysis of p300 acetylation, E-cadherin, ZEB1 and H3K27ac in SKMES1 cells treated with A-485 (10 μM) in combination with sodium propionate (SP, 5 mM) for 24 hours. β-Actin was used as an internal control. B-C) Western blot analysis of E-cadherin and ZEB1 in A549 cells treated with H3K27me3 inhibitors, GSK126 (B) and GSK343 (C), in the indicated dose-dependent concentrations in combination with sodium propionate (SP, 5 mM) for the indicated time points. β-Actin was used as an internal control. D-E) Bar plot of mean intensity levels of H3K27ac (D) and HP1 (E) in A549 cells treated with A-485 (10 μM) in combination with sodium propionate (5 mM) for 24 hours. Significance was calculated from One-way ANOVA with post-hoc analysis of un-paired t-test. F) Western blot analysis of E-cadherin and ZEB1 in A549 cells treated with 5-Aza-2’-deoxycytidine and sodium propionate in the indicated concentrations for 5 days. β-Actin was used as an internal control. ****- p-value <0.0001.

**Supplementary Figure 17. Propionate induces transcriptional histone active marks during epithelial gene expression program.** A) Western blot analysis of E-cadherin, ZEB1, H3K27ac and H3 in A549 cells treated with sodium propionate (5 mM) for 3 hours and 24 hours. β-Actin was used as an internal control. B) Principal component analysis of RNA-seq profiles of A549 cells treated with sodium propionate at 5 mM for 3 hours and 24 hours. C) Enrichr analysis of SP 24 hours up-regulated genes identified from RNA-Seq. D) GSEA analysis of Hallmark EMT gene-set enrichment analysis in a lung cancer gene expression profile (GSE72094, N=442) as a continuous label of SP 24 hours down-regulated gene-set z-score activity. Ranking of genes was based on Pearson’s correlation metric. E) Heatmap representation of histone post-translational modifications in A549 cells treated with sodium propionate at 5 mM in time series for 24 hours. Red highlighted box indicates the increased histone modifications in SP treated conditions compared to the control. Blue highlighted box indicates the decreased histone modifications in SP treated conditions compared to the control. F) Representative images from QIBC analysis of A549 cells treated with SP (5 mM) for 24 hours and stained with H3K4me3 antibody. Scale bar represents 50 μm. G) Bar plot of mean intensity level of H3K4me3 in A549 cells treated with SP (5 mM) for 24 hours. Significance was calculated from un-paired t-test. ****- p-value <0.0001.

**Supplementary Figure 18. Peak enrichment analysis of H3K4me3 marks in *CDH1* gene and top up-regulated genes from SP treated RNA-Seq profile.** Integrative Genomic viewer screen shot visualization of H3K4me3 in A549 cells for the peak enrichment at *CDH1, SCUBE1, ESPN* and *GPR4* locus identified from ChIP-Seq for H3K4me3 in A549 cells obtained from GEO (GSE35583).

**Supplementary Figure 19. Peak enrichment analysis of H3K4me and H3K27ac marks in epithelial and mesenchymal genes upon EMT induction.** Integrative Genomic viewer screen shot visualization of H3K4me1, H3K4me2, H3K4me3 and H3K27ac in A549 cells for the peak enrichment at *CDH1, KRT19, OVOL2*, and *ZEB2* locus identified from ChIP-Seq in A549 spheroid culture treated with TGFβ or TNFα obtained from GEO (GSE42374).

## Notes

### Competing Interest Statement

The authors have declared no competing interest.

