## Supplementary Files for "Propionate reinforces epithelial identity and reduces aggressiveness of non-small cell lung carcinoma via chromatin remodelling"

Supplementary Figure 1

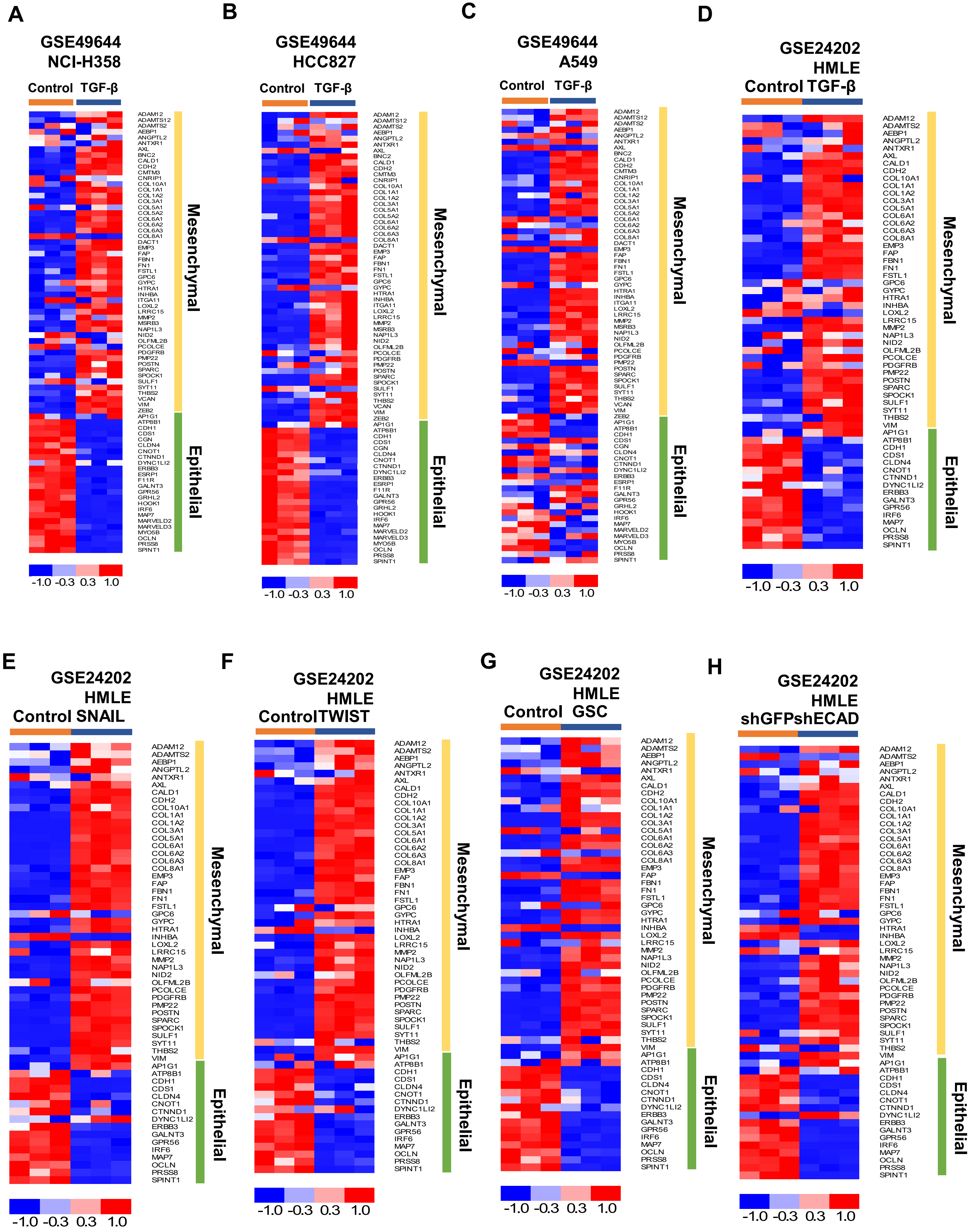

#### Supplementary Figure 2

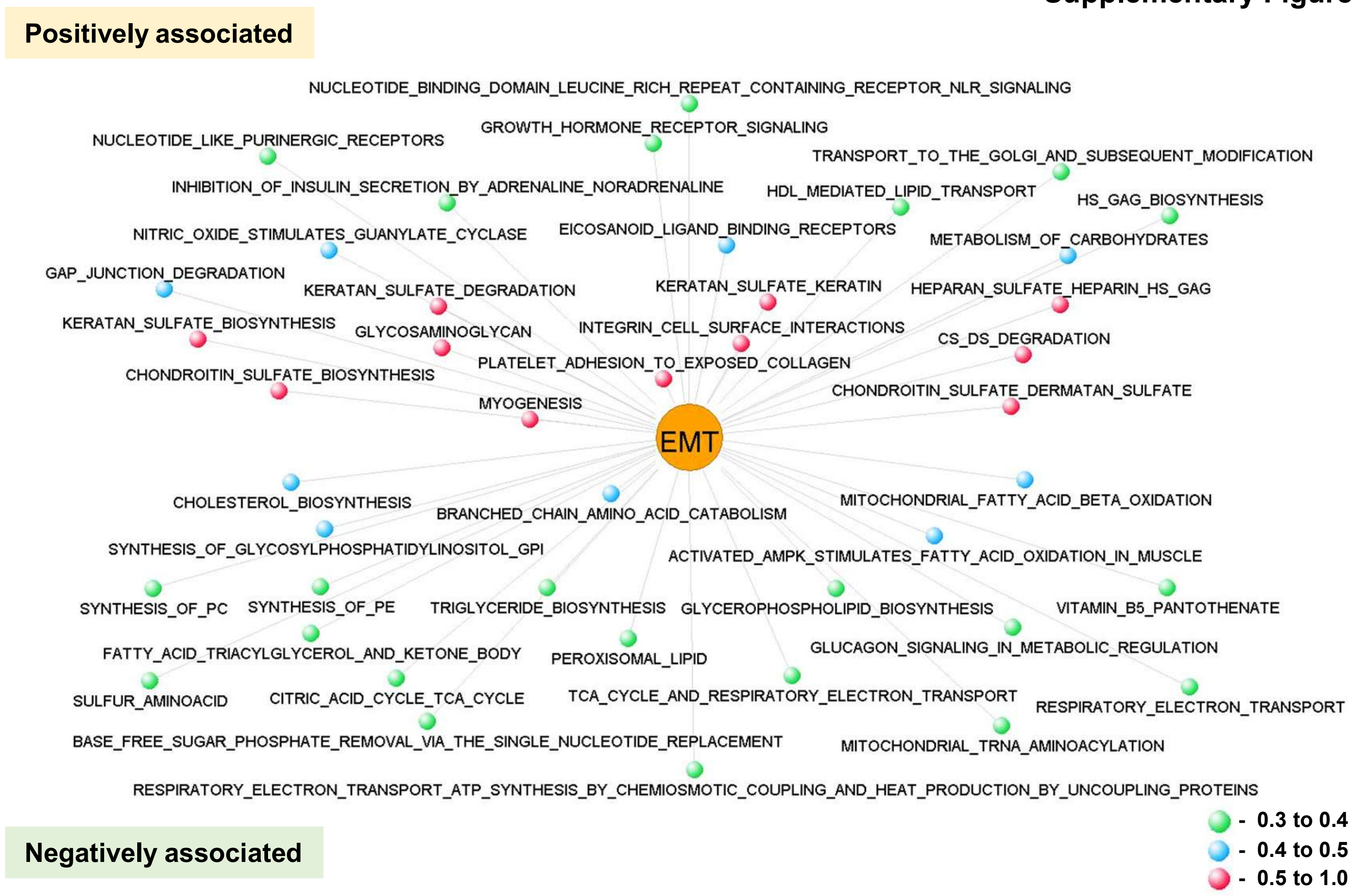

**Supplementary Figure 3**

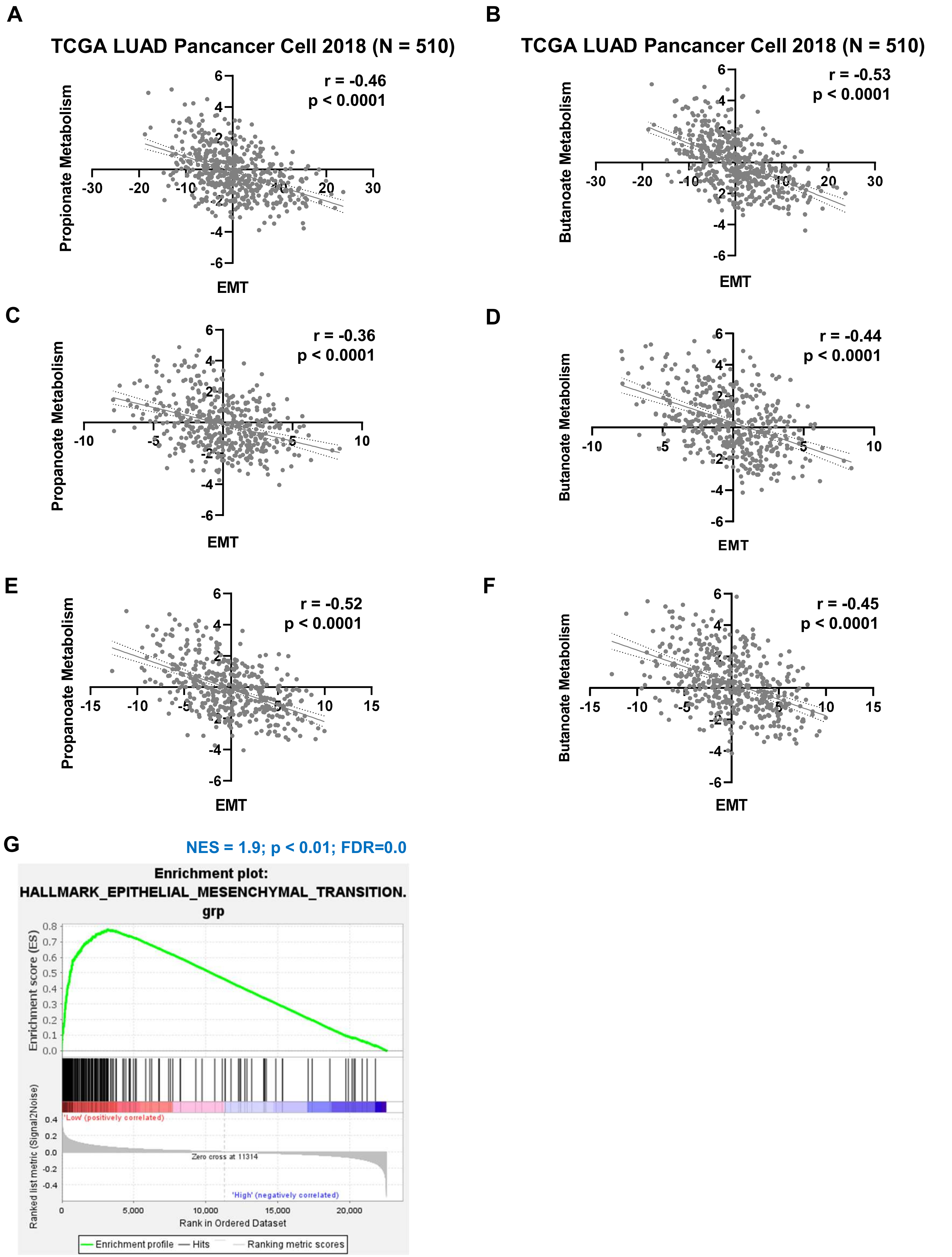

**Supplementary Figure 4**

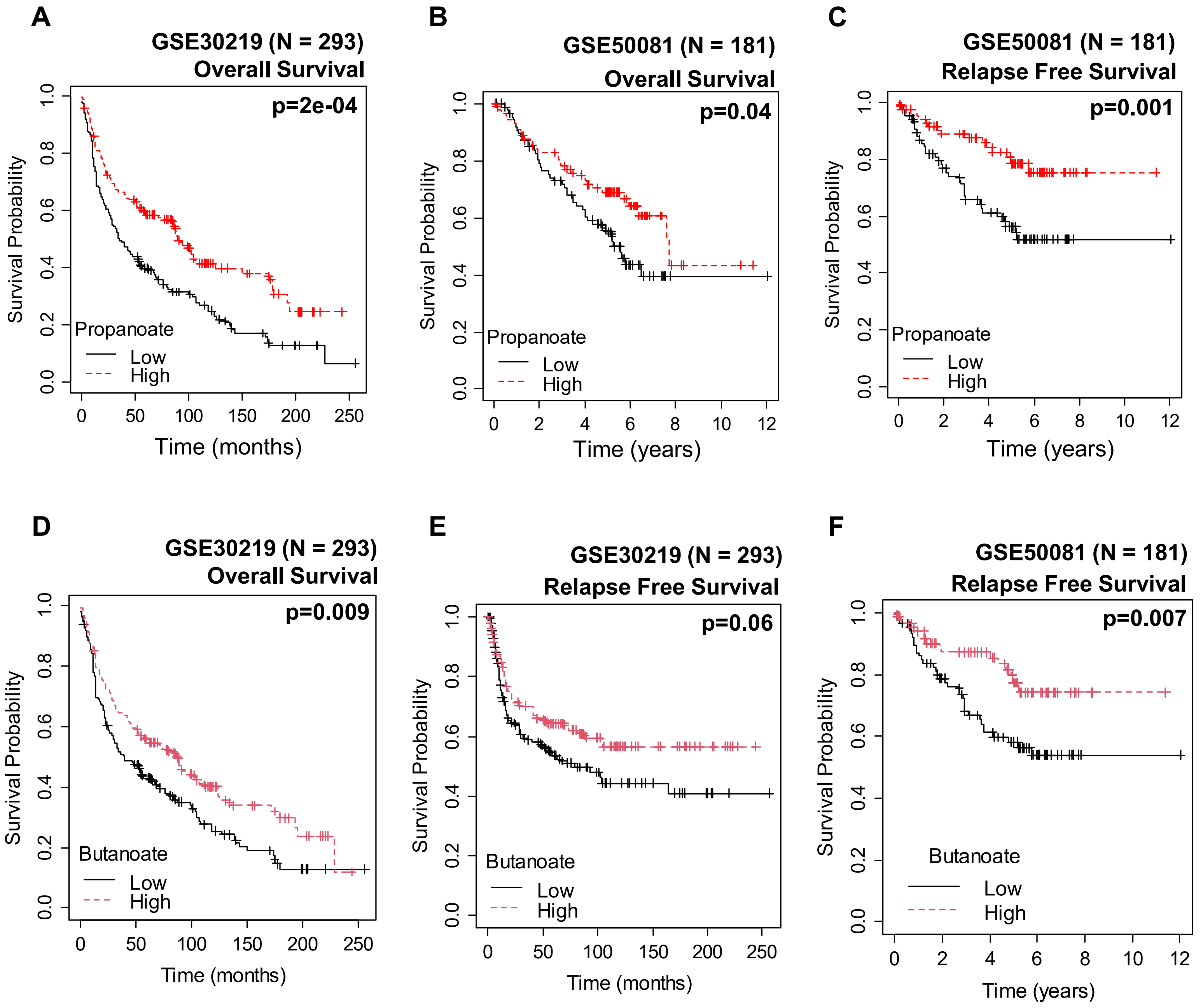

#### Supplementary Figure 5

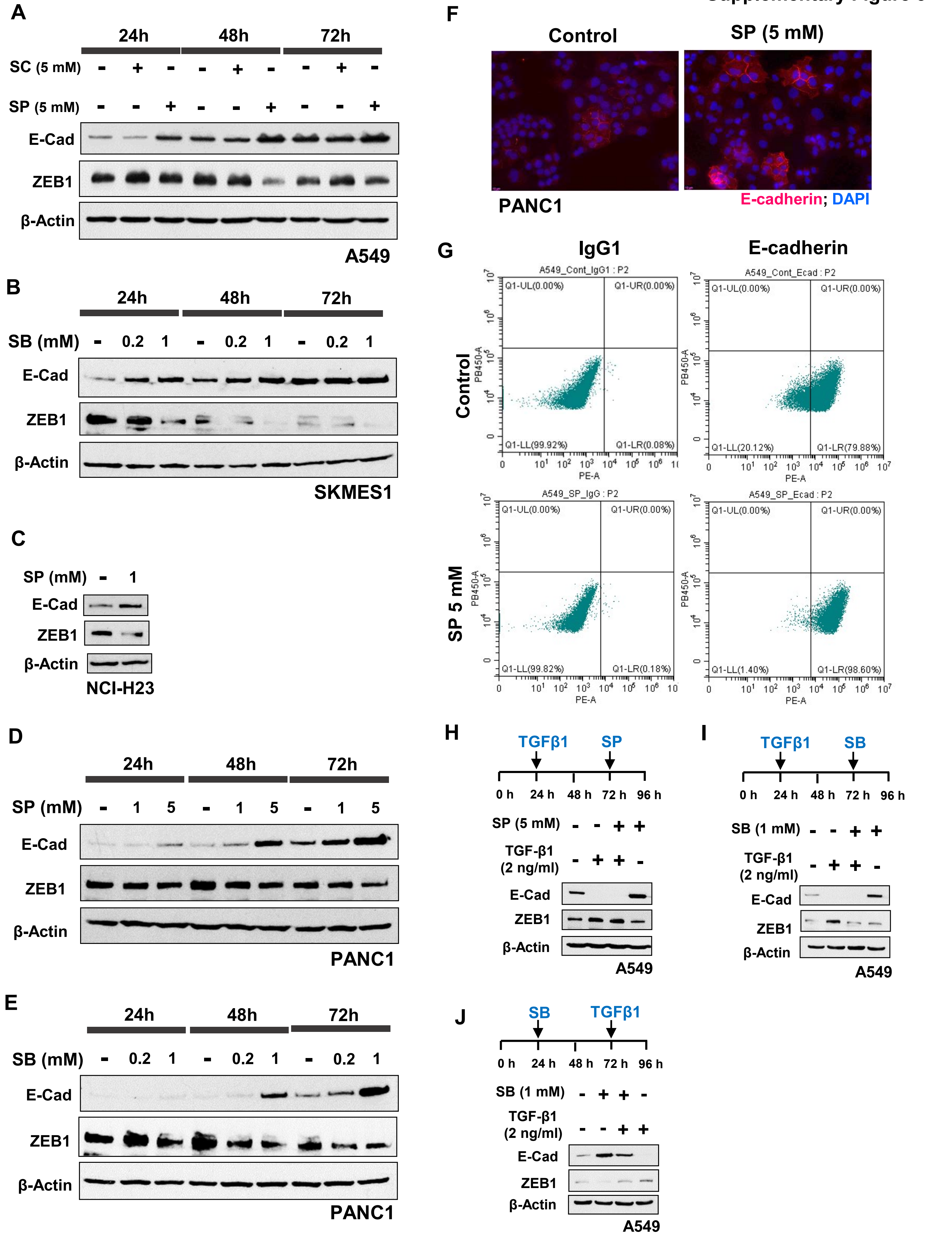

#### Supplementary Figure 6

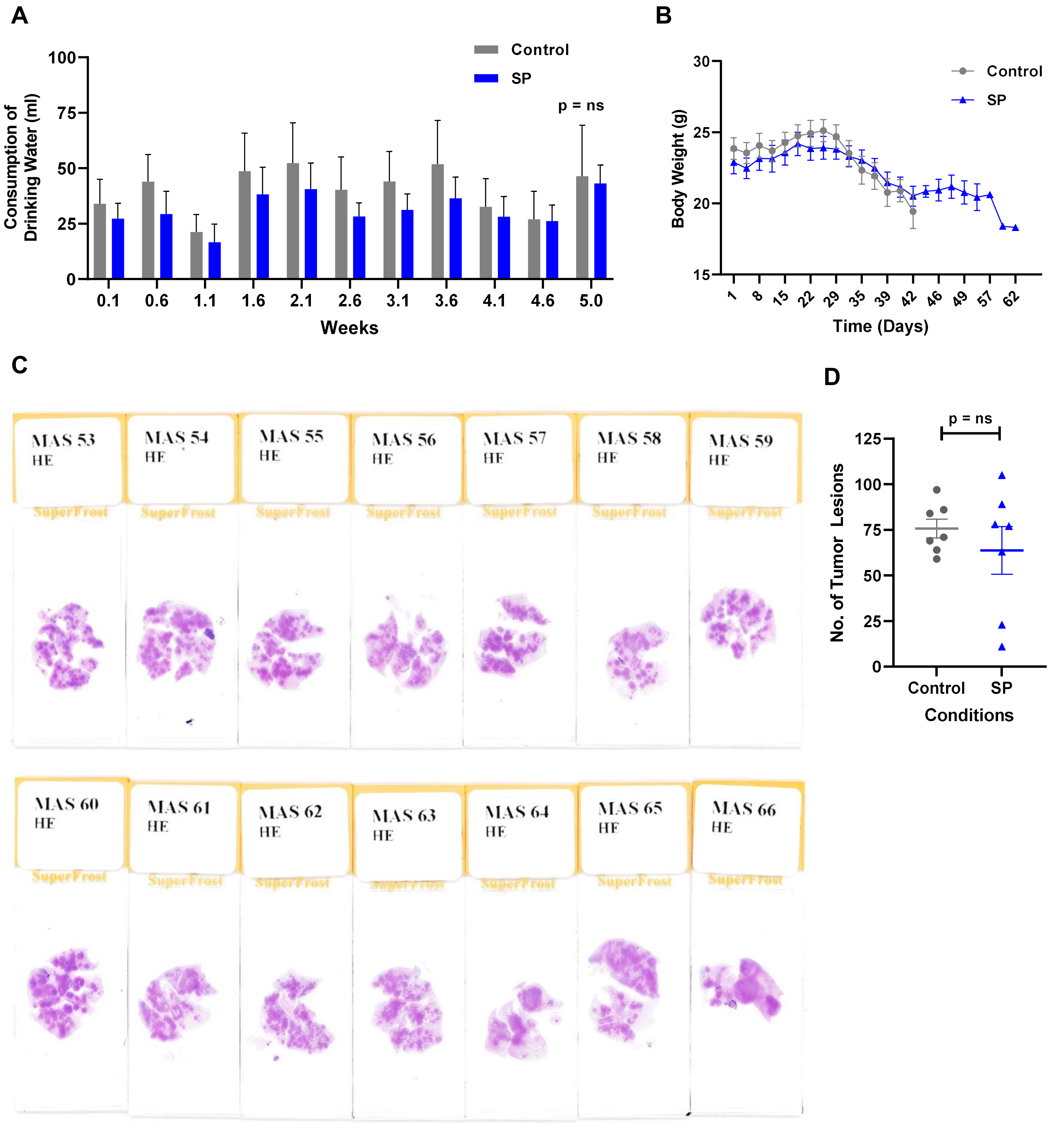

#### Supplementary Figure 7

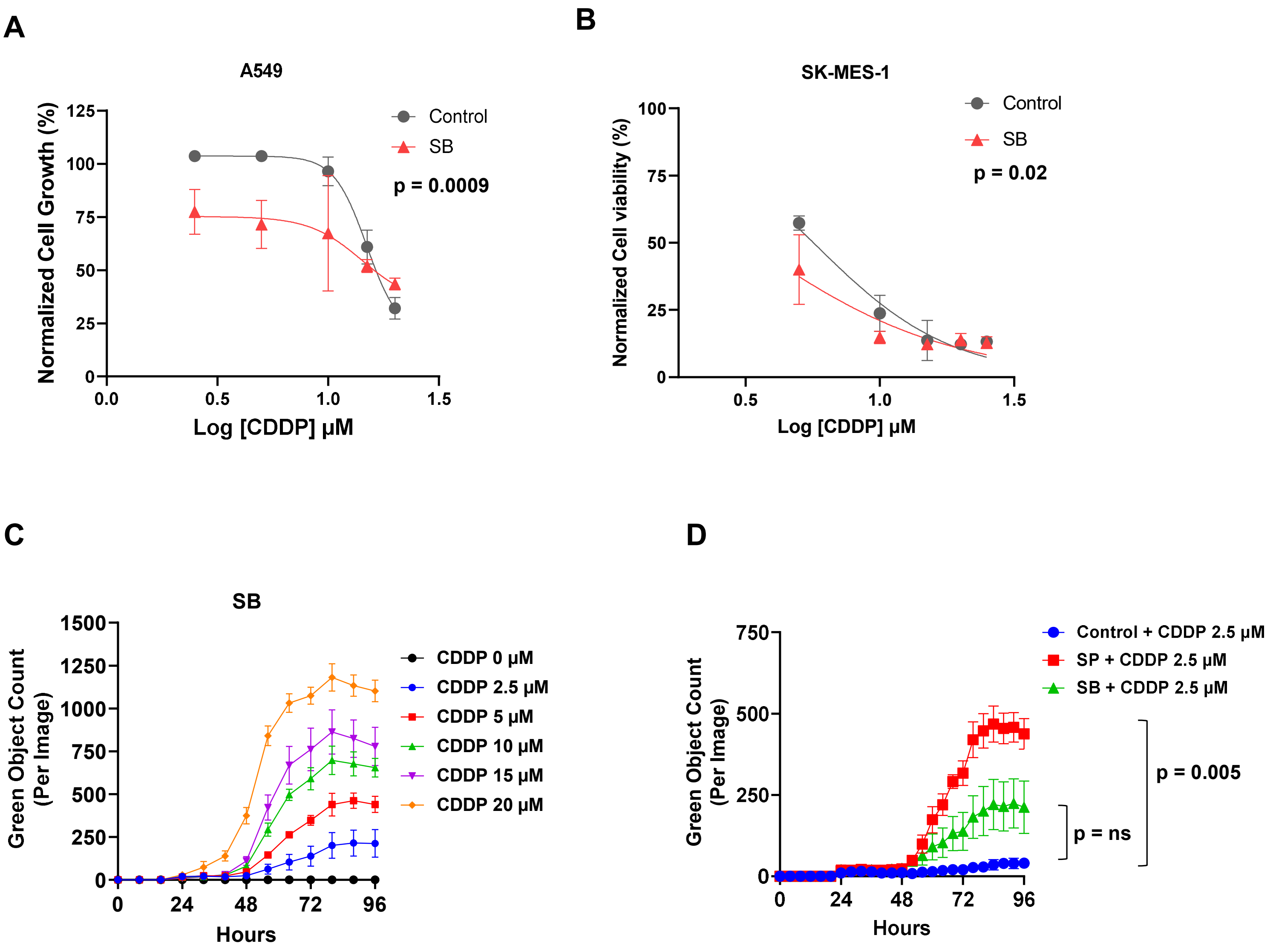

Supplementary Figure 8

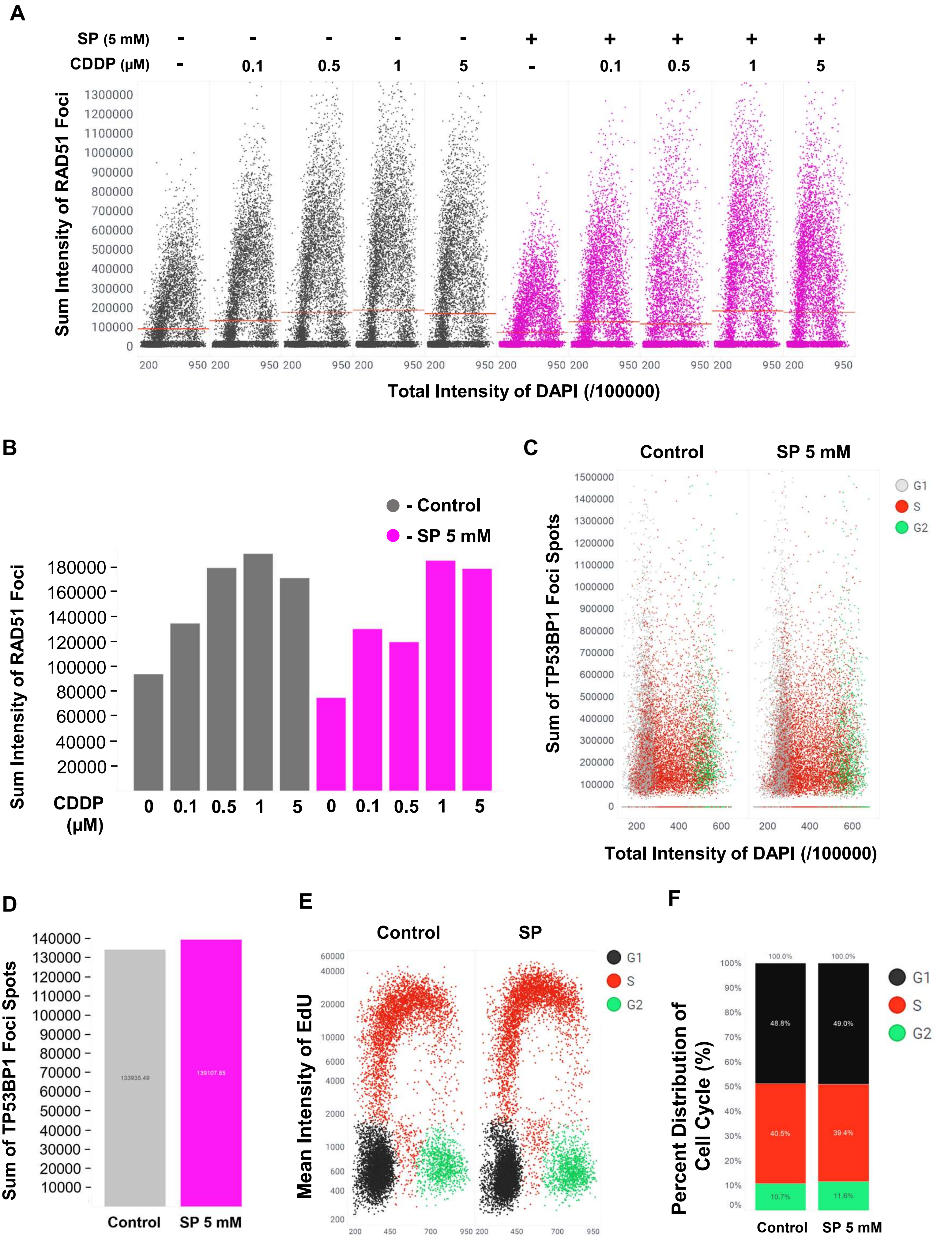

#### Supplementary Figure 9

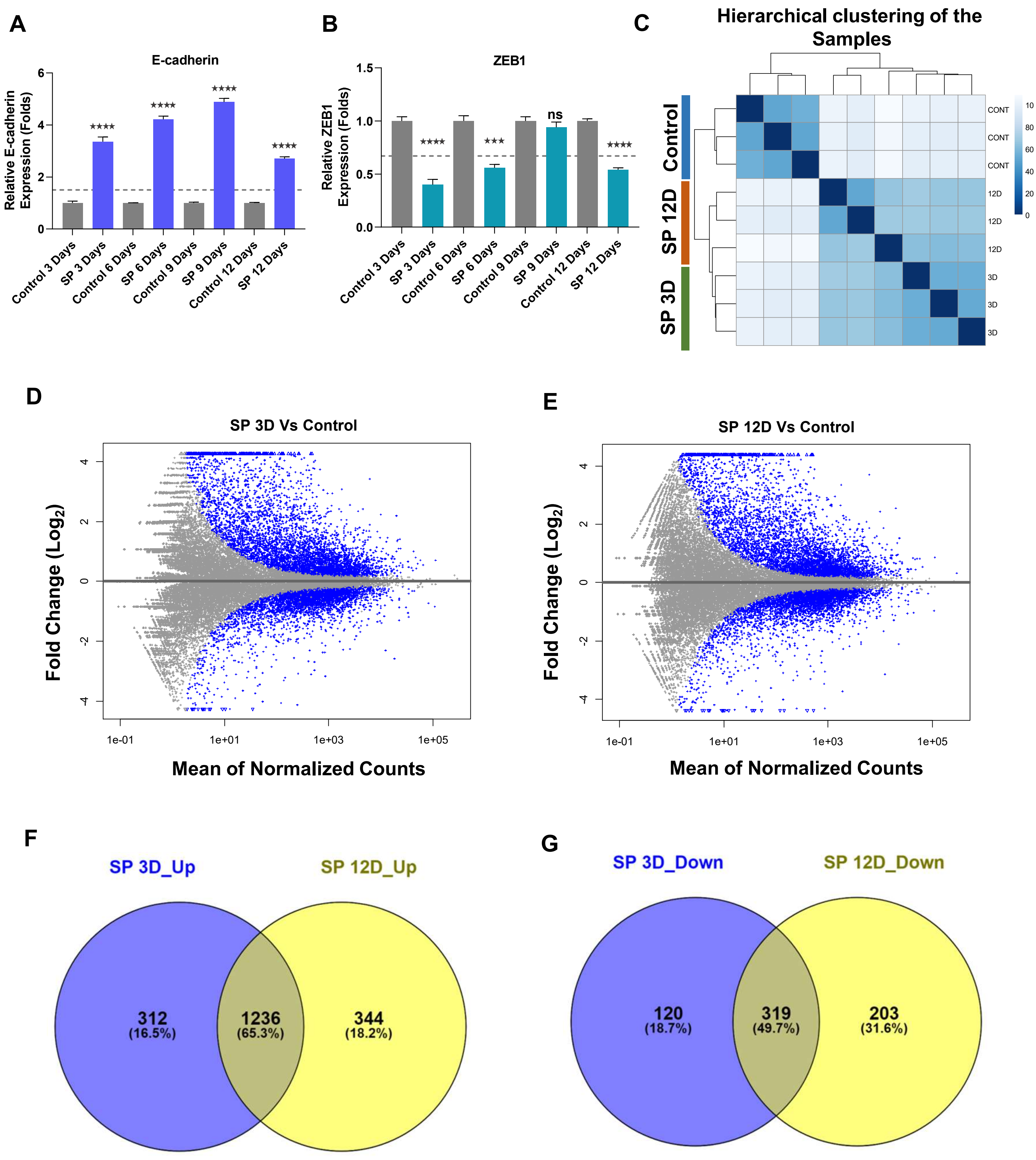

Supplementary Figure 10

A

NES=1.9; p=0.002; FDR=0.002

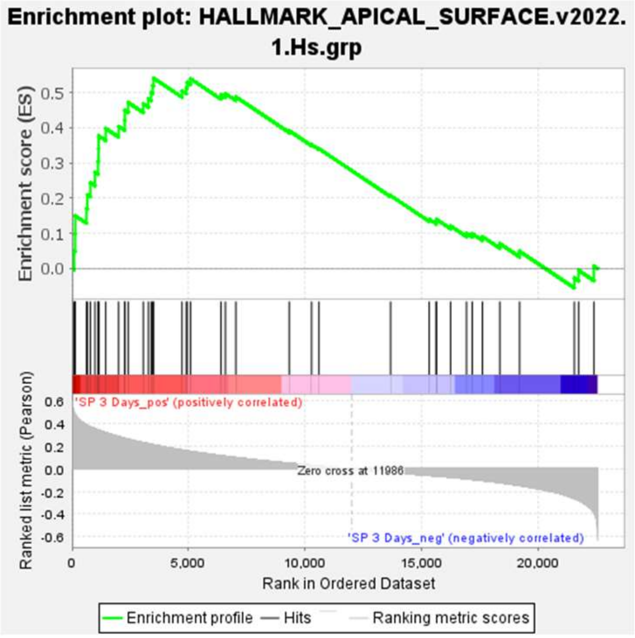


B

NES=2.3; p<0.01; FDR=0.0

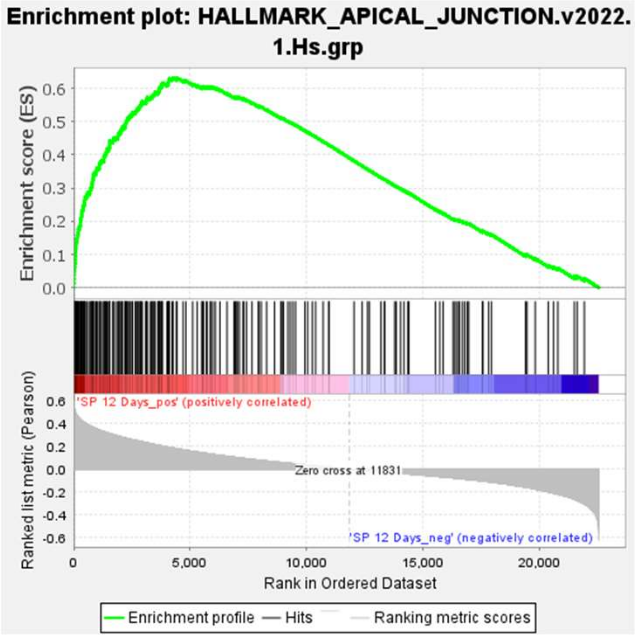


C

NES=1.7; p=0.006; FDR=0.009

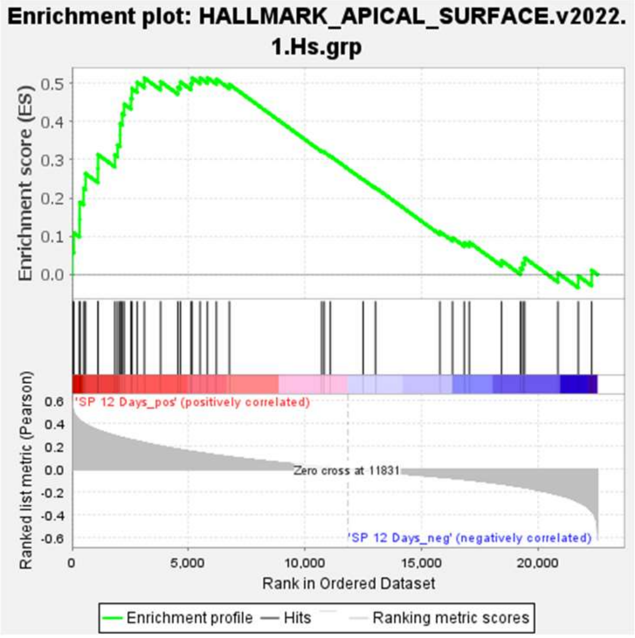

### Supplementary Figure 11

**A**

| Cluster | No. of Cells |
|---|---|
| 0 | 2065 |
| 1 | 1523 |
| 2 | 946 |
| 3 | 363 |
| 4 | 306 |

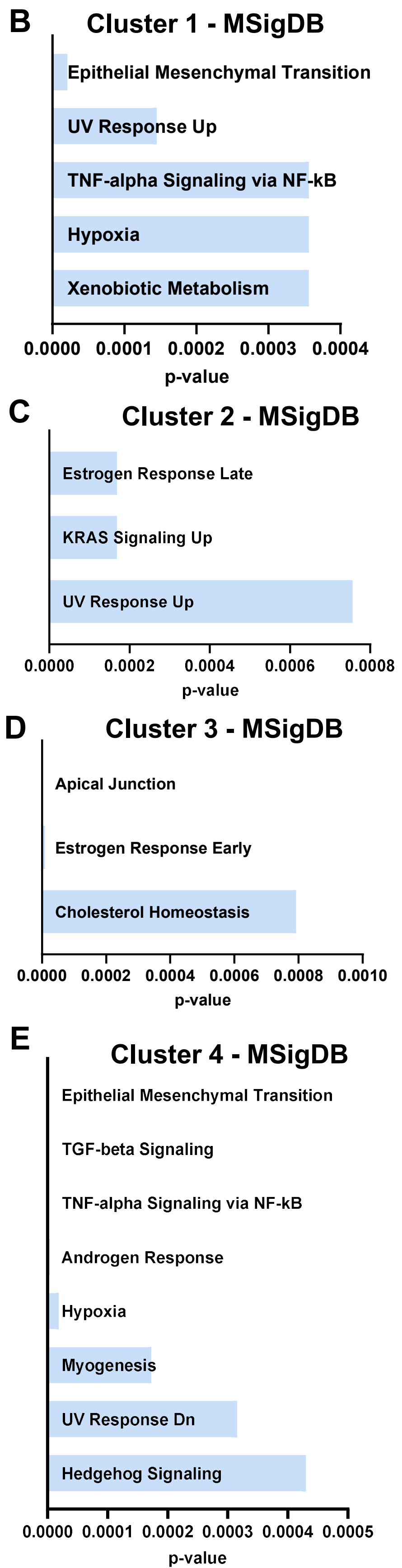


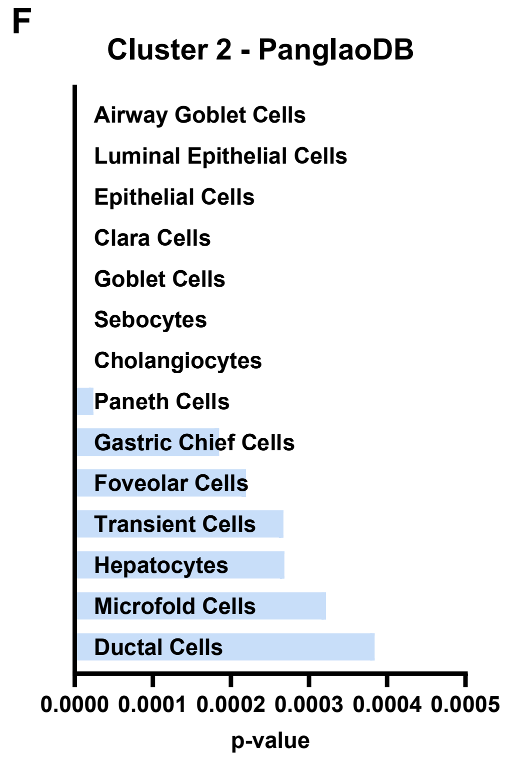


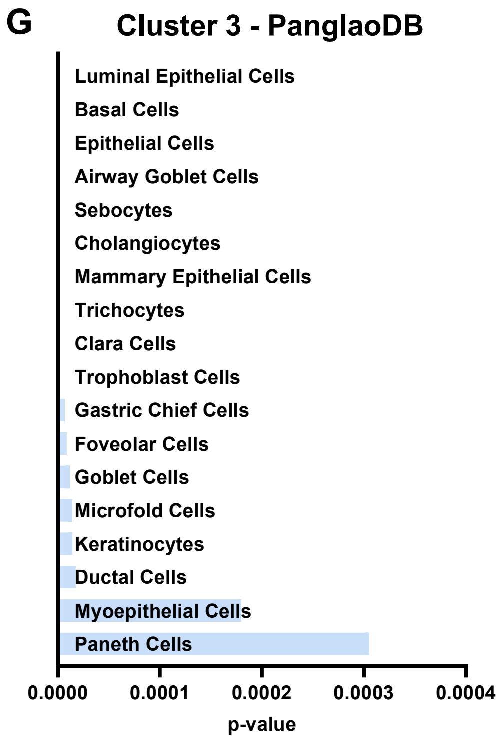


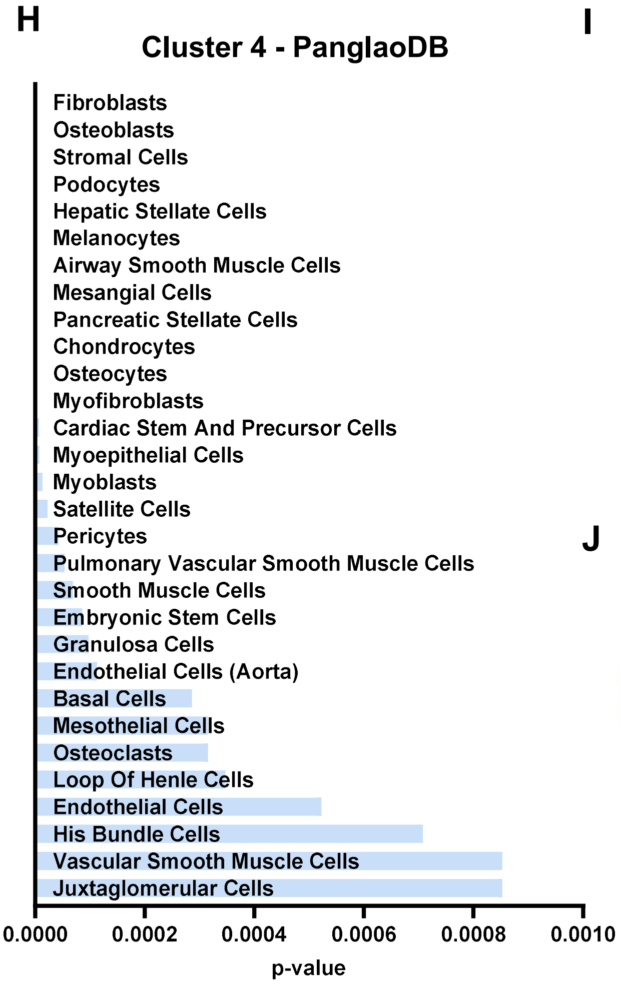


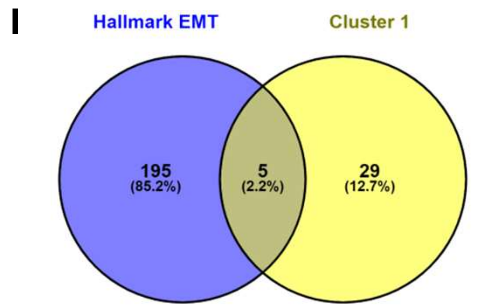


5 common genes in "Cluster 1" and "Hallmark EMT":

ID2
GPC1
SFRP1
PMP22
IGFBP2

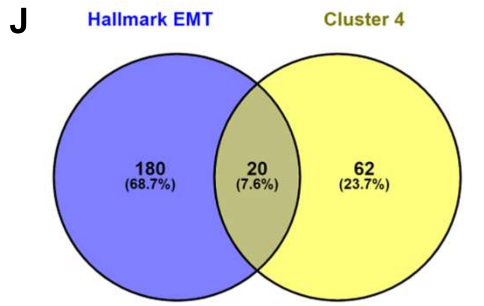


**20 common genes in "Cluster 4" and "Hallmark EMT":**

| | |
|---|---|
| PMEPA1 | COL4A1 |
| TGFBI | FGF2 |
| SERPINE1 | SPOCK1 |
| TAGLN | IL32 |
| GLIPR1 | CADM1 |
| COL4A2 | LAMC2 |
| TGM2 | THBS1 |
| COL5A1 | JUN |
| CDH2 | GREM1 |
| NNMT | ITGA2 |

**Supplementary Figure 12**

**A**

**GSE24202 – HMLE with TGFβ1**

**NES=1.4; p<0.01; FDR=0.12**

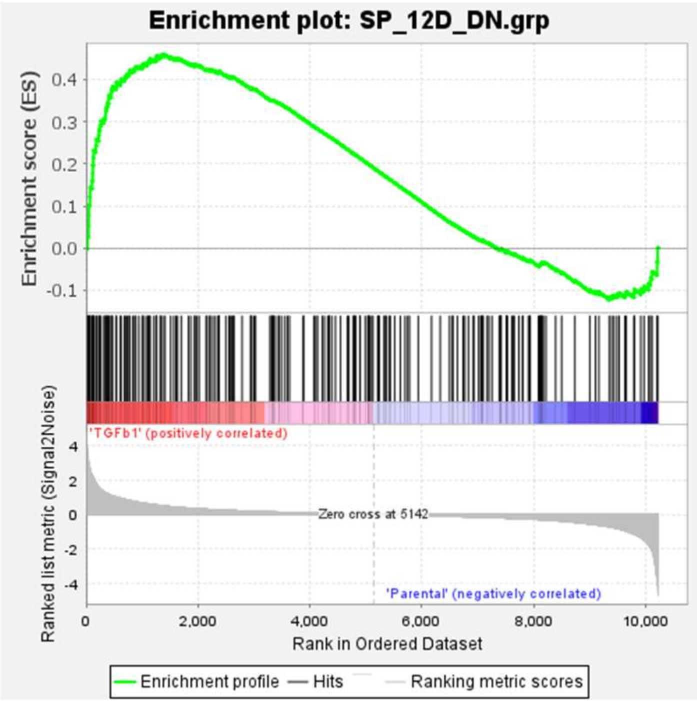


**B**

**GSE24202 – HMLE with TWIST**

**NES=1.7; p<0.01; FDR=0.06**

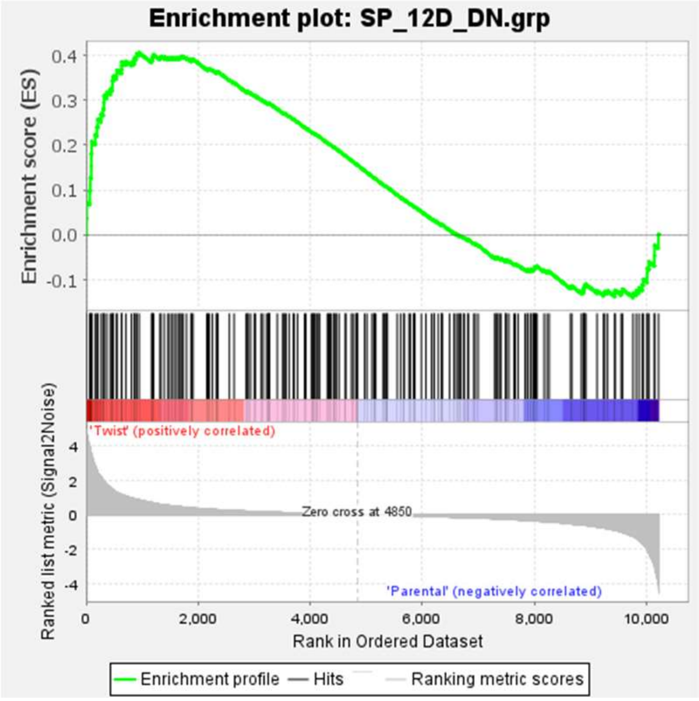


**C**

**GSE24202 – HMLE with GSC**

**NES=1.77; p<0.01; FDR=0.07**

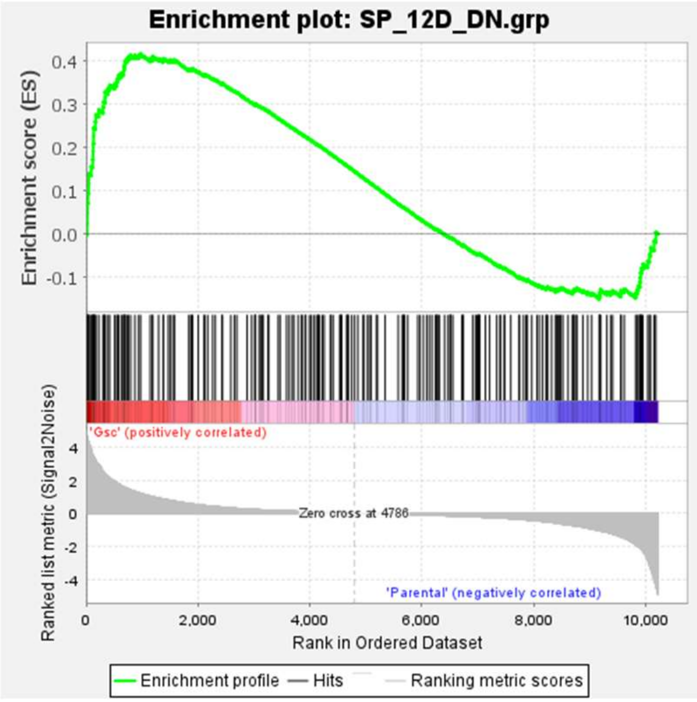


**D**

**GSE24202 – HMLE with SNAIL**

**NES=1.74; p<0.01; FDR=0.06**

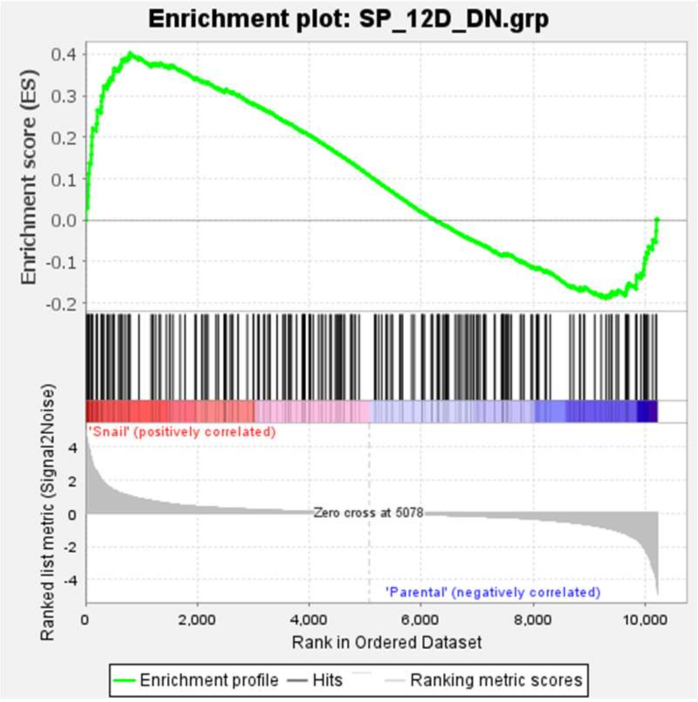

#### Supplementary Figure 13

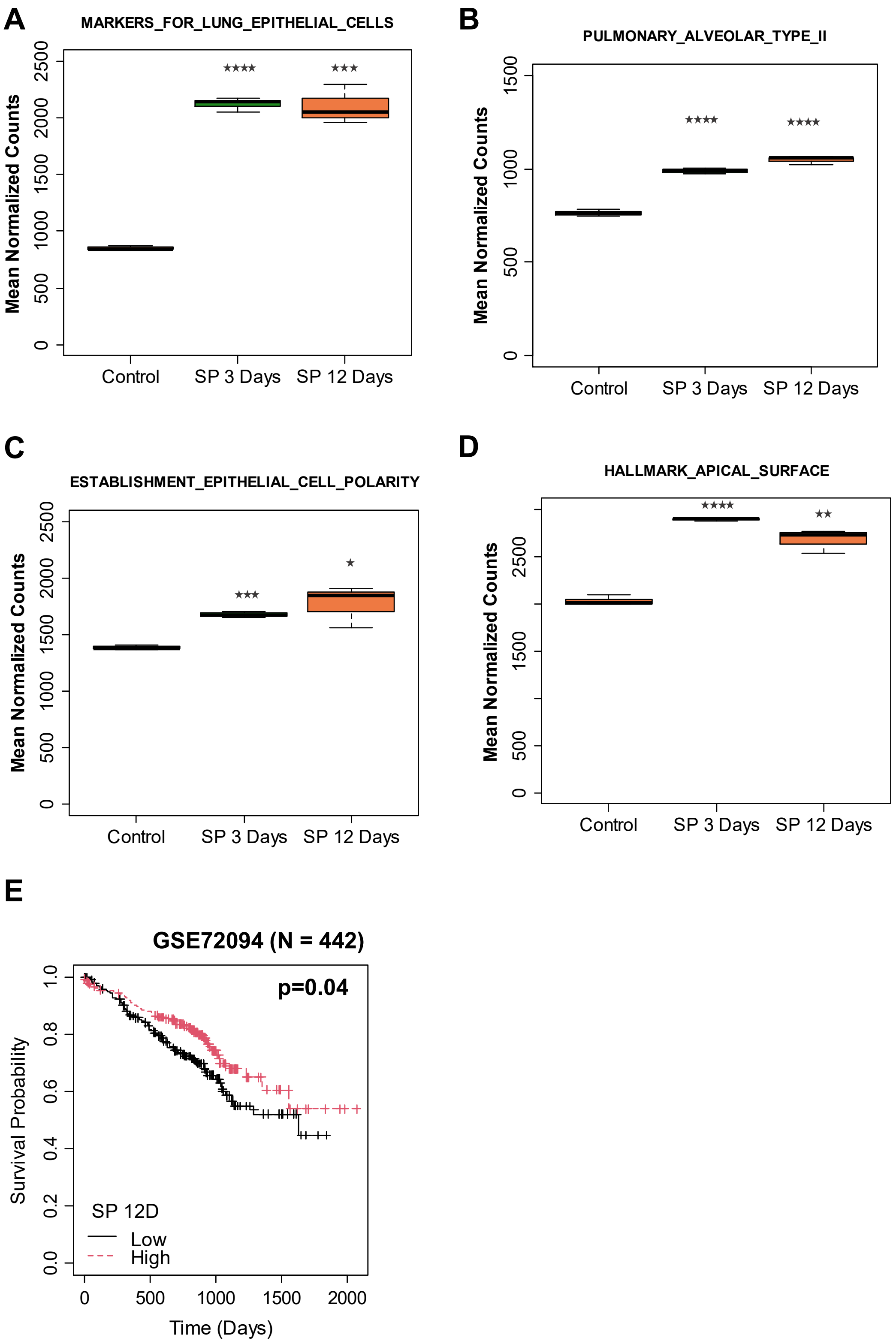

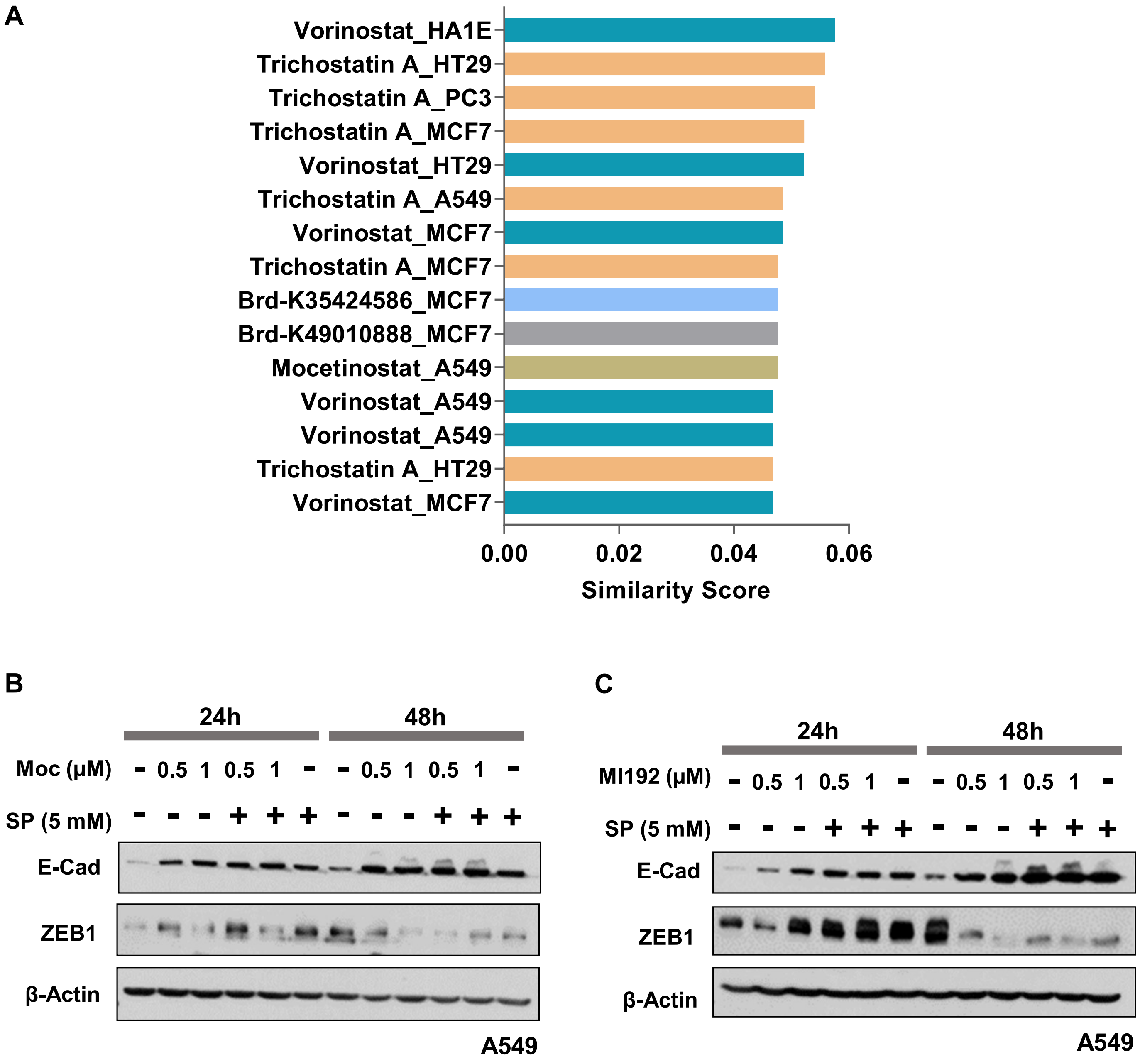
A
Vorinostat_HA1E
Trichostatin A_HT29
Trichostatin A_PC3
Trichostatin A_MCF7
Vorinostat_HT29
Trichostatin A_A549
Vorinostat_MCF7
Trichostatin A_MCF7
Brd-K35424586_MCF7
Brd-K49010888_MCF7
Mocetinostat_A549
Vorinostat_A549
Vorinostat_A549
Trichostatin A_HT29
Vorinostat_MCF7
0.00
0.02
0.04
0.06
Similarity Score
B
24h
48h
Moc (μM)
- 0.5 1 0.5 1 - - 0.5 1 0.5 1 -
SP (5 mM)
- - - + + + - - - + + +
E-Cad
ZEB1
β-Actin
A549
C
24h
48h
MI192 (μM)
- 0.5 1 0.5 1 - - 0.5 1 0.5 1 -
SP (5 mM)
- - - + + + - - - + + +
E-Cad
ZEB1
β-Actin
A549

**Supplementary Figure 15**

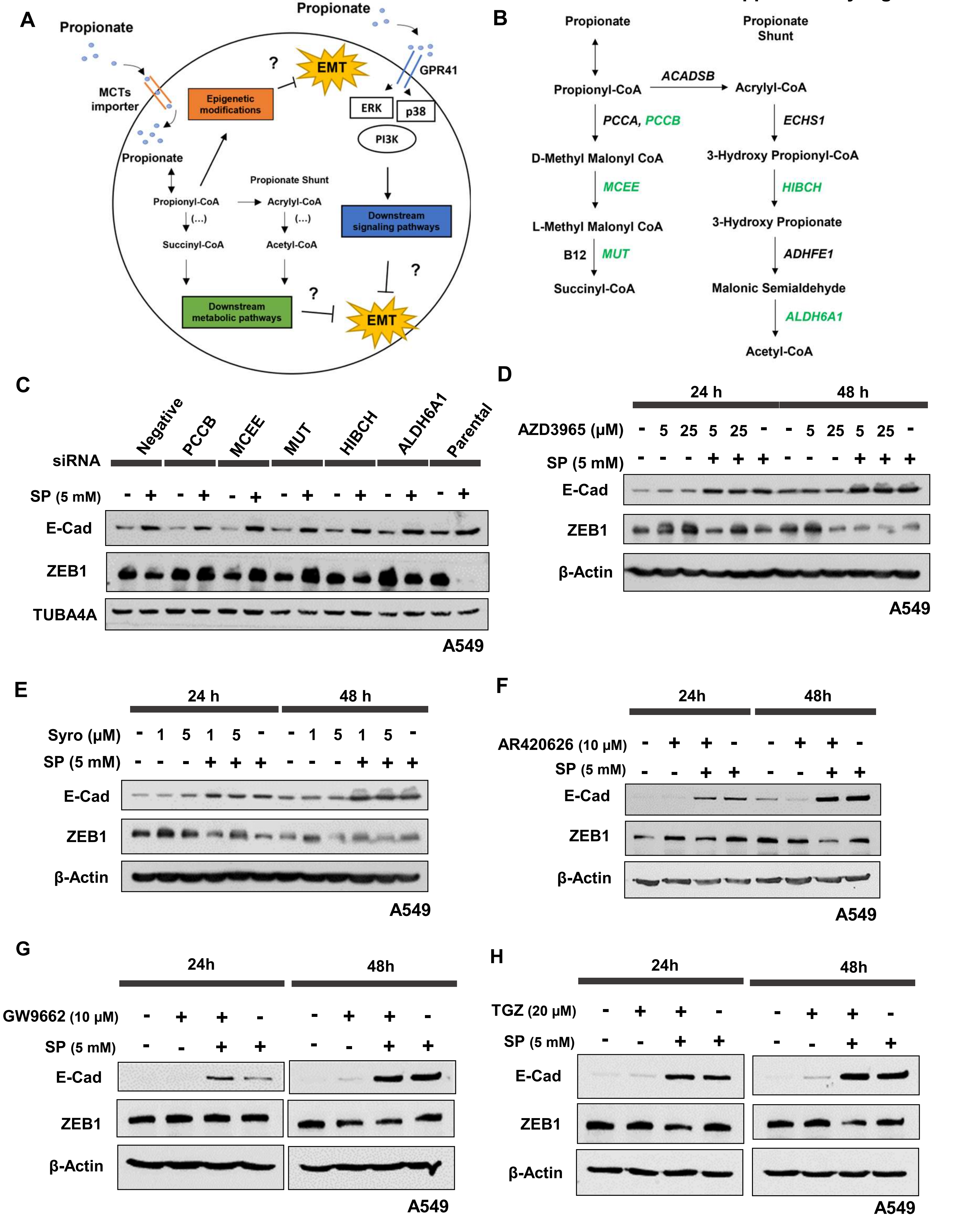

Supplementary Figure 16

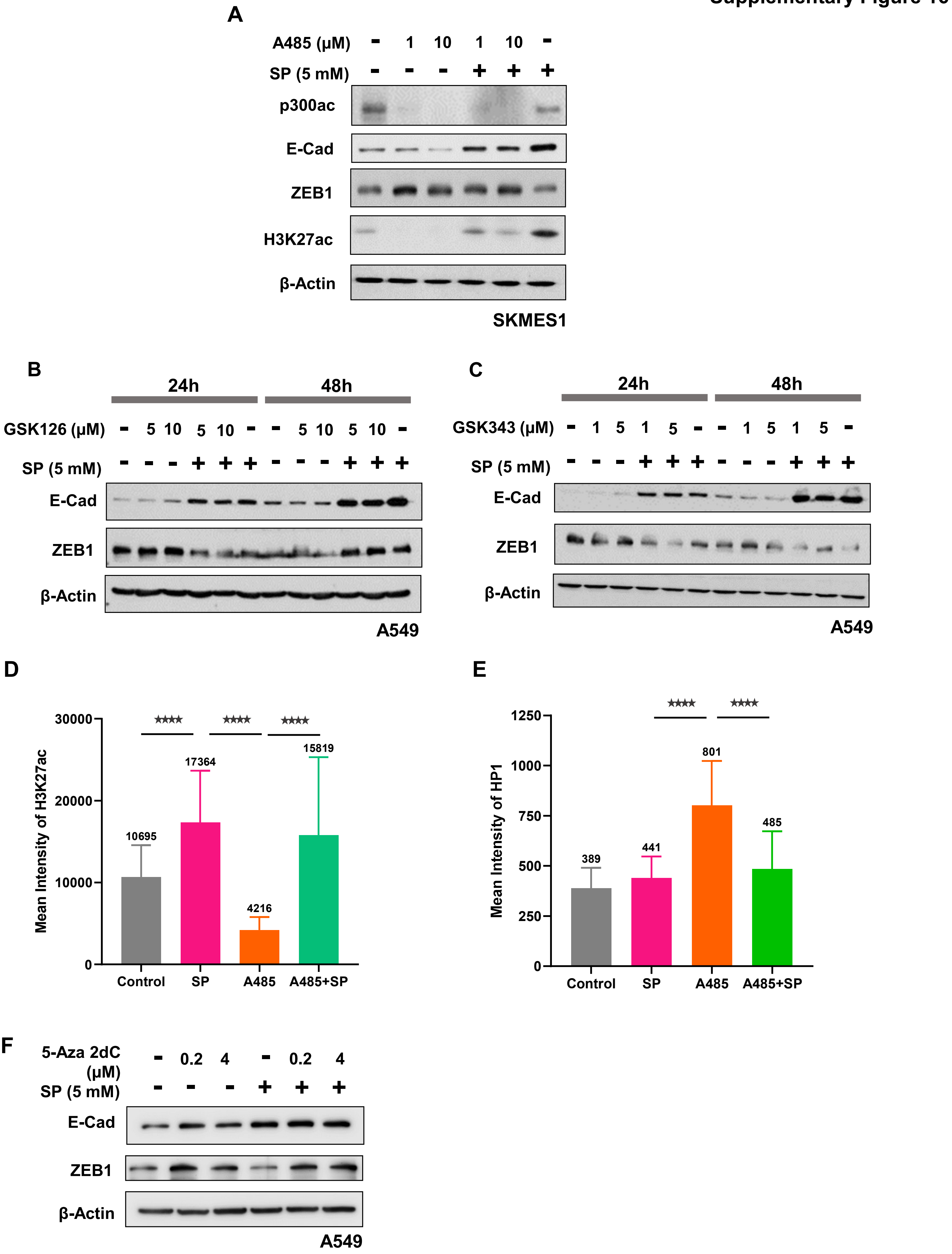

#### Supplementary Figure 17

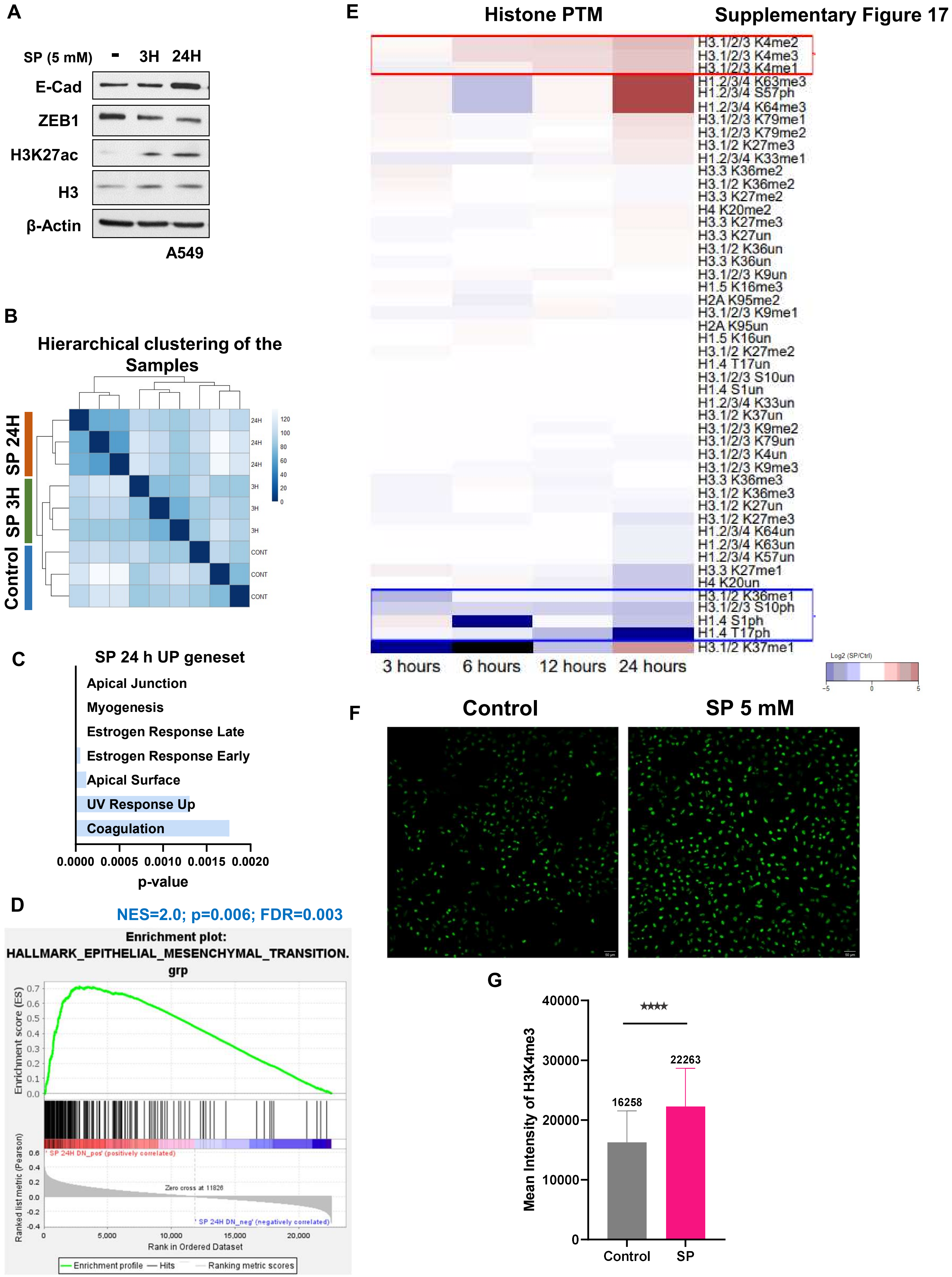

**Supplementary Figure 18**

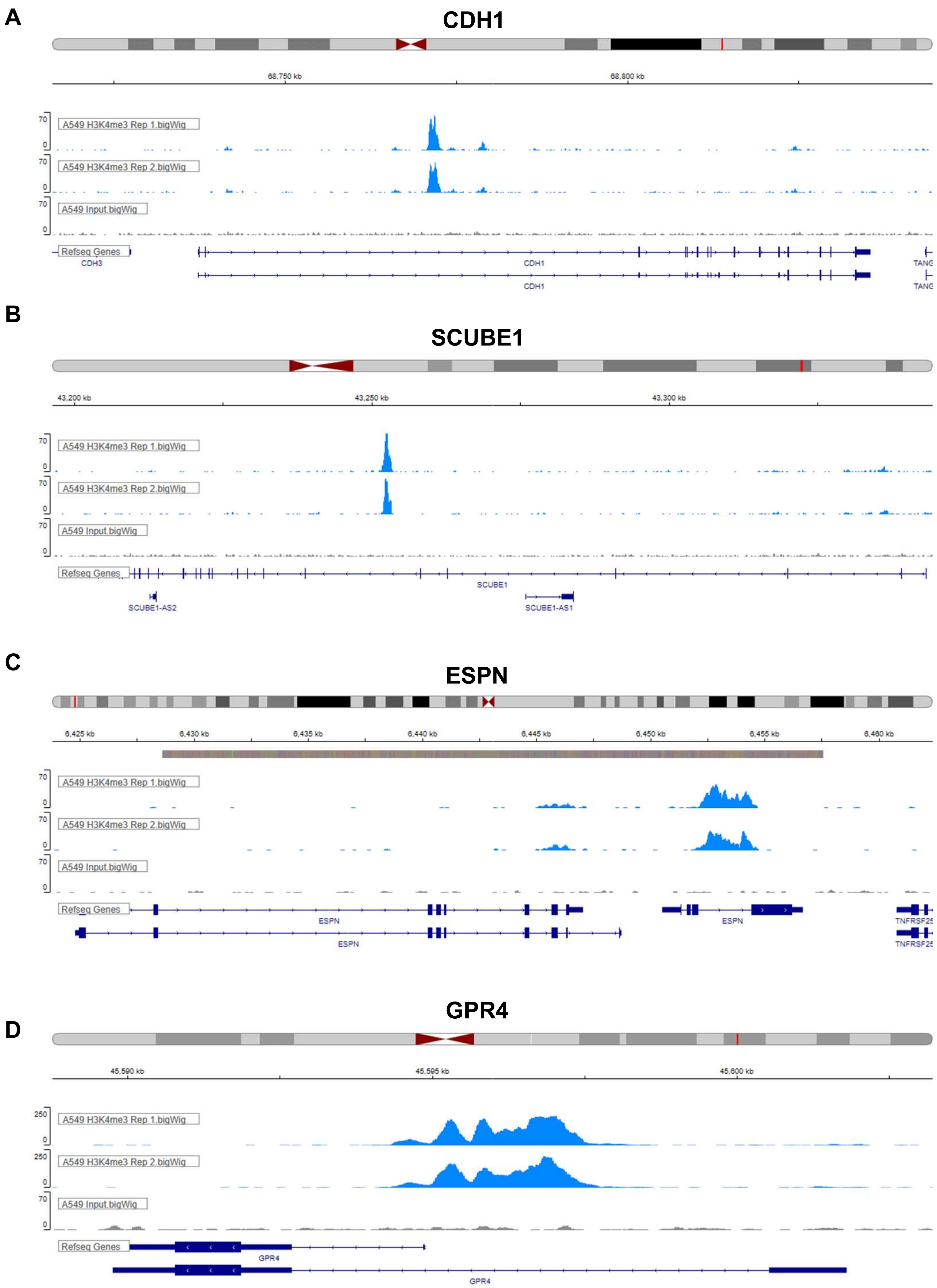

**A**

#### CDH1

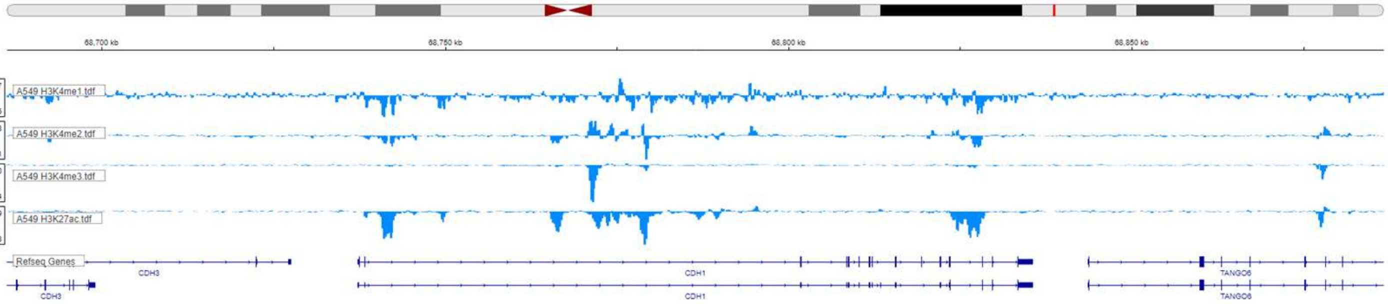


**B**

#### KRT19

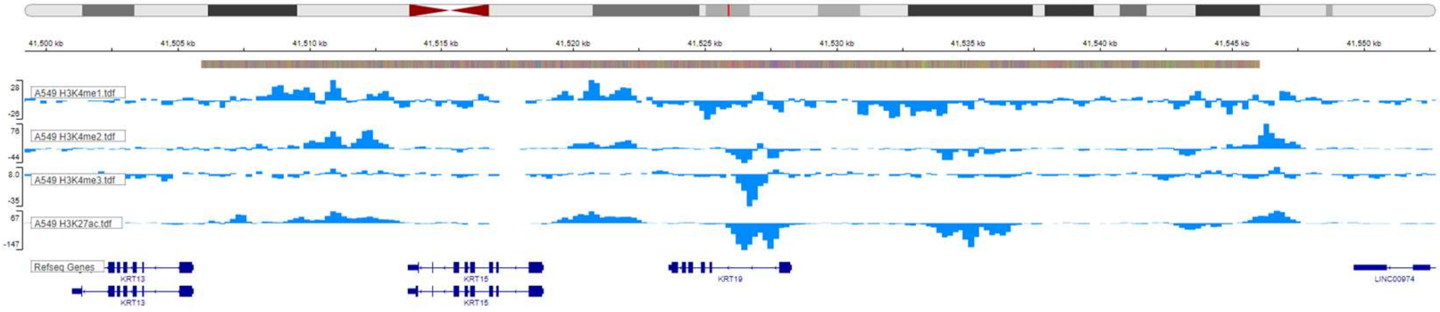


**C**

#### OVOL2

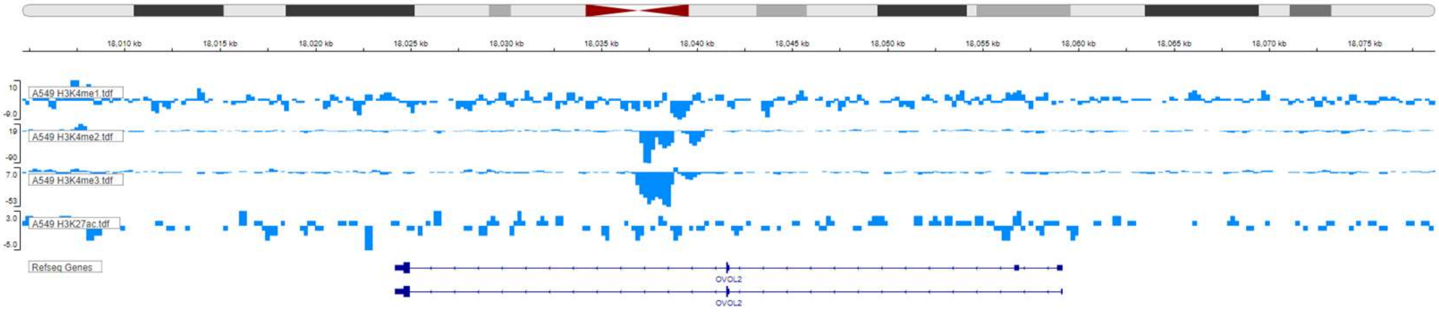


**D**

#### ZEB2

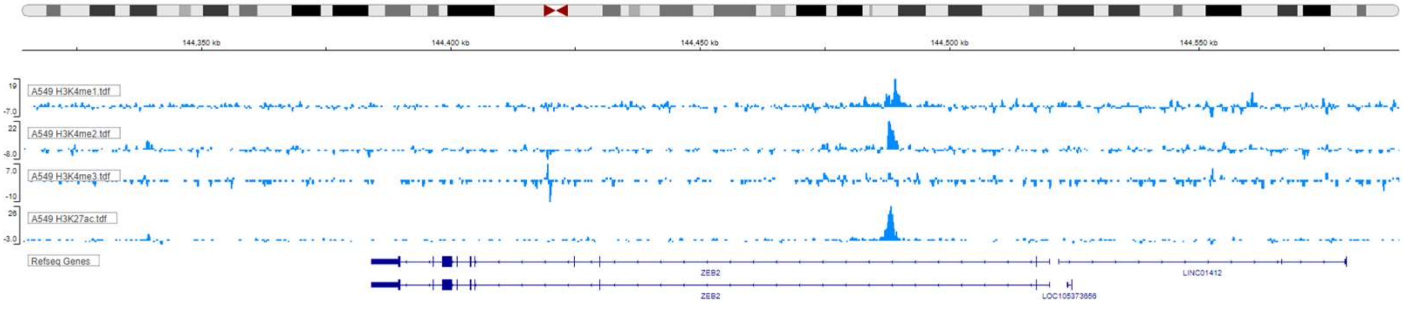

**Supplementary Table 1. List of Gene Expression Profiles with sample numbers used for the integrative functional genomic analysis.**

| Profiles | No. of samples |
|---|---|
| GSE8894 | 138 |
| GSE30219 | 293 |
| GSE31210 | 226 |
| GSE37745 | 196 |
| GSE50081 | 181 |
| GSE72094 | 442 |
| **Total** | **1476** |

**Supplementary Table 2. List of 77 genes in pan-cancer EMT gene signature for Mesenchymal and Epithelial states obtained from Mak *et al.,* 2016 used for the integrative functional genomic analysis.**

| Mesenchymal | | Epithelial |
|---|---|---|
| ADAM12 | PDGFRB | AP1G1 |
| ADAMTS12 | PMP22 | ATP8B1 |
| ADAMTS2 | POSTN | CDH1 |
| AEBP1 | SPARC | CDS1 |
| ANGPTL2 | SPOCK1 | CGN |
| ANTXR1 | SULF1 | CLDN4 |
| AXL | SYT11 | CNOT1 |
| BNC2 | THBS2 | CTNND1 |
| CALD1 | VCAN | DYNC1LI2 |
| CDH2 | VIM | ERBB3 |
| CMTM3 | ZEB2 | ESRP1 |
| CNRIP1 | | ESRP2 |
| COL10A1 | | F11R |
| COL1A1 | | GALNT3 |
| COL1A2 | | GPR56 |
| COL3A1 | | GRHL2 |
| COL5A1 | | HOOK1 |
| COL5A2 | | IRF6 |
| COL6A1 | | MAP7 |
| COL6A2 | | MARVELD2 |
| COL6A3 | | MARVELD3 |
| COL8A1 | | MYO5B |
| DACT1 | | OCLN |
| EMP3 | | PRSS8 |
| FAP | | SPINT1 |
| FBN1 | | |
| FN1 | | |
| FSTL1 | | |
| GPC6 | | |
| GYPC | | |
| HTRA1 | | |
| INHBA | | |
| ITGA11 | | |
| LOXL2 | | |
| LRRC15 | | |
| MMP2 | | |
| MSRB3 | | |
| NAP1L3 | | |
| NID2 | | |
| OLFML2B | | |
| PCOLCE | | |

**Supplementary Table 3. List of significant meta-correlation values with meta-pvalues of KEGG gene-sets associated with pan-cancer EMT gene signature in each gene expression dataset.**

| ID | Meta_Corr | Meta_pval |
|---|---|---|
| KEGG_ECM_RECEPTOR_INTERACTION | 0.82 | 1E-104 |
| KEGG_FOCAL_ADHESION | 0.75 | 2E-105 |
| KEGG_GLYCOSAMINOGLYCAN_BIOSYNTHESIS_CHONDROITIN_SULFATE | 0.71 | 2E-86 |
| KEGG_CYTOKINE_CYTOKINE_RECEPTOR_INTERACTION | 0.57 | 5E-96 |
| KEGG_CHEMOKINE_SIGNALING_PATHWAY | 0.54 | 1E-88 |
| KEGG_FC_GAMMA_R_MEDIATED_PHAGOCYTOSIS | 0.54 | 7E-116 |
| KEGG_TGF_BETA_SIGNALING_PATHWAY | 0.49 | 9E-18 |
| KEGG_GAP_JUNCTION | 0.49 | 2E-81 |
| KEGG_TOLL_LIKE_RECEPTOR_SIGNALING_PATHWAY | 0.48 | 2E-87 |
| KEGG_LEUKOCYTE_TRANSENDOTHELIAL_MIGRATION | 0.48 | 3E-90 |
| KEGG_HEMATOPOIETIC_CELL_LINEAGE | 0.47 | 2E-32 |
| KEGG_NOD_LIKE_RECEPTOR_SIGNALING_PATHWAY | 0.47 | 9E-83 |
| KEGG_NATURAL_KILLER_CELL_MEDIATED_CYTOTOXICITY | 0.44 | 2E-46 |
| KEGG_B_CELL_RECEPTOR_SIGNALING_PATHWAY | 0.44 | 1E-72 |
| KEGG_JAK_STAT_SIGNALING_PATHWAY | 0.44 | 4E-35 |
| KEGG_CELL_ADHESION_MOLECULES_CAMS | 0.43 | 1E-43 |
| KEGG_NEUROACTIVE_LIGAND_RECEPTOR_INTERACTION | 0.40 | 9E-33 |
| KEGG_AXON_GUIDANCE | 0.39 | 1E-41 |
| KEGG_CALCIUM_SIGNALING_PATHWAY | 0.39 | 2E-11 |
| KEGG_T_CELL_RECEPTOR_SIGNALING_PATHWAY | 0.37 | 7E-51 |
| KEGG_CYTOSOLIC_DNA_SENSING_PATHWAY | 0.33 | 1E-20 |
| KEGG_GLYCOSAMINOGLYCAN_BIOSYNTHESIS_HEPARAN_SULFATE | 0.32 | 2E-37 |
| KEGG_GLYCOSPHINGOLIPID_BIOSYNTHESIS_GLOBO_SERIES | 0.32 | 8E-37 |
| KEGG_MAPK_SIGNALING_PATHWAY | 0.30 | 7E-27 |
| KEGG_CITRATE_CYCLE_TCA_CYCLE | -0.31 | 5E-19 |
| KEGG_STEROID_BIOSYNTHESIS | -0.33 | 2E-17 |
| KEGG_HISTIDINE_METABOLISM | -0.35 | 4E-17 |
| KEGG_GLYCEROPHOSPHOLIPID_METABOLISM | -0.37 | 2E-17 |
| KEGG_LIMONENE_AND_PINENE_DEGRADATION | -0.37 | 1E-07 |
| KEGG_PYRUVATE_METABOLISM | -0.39 | 4E-18 |
| KEGG_SELENOAMINO_ACID_METABOLISM | -0.43 | 1E-12 |
| KEGG_TERPENOID_BACKBONE_BIOSYNTHESIS | -0.44 | 1E-57 |
| KEGG_BUTANOATE_METABOLISM | -0.46 | 3E-22 |
| KEGG_FATTY_ACID_METABOLISM | -0.46 | 2E-21 |
| KEGG_PROPANOATE_METABOLISM | -0.46 | 1E-16 |
| KEGG_VALINE_LEUCINE_AND_ISOLEUCINE_DEGRADATION | -0.47 | 2E-14 |
| KEGG_GLYCOSYLPHOSPHATIDYLINOSITOL_GPI_ANCHOR_BIOSYNTHESIS | -0.47 | 3E-29 |
| KEGG_PEROXISOME | -0.52 | 2E-14 |

**Supplementary Table 4. List of significant meta-correlation values with meta-pvalues of REACTOME gene-sets associated with pan-cancer EMT gene signature in each gene expression dataset.**

| ID | Meta_Corr | Meta_pval |
|---|---|---|
| REACTOME_CHONDROITIN_SULFATE_BIOSYNTHESIS | 0.83 | 1E-94 |
| REACTOME_PLATELET_ADHESION_TO_EXPOSED_COLLAGEN | 0.76 | 5E-91 |
| REACTOME_CHONDROITIN_SULFATE_DERMATAN_SULFATE_METABOLISM | 0.70 | 6E-94 |
| REACTOME_GLYCOSAMINOGLYCAN_METABOLISM | 0.70 | 8E-96 |
| REACTOME_INTEGRIN_CELL_SURFACE_INTERACTIONS | 0.67 | 1E-79 |
| REACTOME_CS_DS_DEGRADATION | 0.64 | 5E-38 |
| REACTOME_KERATAN_SULFATE_BIOSYNTHESIS | 0.64 | 3E-56 |
| REACTOME_KERATAN_SULFATE_DEGRADATION | 0.64 | 8E-166 |
| REACTOME_KERATAN_SULFATE_KERATIN_METABOLISM | 0.63 | 1E-39 |
| REACTOME_MYOGENESIS | 0.53 | 5E-22 |
| REACTOME_HEPARAN_SULFATE_HEPARIN_HS_GAG_METABOLISM | 0.51 | 4E-79 |
| REACTOME_EICOSANOID_LIGAND_BINDING_RECEPTORS | 0.49 | 4E-29 |
| REACTOME_NITRIC_OXIDE_STIMULATES_GUANYLATE_CYCLASE | 0.46 | 2E-09 |
| REACTOME_GAP_JUNCTION_DEGRADATION | 0.44 | 2E-18 |
| REACTOME_METABOLISM_OF_CARBOHYDRATES | 0.40 | 4E-18 |
| REACTOME_INHIBITION_OF_INSULIN_SECRETION_BY_ADRENALINE_NORADRENALINE | 0.38 | 3E-08 |
| REACTOME_HDL_MEDIATED_LIPID_TRANSPORT | 0.37 | 1E-44 |
| REACTOME_NUCLEOTIDE_BINDING_DOMAIN_LEUCINE_RICH_REPEAT_CONTAINING_RECEPTOR_NLR_SIGNALING_PATHWAYS | 0.37 | 7E-49 |
| REACTOME_NUCLEOTIDE_LIKE_PURINERGIC_RECEPTORS | 0.35 | 1E-43 |
| REACTOME_TRANSPORT_TO_THE_GOLGI_AND_SUBSEQUENT_MODIFICATION | 0.35 | 2E-26 |
| REACTOME_GROWTH_HORMONE_RECEPTOR_SIGNALING | 0.34 | 1E-26 |
| REACTOME_HS_GAG_BIOSYNTHESIS | 0.33 | 4E-40 |
| REACTOME_GLUCAGON_SIGNALING_IN_METABOLIC_REGULATION | 0.33 | 1E-04 |
| REACTOME_BASE_FREE_SUGAR_PHOSPHATE_REMOVAL_VIA_THE_SINGLE_NUCLEOTIDE_REPLACEMENT_PATHWAY | -0.30 | 8E-09 |
| REACTOME_SULFUR_AMINO_ACID_METABOLISM | -0.31 | 3E-13 |
| REACTOME_RESPIRATORY_ELECTRON_TRANSPORT_ATP_SYNTHESIS_BY_CHEMIOSMOTIC_COUPLING_AND_HEAT_PRODUCTION_BY_UNCOUPLING_PROTEINS | -0.32 | 2E-17 |
| REACTOME_TCA_CYCLE_AND_RESPIRATORY_ELECTRON_TRANSPORT | -0.32 | 6E-23 |
| REACTOME_TRIGLYCERIDE_BIOSYNTHESIS | -0.32 | 8E-29 |
| REACTOME_RESPIRATORY_ELECTRON_TRANSPORT | -0.33 | 4E-22 |
| REACTOME_CITRIC_ACID_CYCLE_TCA_CYCLE | -0.33 | 3E-23 |
| REACTOME_PEROXISOMAL_LIPID_METABOLISM | -0.36 | 3E-28 |
| REACTOME_FATTY_ACID_TRIACYLGLYCEROL_AND_KETONE_BODY_METABOLISM | -0.36 | 2E-15 |
| REACTOME_MITOCHONDRIAL_TRNA_AMINOACYLATION | -0.37 | 8E-48 |
| REACTOME_SYNTHESIS_OF_PC | -0.38 | 5E-22 |
| REACTOME_GLYCEROPHOSPHOLIPID_BIOSYNTHESIS | -0.38 | 2E-13 |
| REACTOME_VITAMIN_B5_PANTOTHENATE_METABOLISM | -0.38 | 2E-26 |
| REACTOME_SYNTHESIS_OF_PE | -0.38 | 2E-09 |
| REACTOME_CHOLESTEROL_BIOSYNTHESIS | -0.40 | 2E-39 |
| REACTOME_SYNTHESIS_OF_GLYCOSYLPHOSPHATIDYLINOSITOL_GPI | -0.46 | 2E-20 |
| REACTOME_ACTIVATED_AMPK_STIMULATES_FATTY_ACID_OXIDATION_IN_MUSCLE | -0.47 | 1E-27 |
| REACTOME_BRANCHED_CHAIN_AMINO_ACID_CATABOLISM | -0.48 | 8E-14 |
| REACTOME_MITOCHONDRIAL_FATTY_ACID_BETA_OXIDATION | -0.50 | 6E-18 |

**Supplementary Table 5. List of marker genes in each cell type cluster identified from single cell RNA sequencing of A549 cells**

| Cluster_0 | Cluster_1 | Cluster_2 | | Cluster_3 | | Cluster_4 | | |
|---|---|---|---|---|---|---|---|---|
| FXYD2 | HMOX1 | AGR2 | SPDEF | CEACAM6 | GACAT2 | PMEPA1 | INPP4B | PGRMC2 |
| CD24 | AKAP12 | SLC12A2 | TST | KRT19 | CALB2 | CCDC80 | LBH | ARHGEF18 |
| AXL | ID2 | FN1 | PTGS2 | AGR2 | NET1 | TGFBI | EPHB2 | ACSL4 |
| KRT19 | RSPO3 | MUC5AC | IFITM2 | PRSS3 | CDC42EP3 | IGFBP7 | MARCKSL1 | SEMA3C |
| MMP7 | CTSB | AKR1C1 | FOS | MUC5AC | GALNT5 | SERPINE1 | JAG1 | MAP7 |
| EEF1A2 | TESC | CPLX2 | PLS1 | ITGB4 | LMO7 | TAGLN | CDK6 | MFGE8 |
| S100A3 | KYNU | CEACAM6 | COL5A2 | FGFBP1 | SLCO1B3 | GLIPR1 | ANKLE2 | NUAK1 |
| | GDF15 | PDLIM5 | CLDN2 | CAVIN3 | ITGA2 | SOX4 | HMGA2 | EEA1 |
| | EFHD2 | MUC5B | TSPAN13 | CRIP2 | JUP | COL4A2 | FGF2 | GRB10 |
| | MFF | LCN2 | KLF13 | CAPG | PCED1B | JUNB | IGF1R | DCBLD1 |
| | NAMPT | ANXA13 | STEAP1 | LCN2 | EPS8 | PCED1B | SKIL | MOB3B |
| | CYP24A1 | SYT1 | LRP10 | SOX4 | ADD3 | PDLIM7 | IL11 | FST |
| | GPC1 | RAP1GAP | JUP | CNTN1 | GAL3ST1 | PGM2L1 | SPOCK1 | SCN9A |
| | PAQR5 | EPHX1 | PLA2G4A | VAMP8 | KRT15 | PLEK2 | PHLDB2 | DBN1 |
| | TNNT1 | MMP7 | HLA-DMB | SYT1 | MDK | CITED4 | C15orf48 | |
| | EPAS1 | CYP1B1 | SMIM14 | PLEK2 | | TGM2 | PALLD | |
| | CCND3 | MTUS1 | BCAS1 | RHOD | | COL5A1 | IL32 | |
| | SRGN | CCPG1 | RNASE4 | ZNF185 | | ETS2 | ANTXR1 | |
| | CEBPB | SLC9A3R2 | | ITPRID2 | | SLC26A2 | CDC42EP3 | |
| | KCNMA1 | EPS8 | | ITGA6 | | CDH2 | RAI14 | |
| | HSPA2 | CFH | | MUC5B | | GLS | CADM1 | |
| | AGFG1 | SLPI | | SERPINB1 | | NCOR2 | LAMC2 | |
| | ATIC | TM4SF20 | | ETHE1 | | NRP2 | GALNT10 | |
| | SFRP1 | SCP2 | | TMBIM4 | | NNMT | THBS1 | |
| | RHOQ | ARHGAP18 | | UPK1B | | RAB3B | CEP170 | |
| | DARS | CAMK2N1 | | ANXA13 | | FOXP1 | JUN | |
| | NPC1 | BAMBI | | MARCKSL1 | | KCNMA1 | EPB41L2 | |
| | PMP22 | AQP3 | | ALDH2 | | FRMD6 | GREM1 | |
| | MAP3K20 | CNTN1 | | IGFL2-AS1 | | COL4A1 | ITGA2 | |
| | ADM | INSL4 | | MMP7 | | MT1X | TANC2 | |
| | HILPDA | RDH10 | | PLAUR | | TGFB1I1 | LYPD1 | |
| | STAT1 | TPD52L1 | | LAMB3 | | DUSP1 | TNFAIP8 | |
| | IGFBP2 | S100P | | NT5E | | PODXL | PACS1 | |
| | SLC16A3 | FGL1 | | AP1S3 | | SPDL1 | CCN2 | |

**Supplementary Table 6. List of mimics identified from the similarity metrics of L1000CDS[2] for the differentially expressed genes compared between SP 3 days and control from**

| Rank | Score | Perturbation | Cell Lines | Dose | Time |
|---|---|---|---|---|---|
| 1 | 0.0576 | vorinostat | HA1E | 11.1um | 6.0h |
| 2 | 0.0558 | trichostatin A | HT29 | 10.0um | 6.0h |
| 3 | 0.054 | trichostatin A | PC3 | 10.0um | 6.0h |
| 4 | 0.0522 | trichostatin A | MCF7 | 10.0um | 24.0h |
| 5 | 0.0522 | vorinostat | HT29 | 10.0um | 24.0h |
| 6 | 0.0486 | trichostatin A | A549 | 10.0um | 6.0h |
| 7 | 0.0486 | vorinostat | MCF7 | 10.0um | 24.0h |
| 8 | 0.0477 | trichostatin A | MCF7 | 10.0um | 24.0h |
| 9 | 0.0477 | BRD-K35424586 | MCF7 | 10.0um | 24.0h |
| 10 | 0.0477 | BRD-K49010888 | MCF7 | 10.0um | 24.0h |
| 11 | 0.0477 | mocetinostat | A549 | 10um | 24h |
| 12 | 0.0468 | vorinostat | A549 | 10.0um | 24.0h |
| 13 | 0.0468 | vorinostat | A549 | 10.0um | 6.0h |
| 14 | 0.0468 | trichostatin A | HT29 | 10.0um | 6.0h |
| 15 | 0.0468 | vorinostat | MCF7 | 10.0um | 24.0h |
| 16 | 0.0468 | mocetinostat | A549 | 3.33um | 24h |
| 17 | 0.0459 | vorinostat | A375 | 10.0um | 6.0h |
| 18 | 0.0459 | vorinostat | HT29 | 10.0um | 6.0h |
| 19 | 0.045 | trichostatin A | A549 | 10.0um | 6.0h |
| 20 | 0.045 | vorinostat | HCC515 | 10.0um | 24.0h |
| 21 | 0.045 | vorinostat | PC3 | 10.0um | 6.0h |
| 22 | 0.045 | trichostatin A | A375 | 10.0um | 6.0h |
| 23 | 0.0441 | vorinostat | HCC515 | 11.1um | 24.0h |
| 24 | 0.0441 | 480743.cdx | HT29 | 80.0um | 24.0h |
| 25 | 0.0441 | HDAC6 inhibitor ISOX | HT29 | 10.0um | 6.0h |
| 26 | 0.0441 | trichostatin A | A549 | 10.0um | 6.0h |
| 27 | 0.0441 | BRD-K77908580 | MCF7 | 10.0um | 24.0h |
| 28 | 0.0441 | trichostatin A | A375 | 10.0um | 6.0h |
| 29 | 0.0432 | vorinostat | HCC515 | 10.0um | 24.0h |
| 30 | 0.0432 | trichostatin A | PC3 | 10.0um | 6.0h |
| 31 | 0.0432 | trichostatin A | HCC515 | 10.0um | 6.0h |
| 32 | 0.0432 | trichostatin A | HCC515 | 10.0um | 24.0h |
| 33 | 0.0432 | HDAC6 inhibitor ISOX | MCF7 | 10.0um | 24.0h |
| 34 | 0.0432 | vorinostat | VCAP | 11.1um | 6.0h |
| 35 | 0.0432 | BRD-K12867552 | MCF7 | 10.0um | 24.0h |
| 36 | 0.0432 | BRD-K52522949 | PC3 | 10.0um | 6.0h |
| 37 | 0.0423 | vorinostat | HA1E | 10.0um | 6.0h |
| 38 | 0.0423 | trichostatin A | A375 | 10.0um | 6.0h |
| 39 | 0.0423 | HDAC6 inhibitor ISOX | VCAP | 10.0um | 6.0h |
| 40 | 0.0423 | trichostatin A | A549 | 10.0um | 6.0h |
| 41 | 0.0423 | trichostatin A | A549 | 10.0um | 6.0h |
| 42 | 0.0423 | trichostatin A | MCF7 | 10.0um | 6.0h |
| 43 | 0.0423 | trichostatin A | HT29 | 10.0um | 6.0h |
| 44 | 0.0423 | BRD-K55116708 | MCF7 | 10.0um | 24.0h |
| 45 | 0.0423 | BRD-A94377914 | MCF7 | 10.0um | 6.0h |
| 46 | 0.0414 | HDAC6 inhibitor ISOX | A375 | 10.0um | 24.0h |
| 47 | 0.0414 | HDAC6 inhibitor ISOX | A549 | 10.0um | 6.0h |
| 48 | 0.0414 | vorinostat | A549 | 11.1um | 6.0h |
| 49 | 0.0414 | vorinostat | HCC515 | 11.1um | 6.0h |
| 50 | 0.0414 | vorinostat | HT29 | 11.1um | 24.0h |

**Supplementary Table 7. List of mimics identified from the similarity metrics of L1000CDS$^2$ for the differentially expressed genes compared between SP 12 days and control from RNA-Seq analysis.**

| Rank | Score | Perturbation | Cell Lines | Dose | Time |
|---|---|---|---|---|---|
| 1 | 0.0595 | vorinostat | HA1E | 11.1um | 6.0h |
| 2 | 0.0578 | vorinostat | A549 | 10.0um | 24.0h |
| 3 | 0.0561 | vorinostat | HCC515 | 10.0um | 24.0h |
| 4 | 0.0553 | trichostatin A | PC3 | 10.0um | 6.0h |
| 5 | 0.0544 | vorinostat | MCF7 | 10.0um | 24.0h |
| 6 | 0.0536 | trichostatin A | HT29 | 10.0um | 6.0h |
| 7 | 0.0536 | phorbol-12-myristate-13-acetate (PMA) | MCF7 | 10.0um | 24.0h |
| 8 | 0.0528 | vorinostat | HCC515 | 10.0um | 24.0h |
| 9 | 0.0528 | vorinostat | HCC515 | 11.1um | 24.0h |
| 10 | 0.0528 | vorinostat | MCF7 | 10.0um | 24.0h |
| 11 | 0.0519 | vorinostat | HCC515 | 10.0um | 24.0h |
| 12 | 0.0519 | trichostatin A | MCF7 | 10.0um | 24.0h |
| 13 | 0.0519 | vorinostat | A549 | 10.0um | 6.0h |
| 14 | 0.0519 | vorinostat | HT29 | 10.0um | 24.0h |
| 15 | 0.0519 | trichostatin A | MCF7 | 10.0um | 24.0h |
| 16 | 0.0511 | trichostatin A | A549 | 10.0um | 6.0h |
| 17 | 0.0511 | vorinostat | PC3 | 10.0um | 6.0h |
| 18 | 0.0503 | HDAC6 inhibitor ISOX | A375 | 10.0um | 24.0h |
| 19 | 0.0503 | BRD-K49010888 | MCF7 | 10.0um | 24.0h |
| 20 | 0.0494 | trichostatin A | HCC515 | 10.0um | 24.0h |
| 21 | 0.0494 | BRD-K12867552 | MCF7 | 10.0um | 24.0h |
| 22 | 0.0494 | mocetinostat | A549 | 10um | 24h |
| 23 | 0.0477 | trichostatin A | A549 | 10.0um | 6.0h |
| 24 | 0.0477 | BRD-K77908580 | MCF7 | 10.0um | 24.0h |
| 25 | 0.0477 | BRD-K52522949 | PC3 | 10.0um | 6.0h |
| 26 | 0.0477 | vorinostat | A549 | 10um | 24h |
| 27 | 0.0469 | vorinostat | PC3 | 10.0um | 24.0h |
| 28 | 0.0469 | trichostatin A | HCC515 | 10.0um | 6.0h |
| 29 | 0.0469 | trichostatin A | A549 | 10.0um | 6.0h |
| 30 | 0.0469 | vorinostat | VCAP | 11.1um | 6.0h |
| 31 | 0.0469 | trichostatin A | HT29 | 10.0um | 6.0h |
| 32 | 0.0469 | BRD-K65814004 | MCF7 | 10.0um | 24.0h |
| 33 | 0.0461 | Scriptaid | HCC515 | 10.0um | 24.0h |
| 34 | 0.0461 | HDAC6 inhibitor ISOX | HCC515 | 10.0um | 24.0h |
| 35 | 0.0461 | HDAC6 inhibitor ISOX | VCAP | 10.0um | 6.0h |
| 36 | 0.0461 | trichostatin A | A375 | 10.0um | 6.0h |
| 37 | 0.0452 | trichostatin A | PC3 | 10.0um | 6.0h |
| 38 | 0.0452 | vorinostat | PC3 | 10.0um | 6.0h |
| 39 | 0.0452 | THM-I-94 | HCC515 | 10.0um | 24.0h |
| 40 | 0.0452 | QUINACRINE HYDROCHLORIDE | A375 | 10.0um | 24.0h |
| 41 | 0.0452 | vorinostat | A375 | 11.1um | 24.0h |
| 42 | 0.0452 | vorinostat | A549 | 11.1um | 6.0h |
| 43 | 0.0452 | vorinostat | PC3 | 10.0um | 6.0h |
| 44 | 0.0452 | BRD-K22503835 | MCF7 | 10.0um | 24.0h |
| 45 | 0.0452 | mitoxantrone | MCF10A | 3.33um | 24h |
| 46 | 0.0444 | vorinostat | HCC515 | 10.0um | 24.0h |
| 47 | 0.0444 | vorinostat | MCF7 | 10.0um | 24.0h |
| 48 | 0.0444 | trichostatin A | A375 | 10.0um | 6.0h |
| 49 | 0.0444 | pracinostat | A549 | 3.33um | 24h |
| 50 | 0.0436 | trichostatin A | HCC515 | 10.0um | 24.0h |

**Supplementary Table 8. List of integrin genes differntially expressed in A549 cells treated with SP for 24 hours identified from RNA-Seq analysis with an adjusted p-value < 0.001 and fold change -2<x>2**

| ENSEMBL ID | Genes | log2FoldChange | pvalue | padj |
|---|---|---|---|---|
| ENSG00000140678 | ITGAX | 2.74 | 9.4E-20 | 1.2E-17 |
| ENSG00000005844 | ITGAL | 2.64 | 9.9E-05 | 9.5E-04 |
| ENSG00000135424 | ITGA7 | 2.22 | 1.0E-17 | 1.0E-15 |
| ENSG00000161638 | ITGA5 | 1.60 | 1.8E-10 | 6.1E-09 |
| ENSG00000115232 | ITGA4 | 1.57 | 1.3E-08 | 3.2E-07 |
